# Ventral hippocampus neurons encode meal-related memory

**DOI:** 10.1101/2023.10.10.561731

**Authors:** Léa Décarie-Spain, Cindy Gu, Logan Tierno Lauer, Keshav S. Subramanian, Samar N. Chehimi, Alicia E. Kao, Iris Deng, Alexander G. Bashaw, Molly E. Klug, Ashyah Hewage Galbokke, Kristen N. Donohue, Mingxin Yang, Guillaume de Lartigue, Kevin P. Myers, Richard C. Crist, Benjamin C. Reiner, Matthew R. Hayes, Scott E. Kanoski

## Abstract

The ability to encode and retrieve meal-related information is critical to efficiently guide energy acquisition and consumption, yet the underlying neural processes remain elusive. Here we reveal that ventral hippocampus (HPCv) neuronal activity dynamically elevates during meal consumption and this response is highly predictive of subsequent performance in a foraging-related spatial memory task. Targeted recombination-mediated ablation of HPCv meal-responsive neurons impairs foraging-related spatial memory without influencing food motivation, anxiety-like behavior, or escape-mediated spatial memory. These HPCv meal-responsive neurons project to the lateral hypothalamic area (LHA) and single-nucleus RNA sequencing and in situ hybridization analyses indicate they are enriched in serotonin 2a receptors (5HT2aR). Either chemogenetic silencing of HPCv-to-LHA projections or intra-HPCv 5HT2aR antagonist yielded foraging-related spatial memory deficits, as well as alterations in caloric intake and the temporal sequence of spontaneous meal consumption. Collective results identify a population of HPCv neurons that dynamically respond to eating to encode meal-related memories.

## INTRODUCTION

Encoding and remembering critical information surrounding food consumption is advantageous to efficiently guide future eating behaviors. Foraging, for example, is facilitated by the retrieval of previously stored spatial information about the location of food sources. Even in the modern environment where food is easily accessible, meal-related memories play an important role in the regulation of eating behaviors. For instance, the ability to recall a recent meal robustly influences subsequent hunger and satiety ratings, as well as the amount of food consumed during the next meal^1–6^.

The hippocampus is a key brain structure for the integration of learning and memory processes with food-seeking and consummatory behaviors^7^. For example, a subnetwork within the human hippocampus implicated in obesity and related eating disorders was recently identified^8^. The ventral subregion of the rodent hippocampus (HPCv) is especially responsive to endocrine and neuropeptide signals that regulate metabolism and food intake control^9–14^, thus making this brain region a prime candidate target for storing meal-related memories. Consistent with this hypothesis, silencing HPCv neurons after food consumption accelerates the onset of a subsequent meal and increases its size^15,16^, indicating that HPCv neuronal activity in close temporal proximity to a meal influences future eating episodes.

Memory engrams are well characterized for fear-related aversive memories where a specific neuronal ensemble in the amygdala encodes fear memory and its reactivation promotes fear-appropriate behavioral responses^17–21^. While engrams encoding aversive events are also formed in the hippocampus^22–26^, engrams encoding positive reinforcing appetitive events such as cocaine administration^27^, female exposure^28,29^, and prosocial interactions^30^ are formed in this region as well. Despite these advances, the neural circuitry encoding memory engrams pertaining to meal consumption remains to be identified. Here we sought to identify a specific ensemble of meal-responsive HPCv neurons and the mechanisms through which they encode meal-related memory engrams.

## RESULTS

### Ventral hippocampus neural activity during meal consumption predicts foraging-related spatial memory performance

To determine whether HPCv neurons are dynamically engaged during meal consumption, we expressed the calcium indicator GCaMP7s in the HPCv (Fig. 1a, b) and assessed bulk calcium-dependent activity over the course of a 30min meal following a 24h fasting period. Results revealed that CA1v bulk calcium-dependent activity dynamically decreases during active eating bouts and increase during inter-bout intervals (Fig. 1c, d). These results suggest that HPCv activity is engaged during a meal at times when the animals are rearing and observing their environment (periods between active eating bouts). These same rats were then trained in a foraging-related spatial memory task (Fig. 1e) and performance index at the memory probe was positively correlated with the magnitude of calcium-dependent activity increases during the interbout intervals of the meal (Fig. 1f). These findings indicate that HPCv activity dynamics during a meal are functionally connected to memory capacity for meal location.

**Figure 1.**
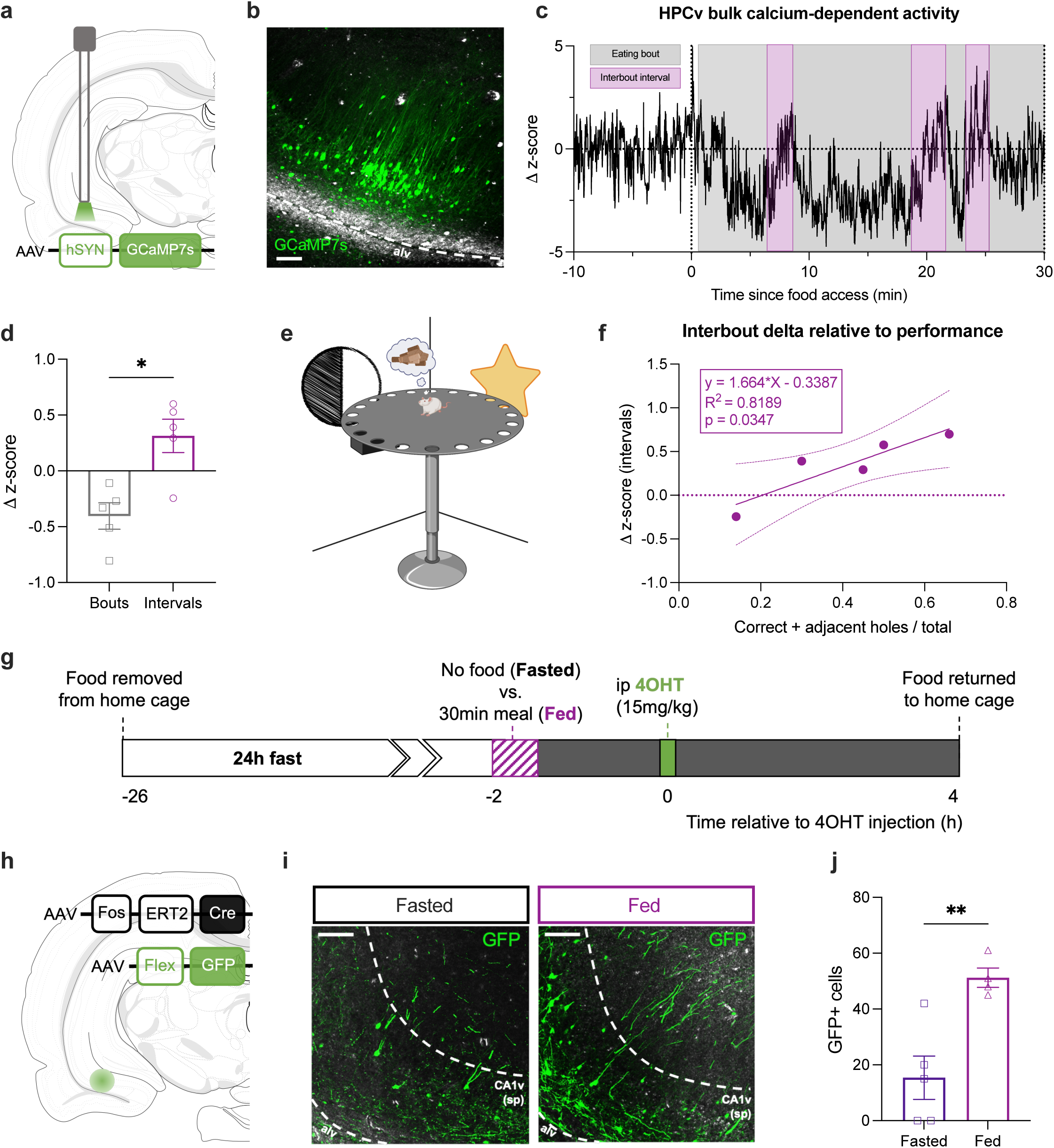
Dynamic changes in ventral hippocampus calcium-dependent activity during a meal are predictive of performance during a foraging-related spatial memory task. (**a**) Diagram of viral approach for fiber photometry. (**b**) Representative photomicrograph of viral expression and optic fiber placement in the ventral CA1 relative to the alveus (alv); scale bar 100 um. (**c**) Representative trace of a single animal of the increase in ventral CA1 calcium-dependent activity during the interbout intervals (purple). (**d**) Average change in z-score for fluorescence over the course of an eating bout versus an interval. (**e**) Foraging-related spatial memory task apparatus. (**f**) Simple linear regression of the increase during interbout intervals predictive of subsequent performance in a separate foraging-related spatial memory task. (**g**) Timeline for intraperitoneal (ip) injection of 4-hydroxitamoxifen (4OHT) under the Fasted or Fed state. (**h**) Diagram of the viral approach for 4OHT-inducible expression of green fluorescent protein (GFP) in ventral CA1 neurons active in a Fasted or Fed state. (**i**) Representative photomicrographs of the pyramidal layer of the ventral CA1 (CA1v(sp)) with GFP^+^ cell bodies from neurons that were active in the Fasted or Fed state, relative to the alveus (alv); scare bar 100 um. (**i**) Quantification of GFP^+^ cell bodies under the Fasted or Fed state in the ventral CA1. Data are presented as mean ± SEM. For Fig. 1d, paired t-test (n=5). For Fig. 1j, unpaired t-test (n=3-5/ group). *p<0.05, **p<0.01.

To identify the subregional anatomical location of HPCv meal-responsive neurons, we employed a ‘targeted recombination in active population’ (TRAP) viral approach to express, following intraperitoneal (ip) administration of 4-hydroxytamoxifen (4OHT; 15mg/kg; H6278, Sigma-Aldrich, St. Louis, USA), green fluorescent protein (GFP) in HPCv field CA1 (CA1v) neurons expressing cFos under a ‘Fasted’ or ‘Fed’ state (Fig. 1g, h). We observed a significantly greater number of GFP^+^ cells bodies under a Fed versus Fasted state, and that GFP^+^ cells in the Fed state were predominantly in the pyramidal layer of CA1v (Fig. 1i, j). These results were further confirmed by cFos immunostaining showing significantly enhanced CA1v cFos expression when rats are perfused under a Fed relative to a Fasted state (Extended Data Fig. 1a, b). In addition, the specificity of the TRAP approach to target CA1v meal-responsive neurons was confirmed by greater overlap of cFos and GFP when feeding conditions at the time of 4OHT administration and perfusion are matched versus mismatched (Extended Data Fig. 1c, d). Altogether, these findings indicate that a subset of HPCv neurons (pyramidal CA1v region) display heightened intracellular activity in response to a meal, that this activity occurs during the periods between eating bouts when animals have an opportunity to observe the meal environment, and that this meal-induced neural response is correlated with foraging-related spatial memory performance.

### Ventral hippocampus meal-responsive neurons selectively promote foraging-related spatial memory

To investigate the function of CA1v meal-responsive neurons, we modified the previous TRAP viral approach to express diphtheria toxin (dTA) in CA1v neurons expressing cFos under a Fasted or Fed state (Fig. 2a), leading to dTA-mediated chronic ablation of CA1v neurons active in response to fasting (Fasted) or meal consumption (Fed). These animals were compared to a group undergoing 4OHT-induced expression of GFP in CA1v neurons (‘Control’). No differences in average daily chow intake nor body weight following 4OHT-induced recombination were observed across experimental groups (Fig. 2b, c). Using a foraging-related spatial memory procedure where animals learn to use visuospatial cues to navigate towards a food source location (Fig. 2d-f), we observed that rats from the Fed group during a memory probe test showed significantly impaired memory performance during a memory probe test for the food location relative to both ‘Control’ and Fasted animals (Fig. 2g and Extended Data Fig. 2a). Animals from the Fed group did perform a greater number of investigations relative to ‘Control’ animals, but no differences in distance traveled were observed (Extended Data Fig. 2b, c).

**Figure 2.**
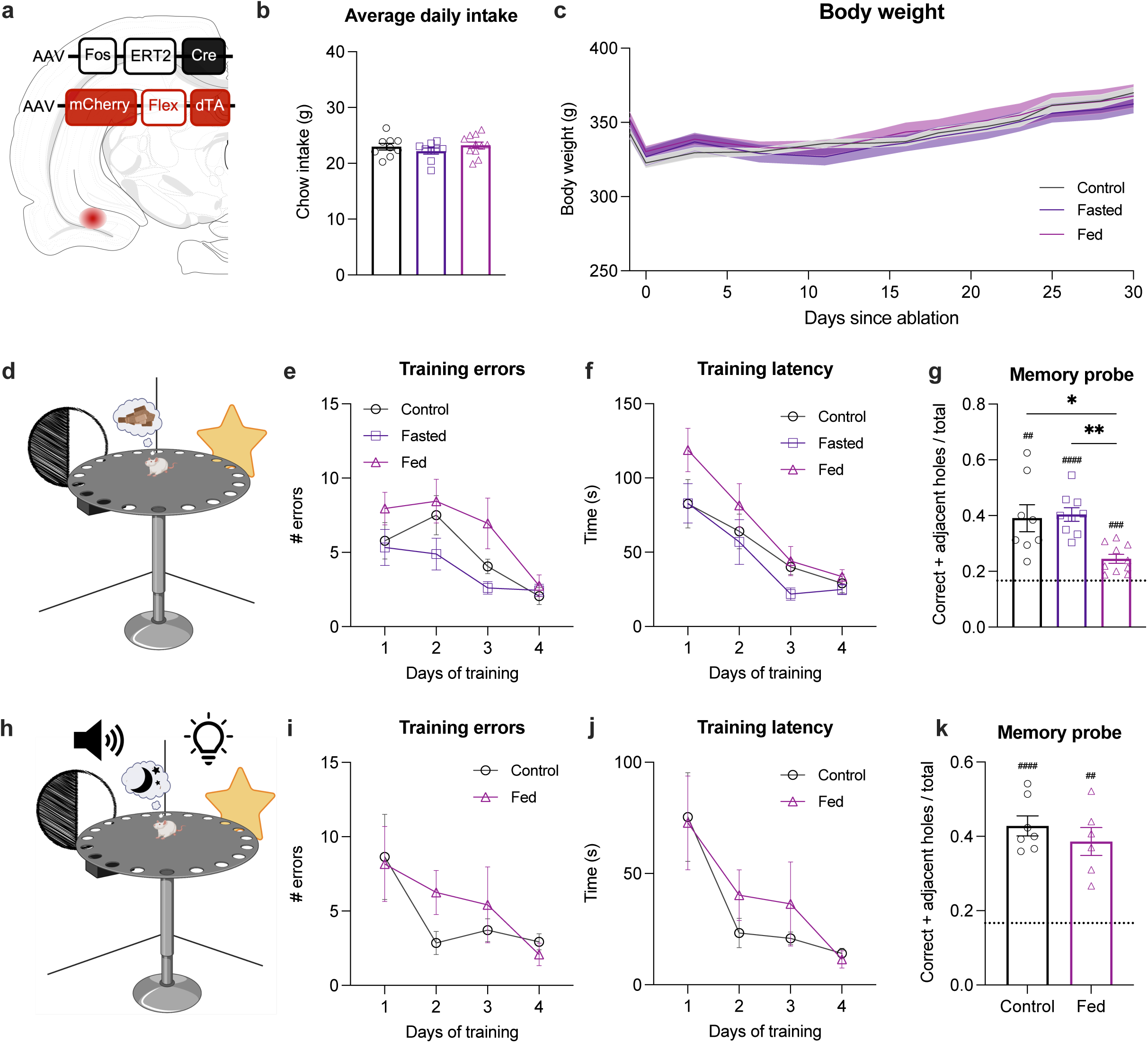
Ablation of ventral hippocampus meal-responsive neurons selectively impairs foraging-related spatial memory. (**a**) Diagram of viral approach for 4-hydroxytamoxifen (4OHT)-inducible Cre-dependent expression of diphteria toxin (dTA) (Fasted and Fed) or green fluorescent protein (Control) in ventral CA1 neurons active in the Fasted or Fed state. (**b**) Average daily food intake following 4OHT intraperitoneal administration to induce dTA-mediated lesion (Fasted and Fed) versus Control. (**c**) Body weight following intraperitoneal administration of 4OHT. (**d**) Diagram of Barnes maze apparatus for the foraging-related spatial memory task. (**e**) Average number of errors and (**f**) latency to find the food during training for the foraging-related spatial memory task. (**g**) Performance index during the probe for the foraging-related spatial memory task. (**h**) Diagram of Barnes maze apparatus for the escape-based spatial memory task. (**i**) Average number of errors and (**j**) latency to find the escape box during training for the escape-based spatial memory task. (**k**) Performance index during the probe for the escape-based spatial memory task. Data are presented as mean ± SEM. For Fig. 2b, one-way ANOVA (n=9-10/group). For Fig. 2c, two-way ANOVA, Tukey post hoc (n=9-11/group). For Fig. 2e-f and Fig. 2i-j, two-way ANOVA (n=9-10/group and n=6-7/group). For Fig. 2g, Kruskal-Wallis, multiple comparisons, (n=9-11/group). For Fig. 2k, two-tailed unpaired t-test, (n=6-7/group); *p<0.05, **p<0.01, ***p<0.005. For Fig. 2g and Fig. 2k, one-sample t-test, different from chance set at 0.1667; ^##^p<0.01, ^###^p<0.005, ^####^p<0.001.

To assess the selectivity of CA1v meal-responsive neurons in mediating meal-related spatial memory (vs. spatial memory in general), another cohort of animals underwent the same meal-related ablation or control treatment as the previous cohort, before training in an escape-based spatial memory task (Fig. 2h) that employs the same apparatus as the foraging version and is of similar difficulty, but without food reinforcement^31^. In this case, Fed rats did not differ from ‘Control’ animals during escape location training (Fig. 2i, j) nor during the memory probe test (Fig. 2k and Extended Data Fig. 2d). The overall number of investigations and distance traveled were similar across groups (Extended Data Fig. 2f, f).

It is possible that the memory impairment in the food-reinforced spatial task in rats with meal-responsive CA1v neurons ablated was secondary to elevated anxiety-like behavior and/or motivation for the sucrose reinforcer, as this brain region is associated with both anxiety^32,33^ and food motivation^34,35^. To test this, a similar cohort of animals was tested in an effort-based operant lever pressing for sucrose rewards (progressive ratio reinforcement schedule), as well as the zero maze test for anxiety-like behavior. Results reveal no group differences in either the progressive ratio test (Extended Fig. 2g) or the time spent in the open areas of the zero maze (Extended data Fig. 2h). Therefore, CA1v meal-responsive neurons appear to selectively modulate spatial memory in the context of foraging without altering spatial memory in general, sucrose motivation, or anxiety-like behavior.

### Ventral hippocampus meal-responsive neurons project to the lateral hypothalamic area

To determine whether HPCv neurons engaged by meal consumption or fasting present distinct projection profiles, whole forebrain GFP^+^ axonal density was assessed in the brains of rats following 4OHT-induced GFP expression in CA1v neurons engaged under a Fasted or Fed state. While GFP^+^ axonal projections were observed in the nucleus accumbens (ACB) and lateral septum (LS) of rats injected with 4OHT under both the Fasted or Fed state (Fig. 3a, b), the presence of GFP^+^ axonal projections in the lateral hypothalamic area (LHA) was exclusive to animals receiving 4OHT under the Fed state (Fig. 3c). In contrast, minimal GFP^+^ axonal density was observed in other known projection targets of CA1v neurons, including the prefrontal cortex (PFC), bed of the stria terminalis (BST), and amygdala (AMY) in either the Fed or Fasted state (Extended Data Fig. 3a-c).

**Figure 3.**
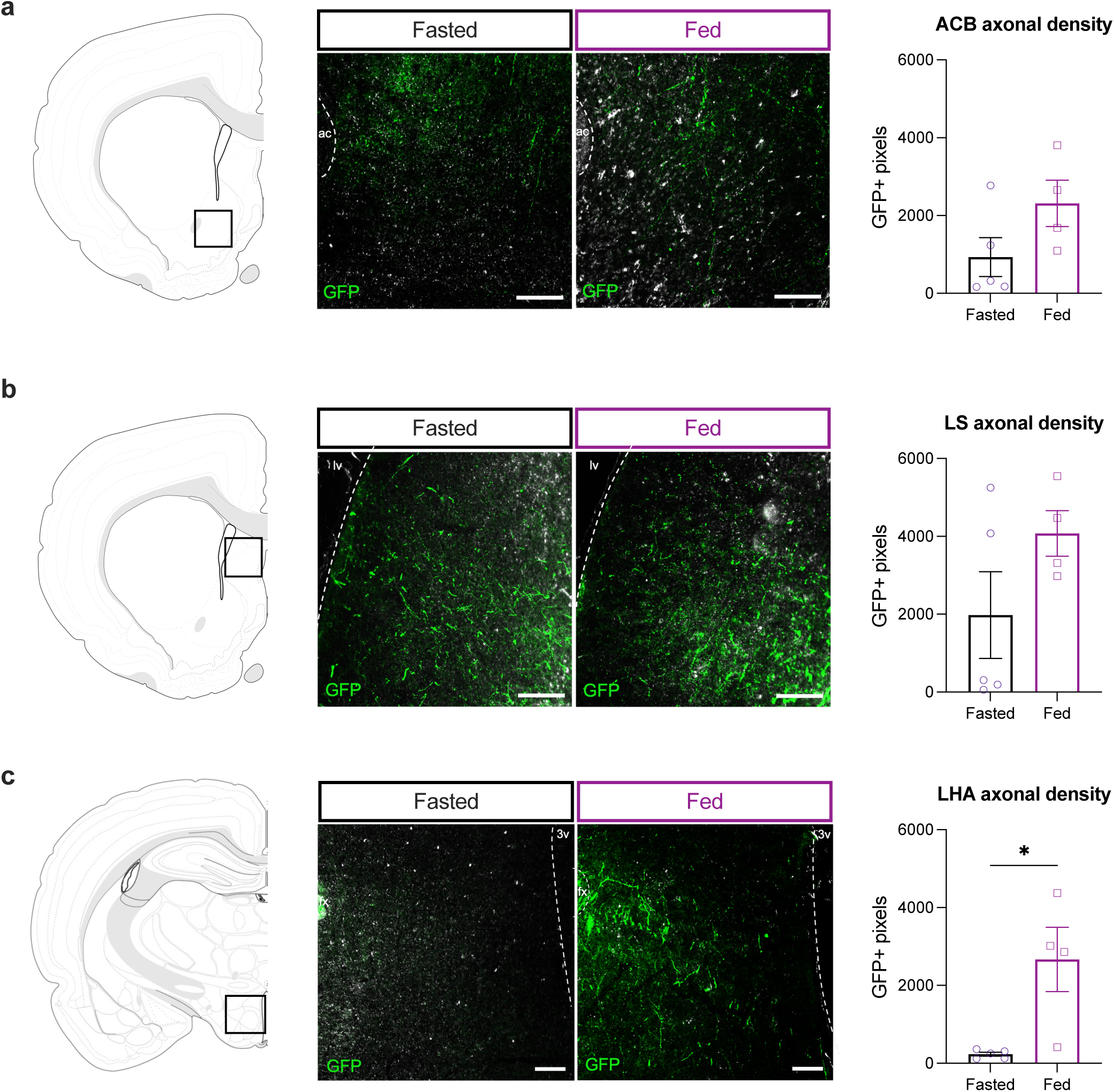
Ventral hippocampus meal-but not fast-responsive neurons project to the lateral hypothalamic area. (**a**) *Left:* Diagram of a coronal section of the nucleus accumbens (ACB). *Middle:* Representative photomicrographs of axonal green fluorescent protein (GFP)^+^ expression in the ACB from ventral CA1 neurons active in the Fasted or Fed state, relative to the anterior commissure (ac); scale bar 100 um. *Right:* Average number of GFP^+^ pixels in the ACB. (**b**) *Left:* Diagram of a coronal section of the lateral septum (LS). *Middle:* Representative photomicrographs of axonal GFP^+^ expression in the LS from ventral CA1 neurons active in the Fasted or Fed state, relative to the lateral ventricle (lv); scale bar 100 um. *Right:* Average number of GFP^+^ pixels in the LS. (**c**) *Left:* Diagram of a coronal section of the lateral hypothalamic area (LHA). *Middle:* Representative photomicrographs of axonal GFP^+^expression in the LHA from ventral CA1 neurons active in the Fasted or Fed state, relative to the fornix (fx) and 3rd ventricle (3v); scale bar 100 um. *Right:* Average number of GFP^+^ pixels in the LHA. Data are presented as mean ± SEM. For Fig. 3a-c, two-tailed unpaired t-test (n=4-5/group); *p<0.05.

To determine the functional relevance of downstream LHA signaling by meal-responsive CA1v neurons in foraging-related memory, we next employed a dual-virus approach to chemogenetically inhibit CA1v neurons sending projections to the LHA following lateral ventricle (LV) infusion of the chemogenetic ligand clozapine-N-oxide (CNO; 18mmol) (Fig. 4a, b). In the absence of LV infusion, rats assigned to the vehicle (VEH) and CNO treatments presented similar number of errors and latencies to reach the food source throughout training (Fig. 4c, d). At the memory probe, rats receiving LV infusion of CNO failed to demonstrate a preference for the hole previously containing food and performed significantly worse than those receiving VEH (Fig. 4e and Extended Data Fig. 4a). LV infusion of CNO did not affect the total number of investigations performed nor the total distance traveled (Extended Data Fig. 4b, c). We have previously shown that LV infusion of CNO in the absence of hM4Di receptors does not impact performance in this task^31^. Thus, CA1v meal-responsive neurons promote foraging-related spatial memory through a CA1v-to-LHA signaling pathway.

**Figure 4.**
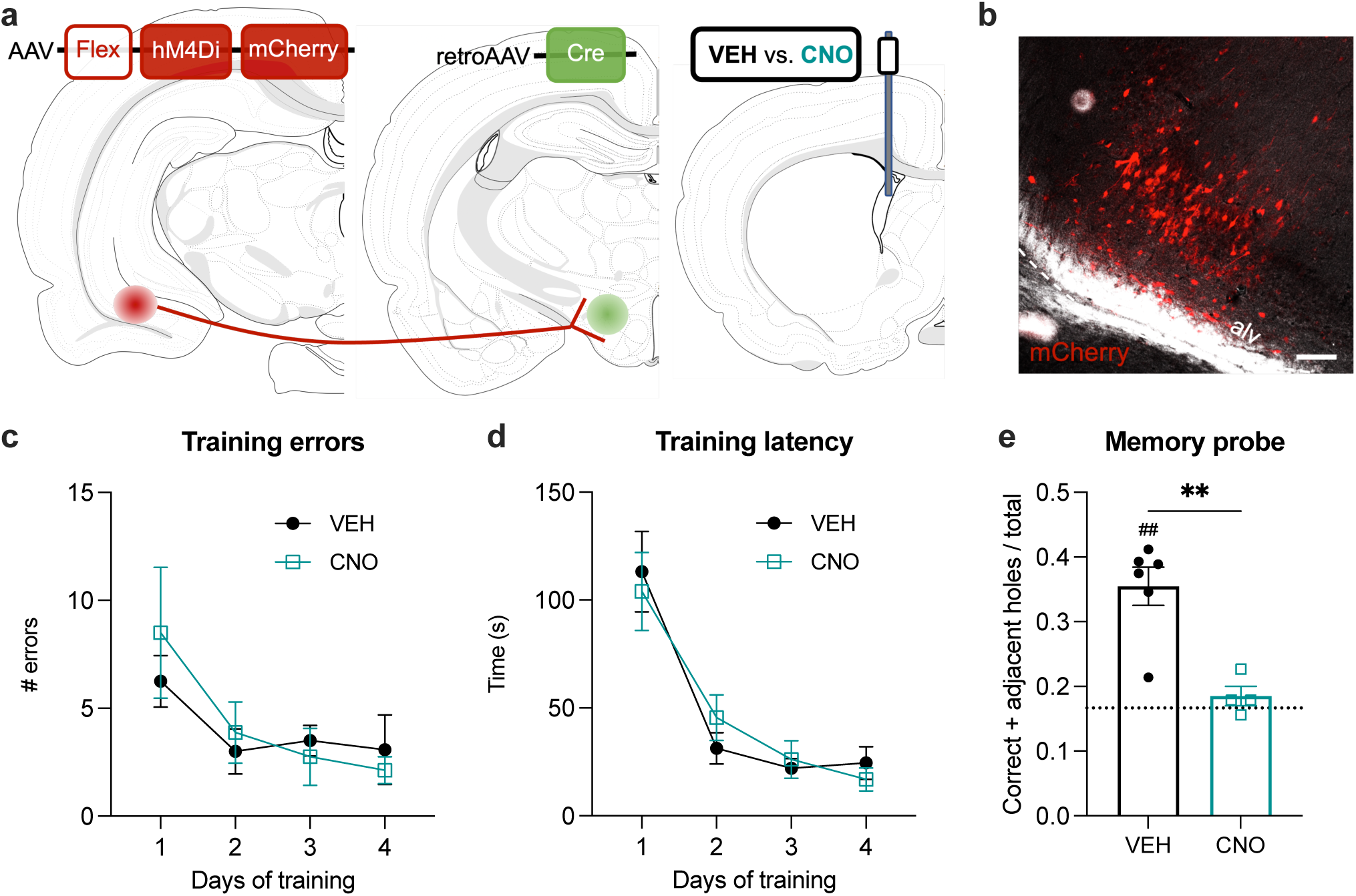
Chemogenetic inhibition of a ventral hippocampus to lateral hypothalamus pathway impairs foraging-related spatial memory. (**a**) Diagram of viral approach for expression of hM4Di receptors in ventral CA1 neurons projecting to the lateral hypothalamic area and administration of vehicle (VEH) or clozapine-N-oxyde (CNO) through a lateral ventricle (LV) cannula. (**b**) Representative photomicrograph of mCherry expression for placement validation of viral injections, relative to the alveus (alv); scale bar 100 um. (**c**) Average number of errors and (**d**) latency to find the food during training for the foraging-related spatial memory task. (**e**) Performance index during the probe for the foraging-related spatial memory task, 1h following LV administration of VEH or CNO. Data are presented as mean ± SEM. For Fig. 5c-d, two-way ANOVA (n=4-6/group). For Fig. 5e, two-tailed unpaired t-test (n=4-6/group); **p<0.01. For Fig. 5e, one-sample t-test, different from chance set at 0.1667; ^##^p<0.01.

### Ventral hippocampus meal-responsive neurons express the serotonin type 2a receptor (5HT2aR)

To identify a molecular signature that distinguishes HPCv meal-responsive neurons from fast-responsive neurons, we performed single nucleus RNA sequencing (snRNAseq) on HPCv microdissections obtained from rats under a Fasted or Fed state. After quality controls, the transcriptome profile of 38,138 nuclei were used to identify 17 cellular subtypes, identified by the expression of prototypical marker genes (Fig. 5a, b). We identified 1,615 cell type-specific differentially expressed genes (Fig. 5c, Supplemental Table 1 and 2), with neuronal transcriptome alterations overrepresented in a small number of cellular subtypes, with endothelial cells (cluster 15; 44 genes downregulated, 301 upregulated) and a population of CA1v excitatory neurons (cluster 5; 224 downregulated, 161 upregulated) undergoing the greatest changes upon meal consumption (Fig 5d, Supplemental Table 1 and 2). Examining marker genes for cFos positive cells in the Fed and Fasted state (Supplemental Table 3), we identified an enrichment of *Htr2a* expression amongst the cFos positive cells in the Fed state (Fig. 5e). This enrichment in *Htr2a* was confirmed via fluorescent in situ hybridization, with greater colocalization of cFos with *Htr2a* observed in the CA1v of rats perfused under a Fed state relative to those perfused under a Fasted state (Fig. 6a, b).

**Figure 5.**
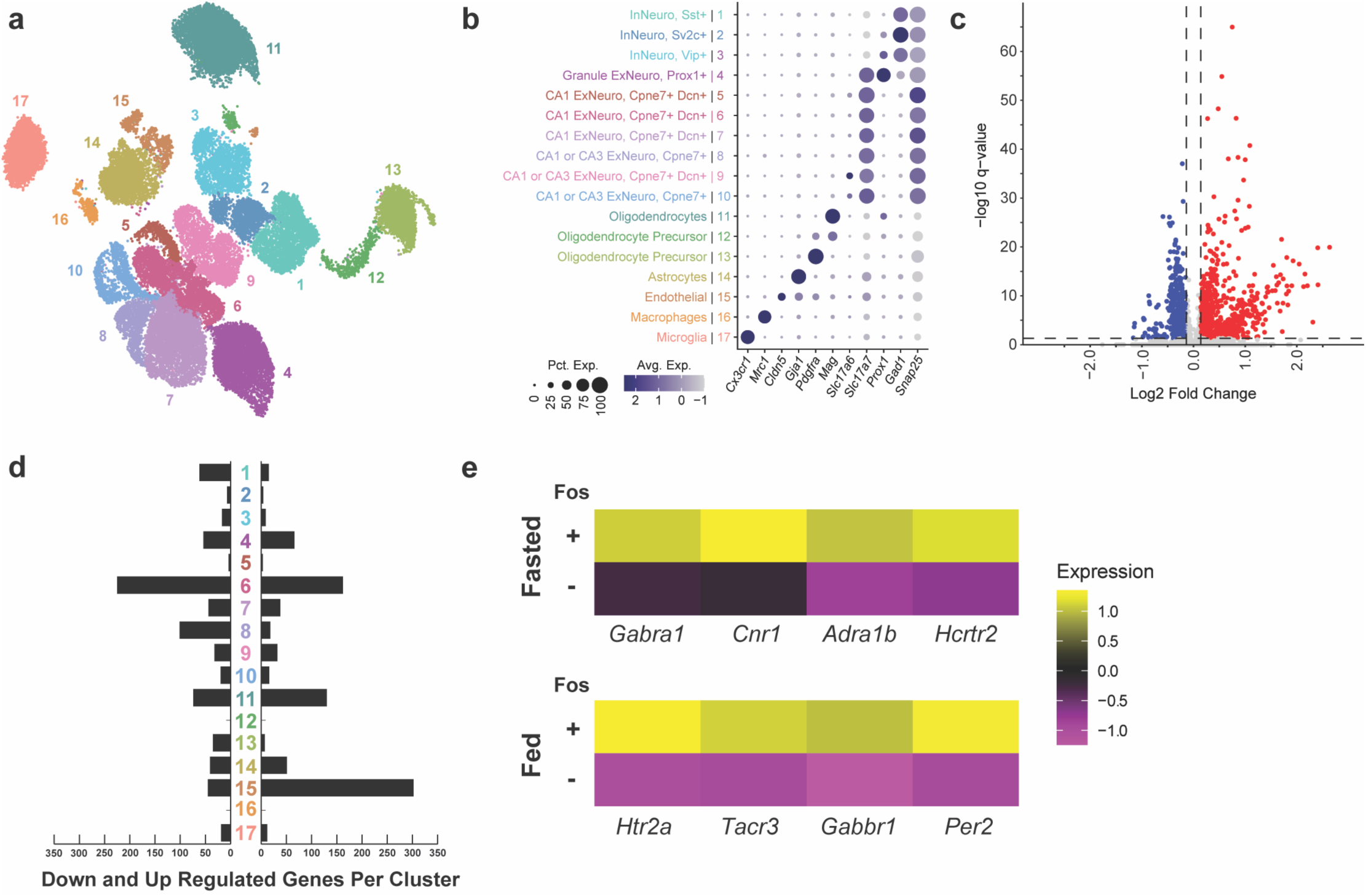
Meal consumption primarily alters the transcriptional profile of ventral hippocampus endothelial cells and CA1v excitatory neurons and engages 5HT2aR expressing cells. (**a**) Uniform manifold approximation and projection (UMAP) of the ventral hippocampus (HPCv) samples identifying 17 clusters. (**b**) Annotation of cellular subtypes with known makers of HPCv cellular subtypes. The size and color of dots are proportional to the percentage of cells expressing the gene (Pct. Exp) and the average expression levels of the gene (Avg. Exp.), respectively. The cluster numbers and colors are matched to that of the UMAP. (**c**) Volcano plot depicting the number of significant differential expression events induced by meal consumption. (**d**) Number of genes with meal consumption-altered increased (right) or decreased (left) expression per cluster. (**e**) Heatmaps of select genes enriched in *Fos*+ cells under a Fasted or Fed state.

**Figure 6.**
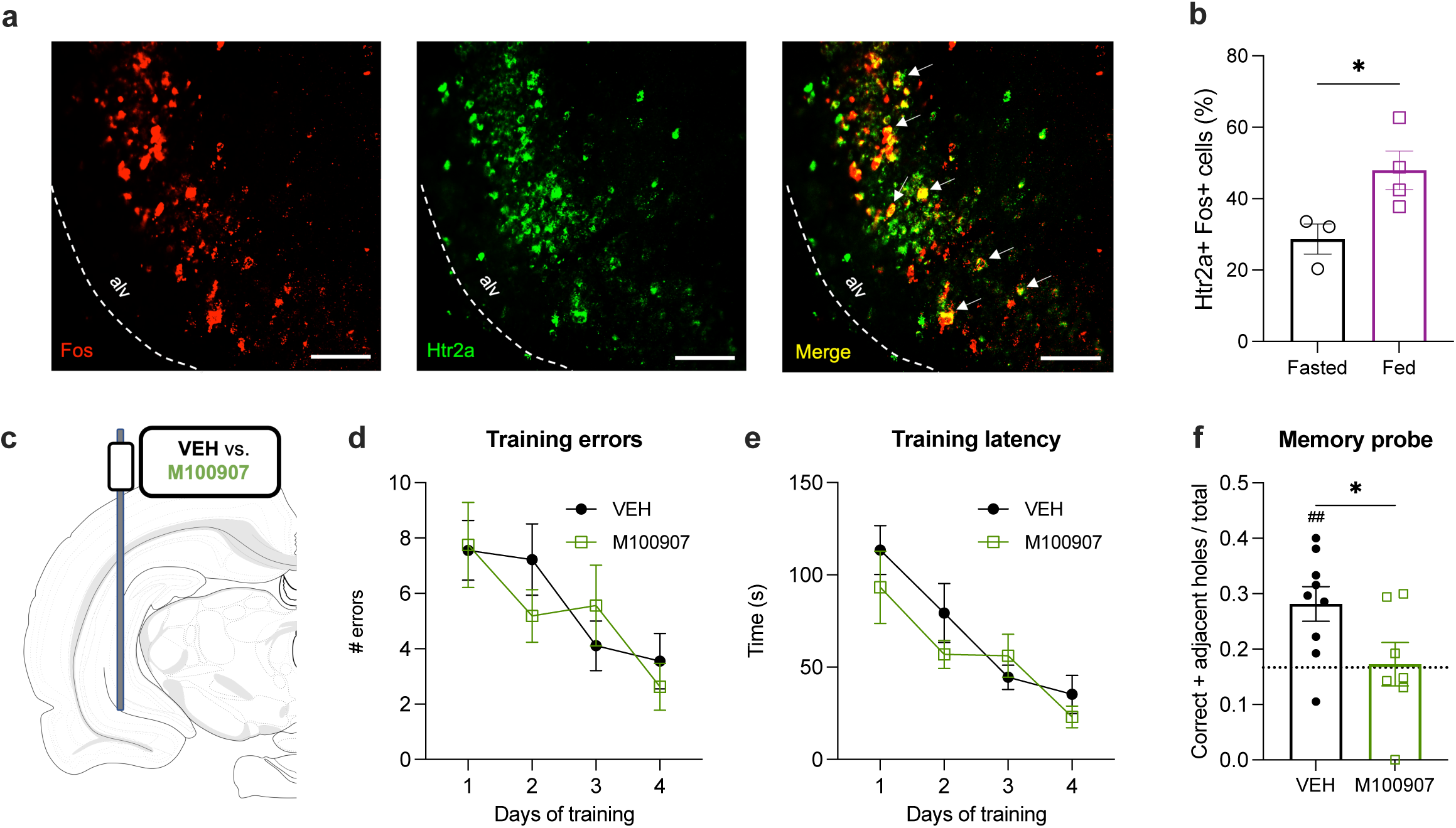
Ventral hippocampus 5HT2aR expressing neurons are engaged by a meal, project to the lateral hypothalamic rea, and are functionally required for foraging-related spatial memory. (**a**) Representative photomicrograph of fluorescent in situ hybridization for *Fos* (red) and *Htr2a* (green) in the ventral CA1, relative to the alveus (alv); scale bar 100um. (**b**) Percentage of ventral CA1 *Fos*+ cells that co-express *Htr2a* in rats perfused under a Fasted or Fed state. (**c**) Diagram of approach for ventral CA1 administration of vehicle (VEH) or the 5HT2aR antagonist M100907 (1g/hemisphere). (**d**) Average number of errors and (**e**) latency to find food during the training for the foraging-related spatial memory task. (**f**) Performance during the probe for the foraging-related spatial memory task, 5min following ventral CA1 infusion of VEH or M100907. Data are presented as mean ± SEM. For Fig. 6b and Fig. 6f, two-tailed unpaired t-test (n=7-9/group and n=5-6/group). For Fig. 6d-e, two-way ANOVA (n=7-9/group); *p<0.05. For Fig. 6f, one-sample t-test, different from chance set at 0.1667; ^##^p<0.01.

To assess the functional relevance of HPCv 5HT2aR signaling in meal-related memory, rats were bilaterally implanted with cannulae targeting the CA1v for infusion of the selective 5HT2aR antagonist M100907 (1μg/hemisphere; M3324, Sigma-Aldrich) or its vehicle (VEH) (Fig. 6c). Rats were trained in the foraging-related spatial memory task and, in the absence of infusions, both groups presented similar number of errors and latencies to reach the food source throughout training (Fig. 6d, e). At the memory probe, rats receiving M100907 failed to demonstrate a preference for the hole previously containing food and performed significantly worse than animals receiving VEH (Fig. 6f and Extended Data Figure 5a), an observation that recapitulates the effect of chemogenetically silencing CA1v to LHA projections. This impairment was not associated with differences in the total number of investigations or distance traveled at the time of the probe (Extended Data Figure 5b, c). These data suggest that HPCv meal-responsive neurons may mediate their effects on foraging-related spatial memory via 5HT2aR signaling.

### Ventral hippocampus meal-responsive neurons regulate the temporal interval between meals

Episodic memory for meal consumption not only includes information about spatial and contextual cues that inform about the location of the meal, but also temporal information regarding when the meal was consumed. The latter powerfully influences meal timing and amount consumed in both humans and rodent models in a manner indicating that memory for time since last eating is factored into the next meal ^1,5,6,16,36,37^. To determine whether acute manipulation of signaling pathways from CA1v meal-responsive neurons influences food intake and meal timing, meal pattern analyses were conducted following either CA1v 5HT2aR antagonism or CA1v-to-LHA chemogenetic silencing prior to dark onset. CA1v infusion of the 5HT2aR antagonist M100907 increased food intake relative to VEH at the 2h timepoint (Fig. 7a), and while no statistically significant differences were detected in either meal size (Fig. 7b) or meal frequency (Fig. 7c), inter-meal intervals were decreased by CA1v 5HT2aR antagonism (Fig. 7d). Similarly, LV infusion of CNO to disconnect CA1v-to-LHA signaling increased food intake relative to VEH treatment over a period of 6h (Fig. 7e) and, while no changes were observed in meal size (Fig. 7f), this effect was driven by an increase in meal frequency (Fig. 7g) and a decrease in inter-meal intervals (Fig. 7h). Together, these results highlight a role for HPCv meal-responsive neurons to encode both spatial and temporal aspects of meal-related memories, and that these processes influence spontaneous eating patterns.

**Figure 7.**
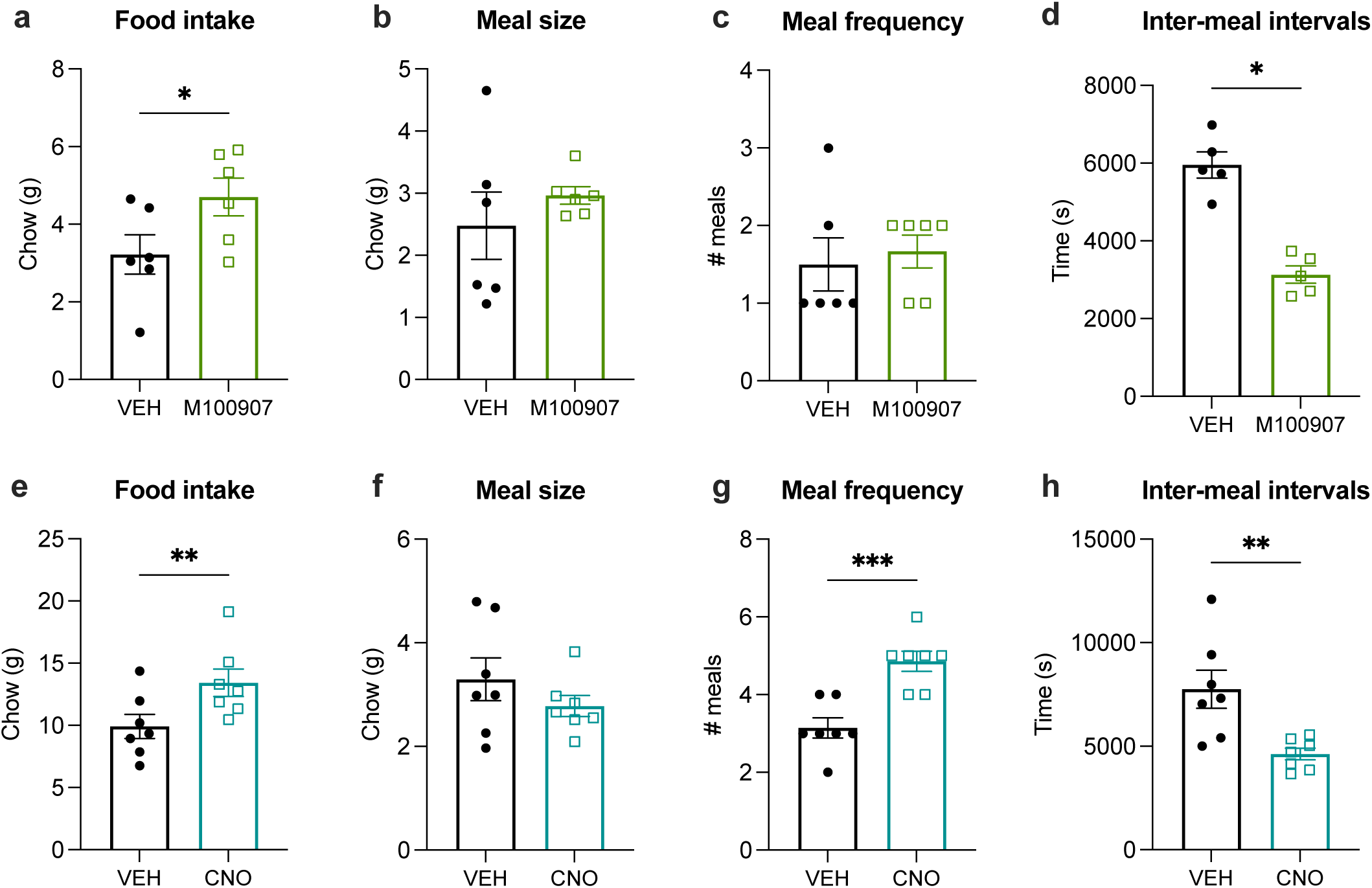
Blockade of ventral hippocampus to lateral hypothalamic area signaling or 5HT2aR signaling increases food intake by reducing temporal intervals between meals during spontaneous feeding. (**a**) Average 2h chow intake, (**b**) meal size, (**c**) meal frequency and (**d**) inter-meal intervals following ventral CA1 administration of vehicle (VEH) or M100907. (**e**) Average 2h chow intake, (**f**) meal size, (**g**) meal frequency and (**h**) inter-meal intervals following lateral ventricle administration of vehicle (VEH) or clozapine-N-oxide (CNO) in rats expressing hM4Di receptors in ventral CA1 neurons projecting to the lateral hypothalamic area. Data are presented as mean ± SEM. Two-tailed paired t-test (n=7/group and n=5-6/group); *p<0.05, **p<0.01, ***p<0.001.

## DISCUSSION

Memory function greatly contributes to the control of eating behavior, yet the neural substrates governing this phenomenon have been minimally investigated. Here, we show that activation of a population of hippocampal neurons (CA1v pyramidal) during meal consumption serves to encode meal-related spatial and temporal mnemonic information. We functionally characterize these meal-responsive neurons and demonstrate that they project to the lateral hypothalamic area and express the 5HT2a receptor. Remarkably, this neuronal ensemble selectively mediates meal-related memory encoding without influencing food motivation or other memory processes. Together, these novel findings provide evidence for the formation of meal engrams in the CA1v that serve to facilitate foraging behavior, and when food is readily available, the temporal parameters of meal consumption.

That HPCv neurons are engaged upon food consumption is consistent with previous studies revealing that optogenetic inhibition of HPCv neurons influences food intake^16,34,37^. Further, fiber photometry recordings of HPCv calcium-dependent activity reveal dynamic changes during foraging and food-seeking behaviors^31,33^. Our present results advance these findings by establishing a role for HPCv neuronal activity during eating behaviors in encoding meal-related memory, thus providing a mechanistic framework for this previous literature. Further, HPCv calcium activity dynamics during meal consumption reveal elevations selectively during the intervals between active eating bouts. This suggests that HPCv neurons more effectively encode meal-associated memories during periods when the animals (and presumably humans) can more easily observe exteroceptive features of the eating environment.

The CA1v regulates eating behaviors through diverse neuronal pathways, including axonal projections targeting the LS^33,34^, ACB^13,38^ and PFC^39^. Our findings reveal that CA1v meal-responsive neurons distinctively send axonal projections to the LHA, which is in line with the well-known contribution of the LHA in regulating eating behaviors^40^. In addition, previous work identified a role for CA1v-to-LHA neuronal projections in modulating appetite and meal parameters^11,14^ through ghrelin signaling. Therefore, it may be the case that periprandial concentrations of ghrelin recruit CA1v meal-responsive neurons to encode mnemonic information. Consistent with this framework, hippocampus ghrelin signaling promotes hippocampal neuronal plasticity, as well as conditioned cue-potentiated overeating^10,41^.

The predominance of the literature on serotonin signaling and food intake control focuses on the anorexigenic effects of the serotonin type 2c receptor (5HT2cR), notably in the hindbrain^42–44^. Here, we show a role for HPCv 5HT2aR in modulating food intake, which is consistent with reports of *Htr2a* variants influencing food intake and preferences in humans^45,46^. In rodents, 5HT2aR-expressing neurons in the central amygdala are responsive to fasting and modulate food consumption^47,48^. That HPCv 5HT2aR influence food intake through memory function is in line with reports in humans with *Htr2a* allele variants influencing hippocampal function^49–51^ and work in rodents illustrating a role for 5HT2aR signaling in learning in memory processes^52–57^, and specifically in the HPCv^58^. Serotonin release within the HPCv may result from serotoninergic projections originating from the dorsal raphe nucleus (DRN) as engram-encoding positive experiences within the dorsal hippocampus dentate gyrus are preferentially reactivated by DRN neurons^59^. However, the median raphe nucleus (MRN) also sends serotoninergic projections to the HPCv that are involved in goal-directed behavior^35^ and fear memory^60^. In addition, the interpeduncular nucleus contains serotonin neurons that innervate the HPCv to influence stress and reward functions^61,62^. While the source of serotonin remains to be determined, our work highlights a novel mechanism through which 5HT2aR signaling in HPCv meal-responsive neurons influences eating behaviors by encoding meal-related memory.

Memories relating to the timing of a meal and the amount of food consumed potently influence subsequent hunger, satiety, and caloric consumption^1–4^. These phenomena are strikingly illustrated in patients suffering from medial temporal lobe damage who are unable to form new episodic memories and will thus, if permitted, consume multiple consecutive meals in a short temporal window, resulting in greater overall intake^63^. Additionally, human obesity and disordered eating behaviors are associated to alterations within an orexigenic hippocampal subnetwork^8^. Our present findings illuminate the neural circuitry mediating these effects as both chemogenetic inhibition of a CA1v-to-LHA signaling pathway and HPCv 5HT2aR antagonism disrupted meal-related memory, increased caloric intake, and shortened the inter-meal temporal window, an effect also consistent with rodent studies demonstrating effects of HPCv neuronal inhibition on subsequent meal patterns^16,36,37^.

Present results should be interpreted in the context of a few limitations. For example, the TRAP-mediated ablation experiment employs a chronic loss of function approach that may lead to compensatory neural adaptations secondary to the loss of CA1v responsive neurons.

Therefore, different outcomes might be observed with an activity-based acute and reversible inhibitory approach. Additionally, despite being driven by activity-based neuroanatomical tracing and RNA sequencing experiments, our CA1v-to-LHA chemogenetic and 5HT2aR pharmacology approaches are not restricted to HPCv meal-responsive neurons. Nonetheless, these findings provide a novel neural systems mechanism through which learning and memory processes synergize with eating behaviors.

Results from our snRNAseq of meal-responsive CA1v neurons open the door to promising new research avenues. For example, future work should explore the function of differentially expressed genes in discrete neural populations of the HPCv in response to meal consumption. Results (sequencing dataset now publicly available) reveal that the largest number of differentially expressed genes are present in a cell cluster delineated by the presence of endothelial cells, which comprise the blood-brain barrier. Whether the blood-brain-barrier undergoes permeability changes to facilitate HPCv function upon meal consumption is an exciting area to investigate. Given that the majority of differentially expressed genes in this cluster were upregulated in the Fed vs. Fasted state, it may be that blood-brain barrier permeability is increased during and immediately after a meal to preclude blood-to-brain entry of potential toxins introduced from the food consumed. Another question arises with regards to the status of these HPCv meal-responsive neurons in the context of obesity and whether they can be targeted to reduce food intake in a diet-induced obese model. Answering these questions would significantly advance our understanding of the neurobiology underlying the higher-order control of eating behavior. Overall, this work identifies a novel population of HPCv neurons that is enriched in 5HT2aR and projects to the LHA to encode the engram for meals.

## METHODS

### Animals

Adult male Sprague–Dawley rats (Envigo; 250-275g on arrival) were individually housed with *ad libitum* access to water and chow (LabDiet 5001, LabDiet, St. Louis, MO) on a 12h:12h reverse light/dark cycle. All procedures were approved by the University of Southern California Institute of Animal Care and Use Committee.

### Intracranial injections

Rats were anesthetized via an intramuscular injection of an anesthesia cocktail (ketamine 90mg/kg body weight [BW], xylazine, 2.8mg/kg BW and acepromazine and 0.72mg/kg BW) followed by a pre-operative, subcutaneous injection of analgesic (buprenorphine SR, 0.65mg/kg BW). Following shaving and preparation of the incision site with iodine and ethanol swabs, rats were placed in a stereotaxic apparatus for viral infusions using a microinfusion pump (Harvard Apparatus, Cambridge, USA) connected to a 33-gauge microsyringe injector attached to a PE20 catheter and Hamilton syringe at a flow rate of 83.3nL/sec. Injectors were left in place for 2min post-injection, prior to placement of an indwelling cannula or closing of the incision site using skin adhesive.

### Fiber photometry

To record bulk calcium-dependent activity in the ventral CA1 (CA1v), rats received bilateral infusion (300nL/side) of a synapsin-driven GCaMP7s-expressing virus (AAV9-hSyn-GCaMP7s-WPRE; Addgene, Watertown, USA) at the following coordinates: -4.9mm AP (defined at bregma), +/-4.8mm ML (defined at bregma), and -7.8mm (defined at skull surface at site). A fiber-optic cannula (Doric Lenses Inc, Quebec, Canada) was implanted at the injection site and affixed to the skull with jeweler’s screws, instant adhesive glue, and dental cement.

Photometry recording sessions were conducted as described previously^31,64^. Briefly, 24h-fasted animals were placed in a familiar arena and were given access to chow pellets for 30min, following a 15min recording of baseline calcium-dependent activity. Photometry signal was acquired using the fiber photometry system (Neurophotometrics, San Diego, USA) at a sampling frequency of 40Hz and administering alternating wavelengths of 470nm (calcium-dependent) or 415nm (calcium-independent). Active eating bouts were timed-stamped in real-time by experimenters observing the animals using the data acquisition software Bonsai.

Photometry signal was corrected by subtracting the calcium-independent from the calcium-dependent signal fitting the result to a biexponential curve. This corrected fluorescence signal was normalized by calculating, for each rat, the ΔF/F using the average fluorescence signal for the entire recording and further converting it to z-scores.

### Targeted recombination in active population (TRAP)

Rats received bilateral CA1v injection of a 1:1 cocktail of viruses (400nL/side): [1] a tamoxifen-inducible virus expressing Cre recombinase under the control of a cFos promoter (AAV8-cFos-ERT2-Cre-ERT2-PEST; Stanford Vector Core, Stanford, USA) and [2] a Cre-dependent virus expressing green fluorescent protein (AAV-Flex-GFP; Addgene) or diphteria toxin (AAV2/9-EF1a-mCherry-Flex-dTA; Neurophotonics, Quebec, Canada). Three weeks following surgery, 24h fasted rats are offered 30min access to chow pellets (Fed) or not (Fasted), 2h prior to receiving an intraperitoneal (ip) injection of 4-hydroxytamoxifen (4OHT; 15mg/kg; H6278, Sigma-Aldrich, St. Louis, USA) dissolved in saline with 2% Tween-80 and 5% DMSO, similarly to^65^. Behavioral measures and histological analyses commenced at least 1 week following 4OHT-induced recombination.

For the histological analyses of GFP+ cell bodies and axonal projections, coronal sections were compared between rats undergoing 4OHT-driven expression of GFP (AAV8-cFos-ERT2-Cre-ERT2-PEST mixed with AAV-Flex-GFP) in the Fed versus Fasted state. For the chronic ablation experiment, rats undergoing dTA-mediated ablation (AAV-8-cFos-ERT2-Cre-ERT2-PEST mixed with AAV2/9-EF1a-mCherry-Flex-dTA) induced by 4OHT administration following meal consumption (Fed) or not (Fasted) were compared to animals with CA1v GFP expression (AAV-8-cFos-ERT2-Cre-ERT2-PEST mixed with AAV-Flex-GFP) induced by 4OHT administration (‘Control’).

### Foraging-related spatial memory task

The foraging-related spatial memory task was conducted as described previously^31^. Briefly, using an elevated circular platform with 18 equally distanced holes surrounding by visuospatial cues on the walls, rats were trained, over the course of 4 days (3min/trial, 2 trial/day), to navigate towards the escape hole containing 5 sucrose pellets (F0023, Bio-Serv Flemington, USA). Distinct spatial cues are present on the walls surrounding the table and low-level ambient lighting is provided by floor lamps. Memory probe occurred 72h after the last training session and consisted of a 2-min trial during which preference of the rat for the previously reinforced hole was assessed in the absence of sucrose pellets. A performance index was calculated by dividing the number of investigations for the previously reinforced hole and its adjacent holes by the total number of investigations. Investigations were quantified by tracking animal’s head using the AnyMaze Behavior Tracking Software (Stoelting, Wood Dale, USA).

### Escape-based spatial memory task

The escape-based spatial memory task was conducted as described previously^31^, using a protocol that is identical to the foraging-related spatial memory task, with the exception that sucrose pellets were absent, and a mildly aversive overhead bright light (120W) and white noise (75dB) were presented until the rat found the escape box.

### Chemogenetic silencing of the HPCv-lateral hypothalamic area (LHA) pathway

Chemogenetic silencing of CA1v projections to the LHA was performed using a previously published dual-viral approach^31^. Rats received bilateral HPCv injection (300nL/side) of a Cre-dependent hM4Di-expressing virus (AAV2-Flex-hM4Di-mCherry; Addgene) using the same stereotaxic coordinates as the TRAP approach, as well as bilateral LHA injection (200nL/side) of a retrograde AAV expressing Cre recombinase (AAV2[retro]-hSYN1-EGFP-2A-iCre-WPRE; Vector BioLabs) at the following stereotaxic coordinates: -2.9mm AP, +/- 1.1mm ML, and -8.6mm DV (all defined at bregma). Animals were also implanted with a unilateral indwelling cannula (26-gauge, Plastics One, Roanoke, USA), affixed to the skull with jeweler’s screws, instant adhesive glue and dental cement, targeting the lateral ventricle (LV) at the following stereotaxic coordinates: -0.9mm AP (defined at bregma), +1.8mm ML (defined at bregma), and -2.6mm DV (defined at skull surface at site).

Placement of the LV cannula was confirmed by elevation of at least 100% of baseline glycemia following infusion of 210μg (2uL) of 5-thio-D-glucose (5TG) through the cannula using an injector extending 2mm beyond the end of the cannula guide (Ritter et al, 1981). In animals that failed to present the glycemic response, the test was repeated with a 2.5mm injector, and upon passing the 5TG test, 2.5mm injectors were used for the remainder of the study.

For assessment in the foraging-related spatial memory task and meal pattern analysis, measures were performed 1h following LV infusion (2uL) of artificial cerebrospinal fluid (Veh) or clozapine-N-oxide (CNO; 18mmol).

### Single nucleus RNA sequencing

Following a 24h fast, rats were offered a 30min access to chow pellets (Fed) or not (Fast), 90min prior to sacrifice. Fresh brains were harvested, and flash frozen in isopentane surrounded by dry ice. Coronal sections (250um) were made on a cryostat and bilateral microdissections of the HPCv were obtained using a tissue puncher (2mm diameter). Nuclei suspensions were generated from the frozen samples and data, similar to our prior descriptions^66–68^. Nuclei were processed for the 10x Genomics 3’ gene expression assay (v3.1) per manufacturer protocols, and 10x Genomics Cell Ranger was used to align sequencing reads. Filtered read count matrixes were merged via Seurat, and nuclei with high UMI, low gene count and >5% mitochondrial reads were removed as doublets or low quality as we previously described^68^. Counts were normalized to 10,000 reads per nucleus and scaled, and variable genes were identified mean.var.plot method in the FindVariableFeatures function in Seurat. The first 50 principal components were used to create a neighbors graph and the data was then clustered at a resolution of 0.8. Subsequent quality control steps removed a cluster with abnormally low UMI, a cluster with mixed cell type markers, and three clusters with most nuclei contributed by only 1-2 samples. The remaining data was normalized and scaled again, clustered at a resolution of 0.15, resulting in identification of 17 clusters across 38,138 nuclei. Clusters were identified by expression of major cell type markers: microglia – *Cx3cr1*; macrophages – *Mrc1*; endothelial cells – *Cldn5*; astrocytes – *Gja1*; oligodendrocyte precursor cells – *Pdgfra*; oligodendrocytes – *Mag*; neurons – *Snap25*; excitatory neurons – Slc17a6, *Slc17a7*; granule cells – *Prox1*; inhibitory neurons – *Gad1*. Differential gene expression analysis between the two experimental groups was performed within each cluster using a Wilcoxon rank sum test and p-values were adjusted for multiple testing using a Bonferroni correction for the total number of genes in the data set.

### Pharmacological manipulations of ventral hippocampus serotonin receptor type 2a

Rats were bilaterally implanted with indwelling cannulae in the CA1v, at the following stereotaxic coordinates: -4.9mm AP (defined at bregma), +/-4.8mm ML (defined at bregma), and -5.8mm (defined at skull surface at site). Cannulae were fixed to the skull using jeweler’s screws, instant adhesive glue and dental cement.

For assessment in the foraging-related spatial memory task and meal pattern analysis, measures were performed 5min following CA1v infusion (200nL/side) of 33% DMSO in artificial cerebrospinal fluid (Veh) or the serotonin type 2a receptor antagonist M100907 (1μg/side; M3324, Sigma-Aldrich).

Cannula placement was confirmed post-mortem via a 200nL infusion of 2% Chicago sky blue ink through the guide cannula. Data from animals with dye confined to the CA1v were included in the analyses.

### Meal pattern analysis

The Biodaq automated food intake monitoring cage system (Research Diets, New Brunswick, USA) was used to assess 24h food intake, meal size, meal frequency and post-meal intervals. Two 24h measures spaced a week apart were conducted for meal pattern analysis in the CA1v-to-LHA chemogenetic silencing (VEH vs CNO) and CA1v 5HT2aR antagonism (VEH vs M100907) experiments. Meal parameters were set at a minimum meal size of 0.2g and inter-meal interval of 900s^69^.

### Immunohistochemistry

Rats received an intramuscular injection of an anesthesia cocktail (ketamine 90mg/kg BW xylazine, 2.8mg/kg BW and acepromazine and 0.72mg/kg BW) prior to transcardiac perfusion with 0.9% sterile saline (pH 7.4) followed by 4% paraformaldehyde (PFA) in 0.1M borate buffer (pH 9.5; PFA). Harvested brains were post-fixed in PFA with 12% sucrose for 24h, then flash frozen in isopentane cooled in dry ice. Coronal sections (30um) were obtained using a microtome, collected in 5-series and stored in antifreeze solution at −20°C until further processing.

To amplify the native GFP signal in axonal projections from animals undergoing 4-OHT-induced expression of GFP under the Fed or Fasted state, the chicken anti-GFP primary antibody (1:500; Ab13970, Abcam, Boston USA) was used followed by a donkey anti-chicken secondary antibody conjugated to AF488 (1:500; AB_2340375, Jackson Immunoresearch, West Grove, USA).

To amplify the native mCherry signal from animals undergoing CA1v-to-LHA chemogenetic silencing, the rabbit anti-RFP primary antibody (1:2000; AB_2209751, Rockland, Limerick, USA) was used followed by a donkey anti-rabbit secondary antibody conjugated to Cy3 (1:500; AB_2307443, Jackson Immunoresearch).

To measure cFos protein expression in the CA1v under the Fed and Fasted state, the mouse anti-c-Fos primary antibody (1:1000; Ab208942, Abcam) was used followed by a donkey anti-mouse secondary antibody conjugated to AF647 (1:500; Jackson Immunoresearch).

Antibodies were prepared in 0.02M potassium phosphate buffered saline (KPBS) solution containing 0.2% bovine serum albumin and 0.3% Triton X-100 at 4°C overnight. After thorough washing with 0.02M KPBS, sections were incubated at 4°C overnight in secondary antibody solution. Sections were mounted and coverslipped using 50% glycerol in 0.02 M KPBS and the edges were sealed with clear nail polish. Photomicrographs were acquired using a Nikon 80i (Nikon DSQI1,1280X1024 resolution, 1.45 megapixel) under epifluorescence or darkfield illumination, as described previously (Cell Rep).

### Fluorescent in situ hybridization

Fluorescent in situ hybridization analyses for cFos (ACD, 403591, Newark, USA) and Htr2a (ACD, 424551) were performed in HPCv coronal sections from rats perfused under a Fed or Fasted state. Co-localization of cFos positive cells with Htr2a in the CA1v was quantitively analyzed.

### Effort-based lever pressing

Using operant conditioning boxes (Med Associates Inc, St Albans, USA), rats were trained to lever press for 45mg sucrose pellets (Bio-Serv, F0023) over the course of 6 days with 1 session each day (2 days of fixed ratio 1 with autoshaping procedure, 2 days of fixed ratio 1 and 2 days of fixed ratio 3 reinforcement schedule). For the test session, rats were placed in the chambers to lever press for sucrose under a progressive ratio reinforcement schedule. The response requirement increased progressively using the following formula: F(i) = 5ê0.2i-5, where F(i) is the number of lever presses required for the next pellet at i = pellet number and the breakpoint was defined as the final completed lever press requirement that preceded a 20-min period without earning a reinforcer, as described previously^12,31^.

### Zero-maze

Rats were placed on an elevated circular track consisting of 2 open zones (3cm high curbs) and 2 closed zones (17.5cm high walls) and left to explore for 5min during the dark cycle. The total time spent in open zones (defined as body center in open sections) was measured using the AnyMaze Behavior Tracking Software (Stoelting).

### Statistical analysis

Data are expressed as mean +/- SEM. Differences were considered statistically significant at p<0.05. All variables were analyzed, and all graphs were created using the GraphPad Prism 9 software.

Two-tailed paired t-test as used to compare changes in CA1v calcium activity dynamics during eating bouts versus interbout intervals, while two-tailed unpaired t-tests were used to compare the number of GFP+ and cFos+ cells, as well as number of Htr2a+ and cFos+ cells, under the Fed and Fasted states. A simple linear regression analysis was conducted to investigate the relationship between CA1v increases in calcium-dependent activity during interbout intervals and performance in the foraging-related spatial memory task. A two-way ANOVA (group x condition) was used to analyze overlap of GFP with cFos in animals injected with 4OHT under a Fed or Fasted state and perfused under a matched or mismatched condition. Repeated measures two-way ANOVAs (group x time) were employed to assess body weight and acquisition in both the foraging-related and escape-based spatial memory tasks (errors and latencies) in the TRAP ablation (4OHT-induced dTA expression) experiment. A repeated measure two-way ANOVA (group x time) was also used to assess acquisition in the foraging-related memory task in the chemogenetic silencing of the CA1v-to-LHA pathway and CA1v 5HT2aR antagonism experiments. A Kruskal-Wallis test with Dunn’s posthoc comparison was employed to assess performance in the foraging-related spatial memory task in the TRAP ablation experiment. One-way ANOVAs were used to compare performance in the effort-based lever pressing and zero-maze tasks in the TRAP ablation cohort. Two-tailed unpaired t-test were employed to assess performance in the escape-based spatial memory task for the TRAP ablation cohort and foraging-related spatial memory task in the CA1v-to-LHA chemogenetic and CA1v 5HT2aR antagonism experiments. Two-tailed paired t-tests were employed to evaluate meal patterns in the CA1v-to-LHA chemogenetic and the CA1v 5HT2aR antagonism experiments.

Significant ANOVAs were analyzed with a Tukey posthoc test, where appropriate. Outliers were identified as being more extreme than the median +/- 1.5 * interquartile range. For all experiments, assumptions of normality, homogeneity of variance (HOV), and independence were met where required.

## ACKNOWLEDGEMENTS

This work was supported by a Quebec Research Funds Postdoctoral Fellowship (315201) and an Alzheimer’s Association Research Fellowship to promote diversity (AARFD-22-972811) to LDS, a National Science Foundation Graduate Research Fellowship to KSS and National Institute of Diabetes and Digestive and Kidney Diseases grants: DK105155 to MRH and DK104897 to SEK. Clozapine-N-Oxide was kindly provided by the National Institute of Mental Health.

## CONFLICT OF INTEREST

BCR and MRH both receive research funding from Novo Nordisk and Boehringer Ingelheim that was not used in support of these studies. MRH receives research funding from Eli Lilly & Co., Gila Therapeutics, and Pfizer that was not used in support of these studies. MRH is CEO of Cantius Therapeutics, LLC which pursues biological work unrelated to the current study. All other authors declare no competing interests.

---

**Supplementary Fig. 1.**
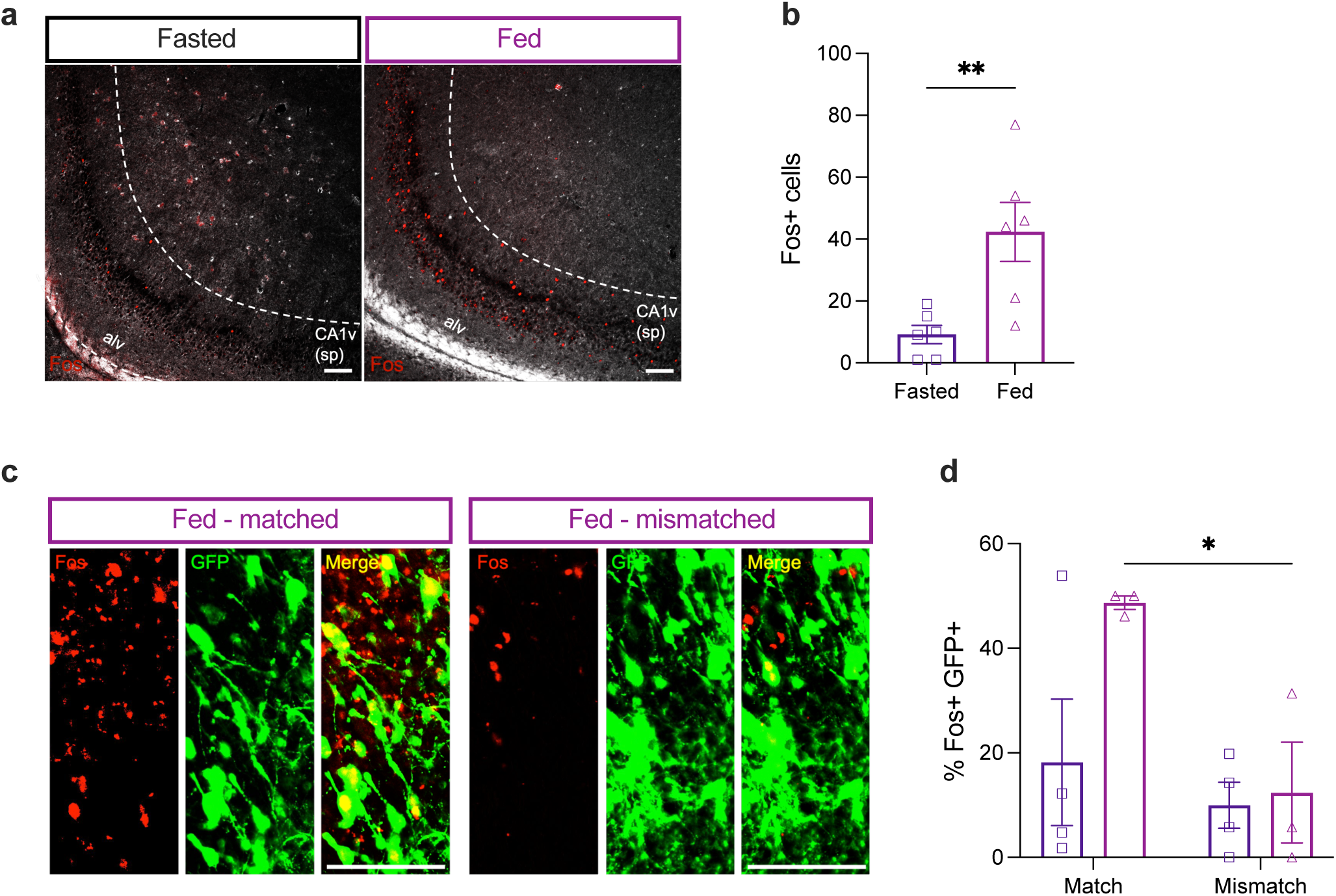
Validation of the targeted recombination in active population viral approach. (**a**) Representative photomicrographs of Fos immunolabelling in the pyramidal layer of the ventral CA1 (CA1v(sp)) of rats perfused under a Fasted or Fed state, relative to the alveus (alv); scale bar 100 um. (**b**) Quantification of Fos+ cells in the ventral CA1 of rats perfused under a Fasted or Fed state. (**c**) Representative photomicrograph of the ventral CA1v with immunolabelling for green fluorescent protein (GFP; green) and Fos (red) in a rat from the Fed group perfused under a Fed state (match) or Fasted state (mismatched); scale bar 100 um. (**d**) Percentage of ventral CA1 GFP+ cells that co-express Fos in rats from the Fed and Fasted group perfused under matched or mismatched conditions. Data are presented as mean ± SEM. For Extended Fig. 1b, Mann-Whitney test (n=6/group). For Extended Fig. 1d, two-way ANOVA, Tukey post hoc (n=3-4/group); *p<0.05, **p<0.01.

**Supplementary Fig. 2.**
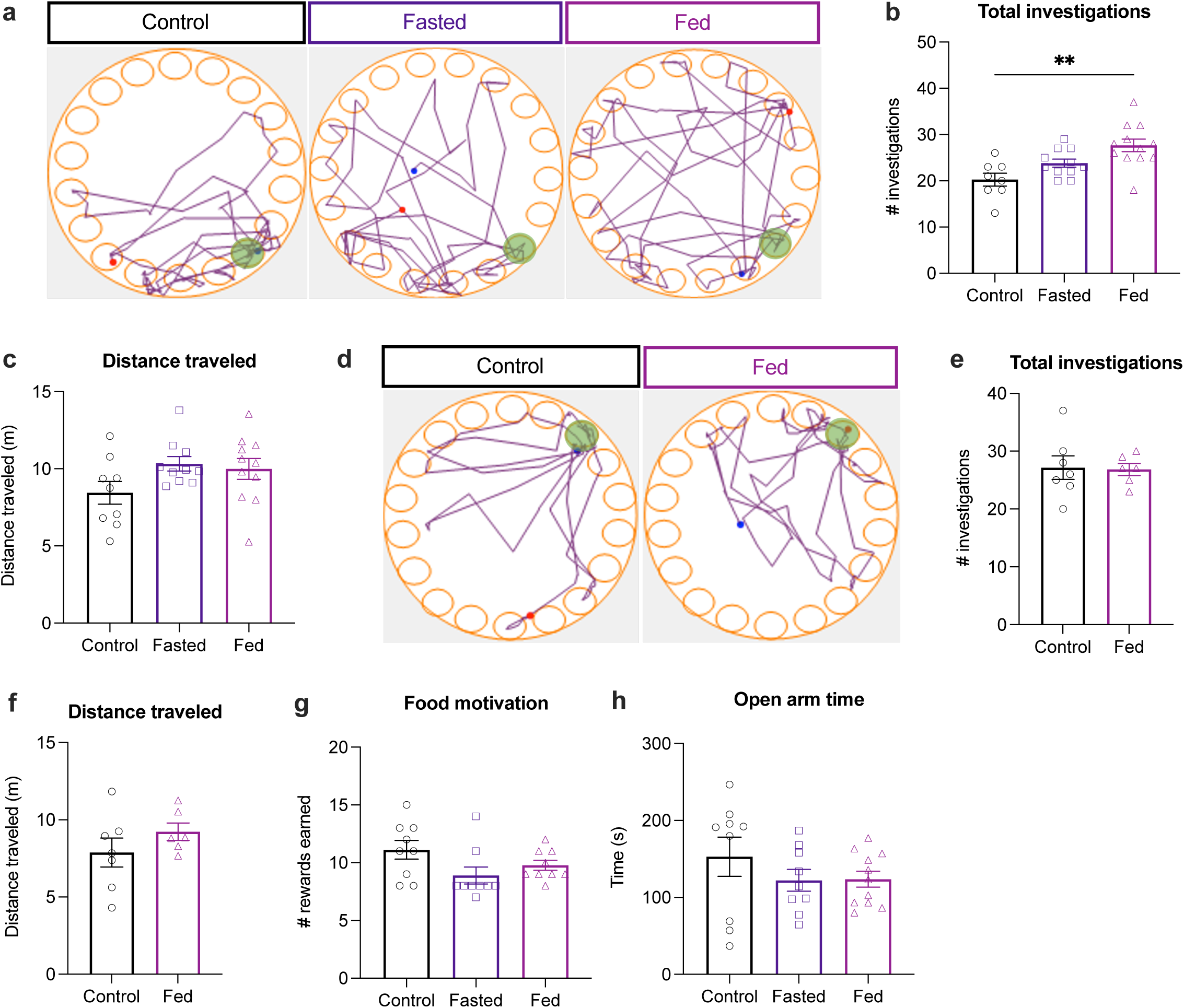
Ablation of ventral hippocampus meal-responsive neurons does not influence effort-based lever pressing for sucrose nor anxiety-like behavior. (**a**) Representative traces of individual animals trajectory during the foraging-related spatial memory probe with the correct hole in green. (**b**) Total number of investigations and (**c**) total distance travelled during the foraging-related spatial memory probe. (**d**) Representative traces of individual animals trajectory during the escape-based spatial memory probe. (**e**) Total number of investigations and (**f**) distance travelled during the escape-based spatial memory probe with the correct hole in green. (**g**) Number of sucrose pellets earned in an effort-based lever pressing task with a progressive ratio reinforcement schedule. (**h**) Time spent in the open arms of the zero-maze. Data are presented as mean ± SEM. For Extended Fig. 2b-c and Extended Fig. 2g-h, one-way ANOVA (n=9-11/group); **p<0.01. For Extended Fig. 2e-f, two-tailed unpaired t-test (n=6-7/group).

**Supplementary Fig. 3.**
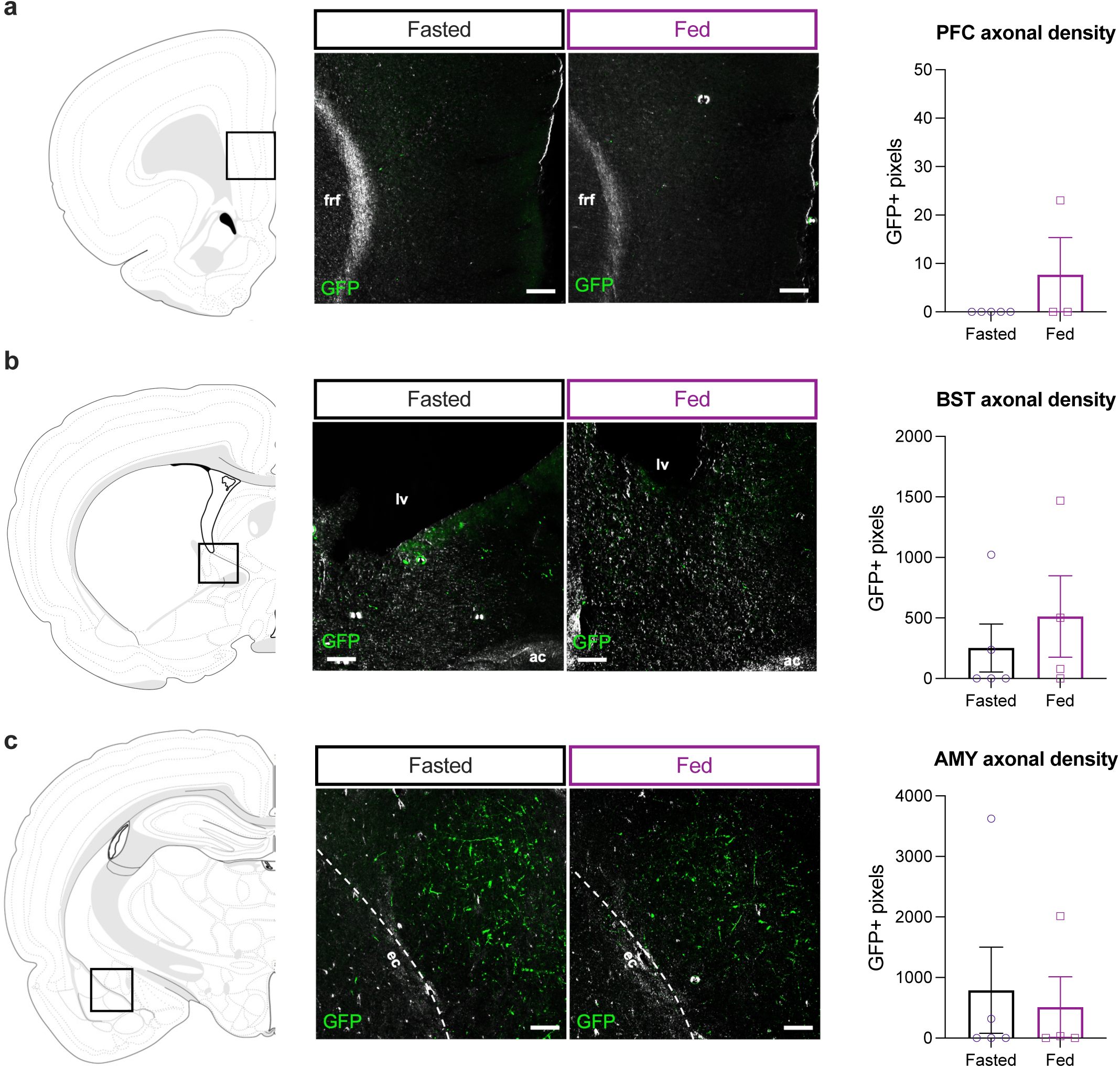
Ventral hippocampus meal and fasting-responsive neurons send minimal projections to the prefrontal cortex, bed of the stria terminalis and amygdala. (**a**) *Left:* Diagram of a coronal section of the prefrontal cortex (PFC). *Middle:* Representative photomicrographs of axonal green fluorescent protein (GFP)^+^ expression in the PFC from ventral CA1 neurons active in the Fasted or Fed state, relative to the fasciculus retroflexus (frf); scale bar 100 um. *Right:* Proportion of animals with GFP^+^ axons in the PFC. (**b**) *Left:* Diagram of a coronal section of the bed of the stria terminalis (BST). *Middle:* Representative photomicrographs of axonal GFP^+^ expression in the BST from ventral CA1 neurons active in the Fasted or Fed state, relative to the anterior commissure (ac) and lateral ventricle (lv); scale bar 100 um. *Right:* Proportion of animals with GFP^+^ axons in the BST. (**c**) *Left:* Diagram of a coronal section of the amygdala (AMY). *Middle:* Representative photomicrographs of axonal GFP^+^ expression in the AMY from ventral CA1 neurons active in the Fasted or Fed state, relative to the external capsule (ec); scale bar 100 um. *Right:* Proportion of animals with GFP^+^ axons in the AMY. Data are presented as mean ± SEM. Two-tailed unpaired t-test (n=4-5/group).

**Supplementary Fig. 4.**
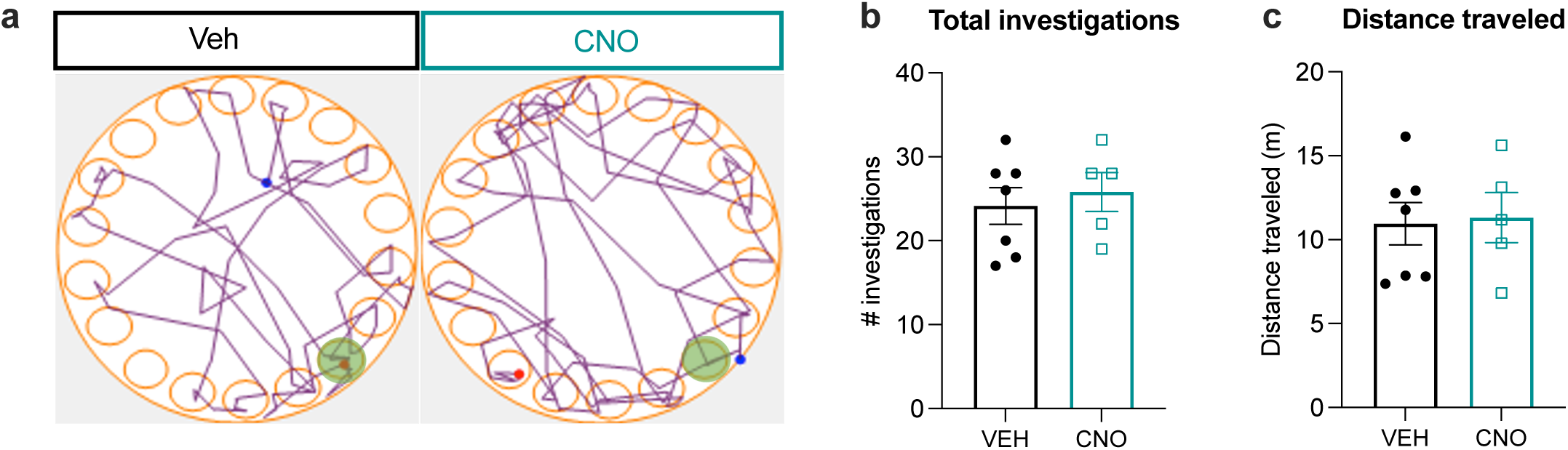
Ventral hippocampus to lateral hypothalamic area signaling contributes to foraging-related spatial memory. (**a**) Representative traces of individual animals trajectory during the foraging-related spatial memory probe with the correct hole in green. (**b**) Total number of investigations and (**c**) total distance travelled during the foraging-related spatial memory probe. Data are presented as mean ± SEM. For Extended Data Fig. 4b-c two-tailed paired t-test (n=4-6/group).

**Supplementary Fig. 5.**
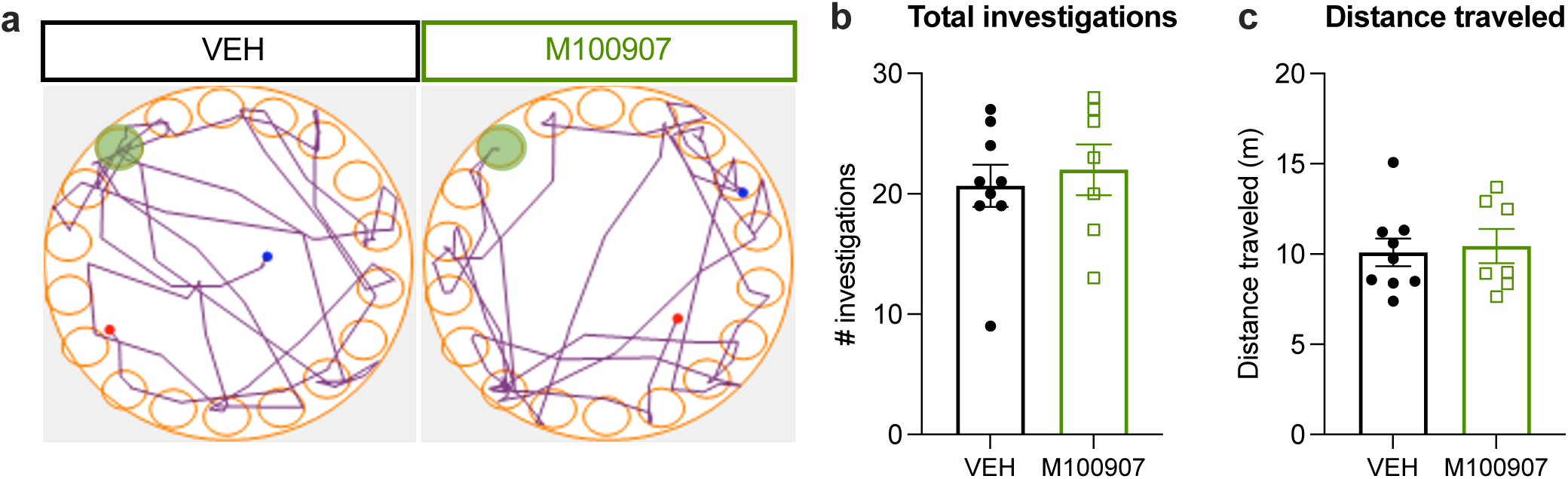
Ventral hippocampus 5HT2aR signaling contributes to foraging-related spatial memory. (**a**) Representative traces of individual animals trajectory during the foraging-related spatial memory probe with the correct hole in green. (**b**) Total number of investigations and (**c**) total distance travelled during the foraging-related spatial memory probe.

**Supplemental Table 1.** Number of differentially expressed genes by cell cluster1.

| Cluster Number | Cluster Label | DEGs - Down | DEGs - Up | DEGs Sum |
| --- | --- | --- | --- | --- |
| 1 | InNeuro, Sst+ | 61 | 14 | 75 |
| 2 | InNeuro, Sv2c+ | 6 | 3 | 9 |
| 3 | InNeuro, Vip+ | 16 | 8 | 24 |
| 4 | Granule, ExNeuro, Prox1+ | 53 | 65 | 118 |
| 5 | CA1 ExNeuro, Cpne7+ Dcn+ | 3 | 2 | 5 |
| 6 | CA1 ExNeuro, Cpne7+ Dcn+ | 224 | 161 | 385 |
| 7 | CA1 excitatory, Cpne7+ Dcn+ | 43 | 37 | 80 |
| 8 | CA1 or CA3 Excitatory, Cpne7+ | 100 | 17 | 117 |
| 9 | CA1 or CA3 Excitatory, Cpne7+ Dcn+ | 31 | 31 | 62 |
| 10 | CA1 or CA3 Excitatory, Cpne7+ | 19 | 15 | 34 |
| 11 | Oligodendrocytes | 73 | 129 | 202 |
| 12 | Oligodendrocyte precursor cells | 0 | 0 | 0 |
| 13 | Oligodendrocyte precursor cells | 34 | 6 | 40 |
| 14 | Astrocytes | 40 | 50 | 90 |
| 15 | Endothelial | 44 | 301 | 345 |
| 16 | Macrophages | 0 | 0 | 0 |
| 17 | Microglia | 18 | 11 | 29 |

**Supplemental Table 2.**
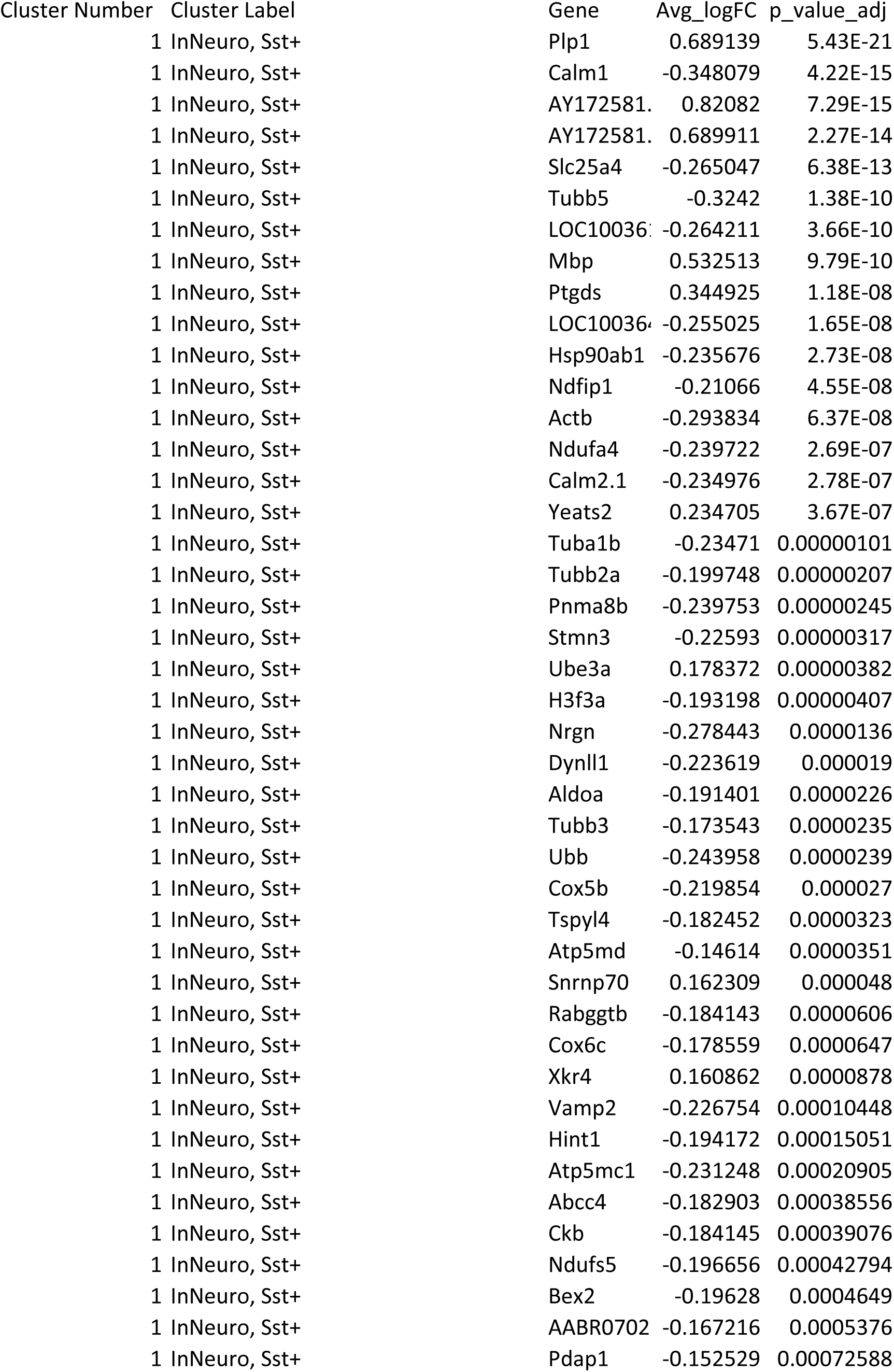
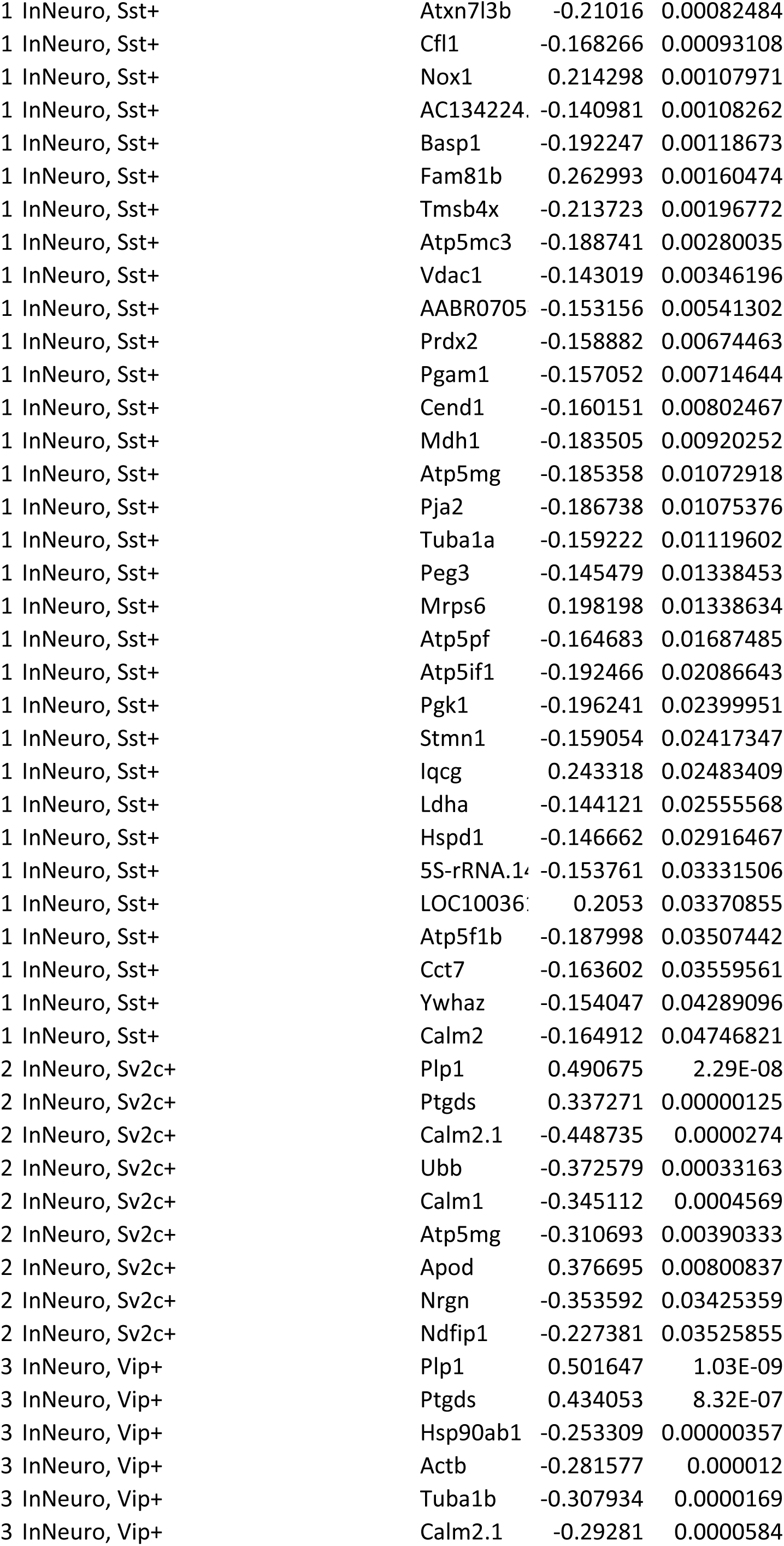
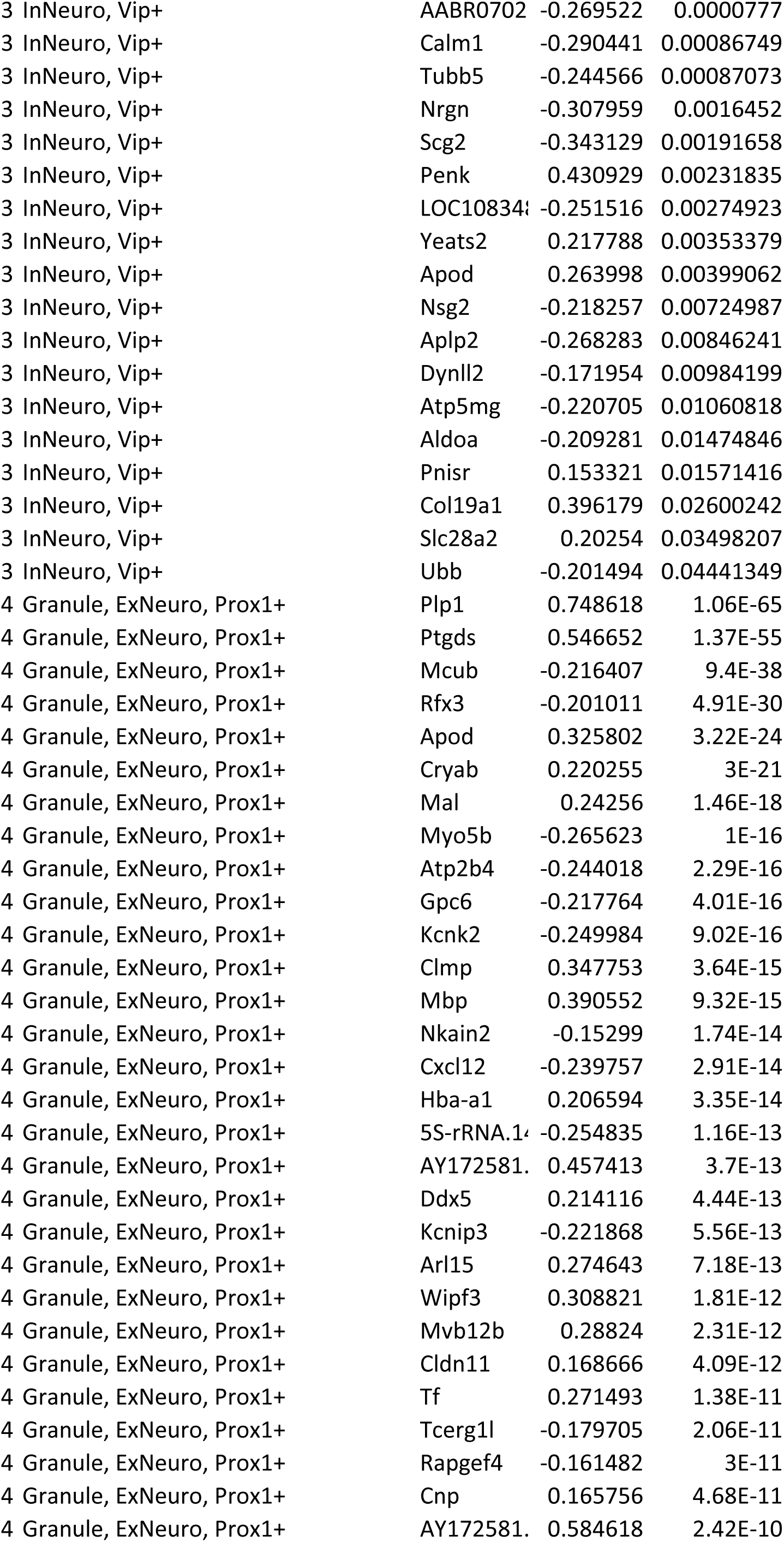
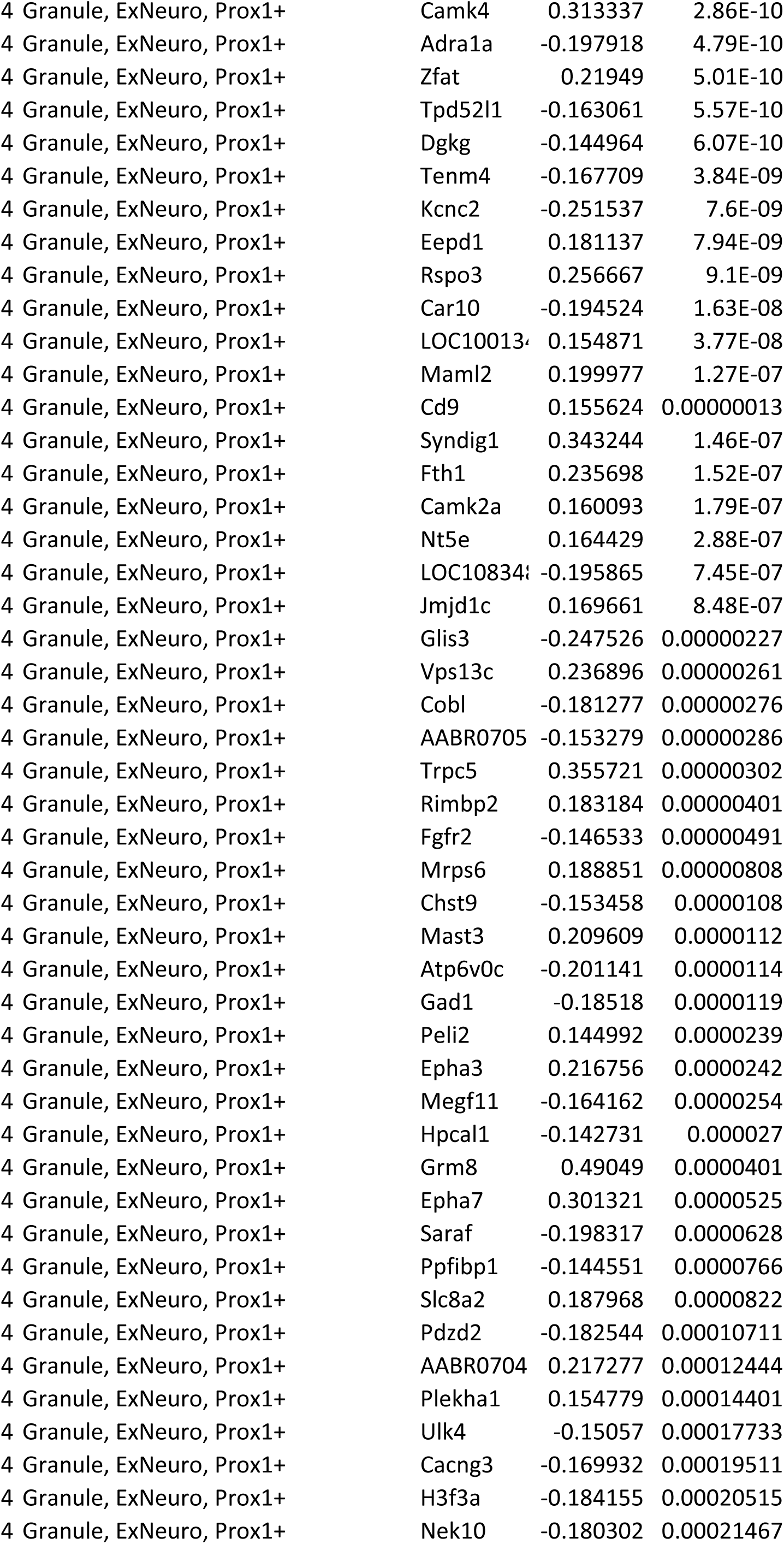
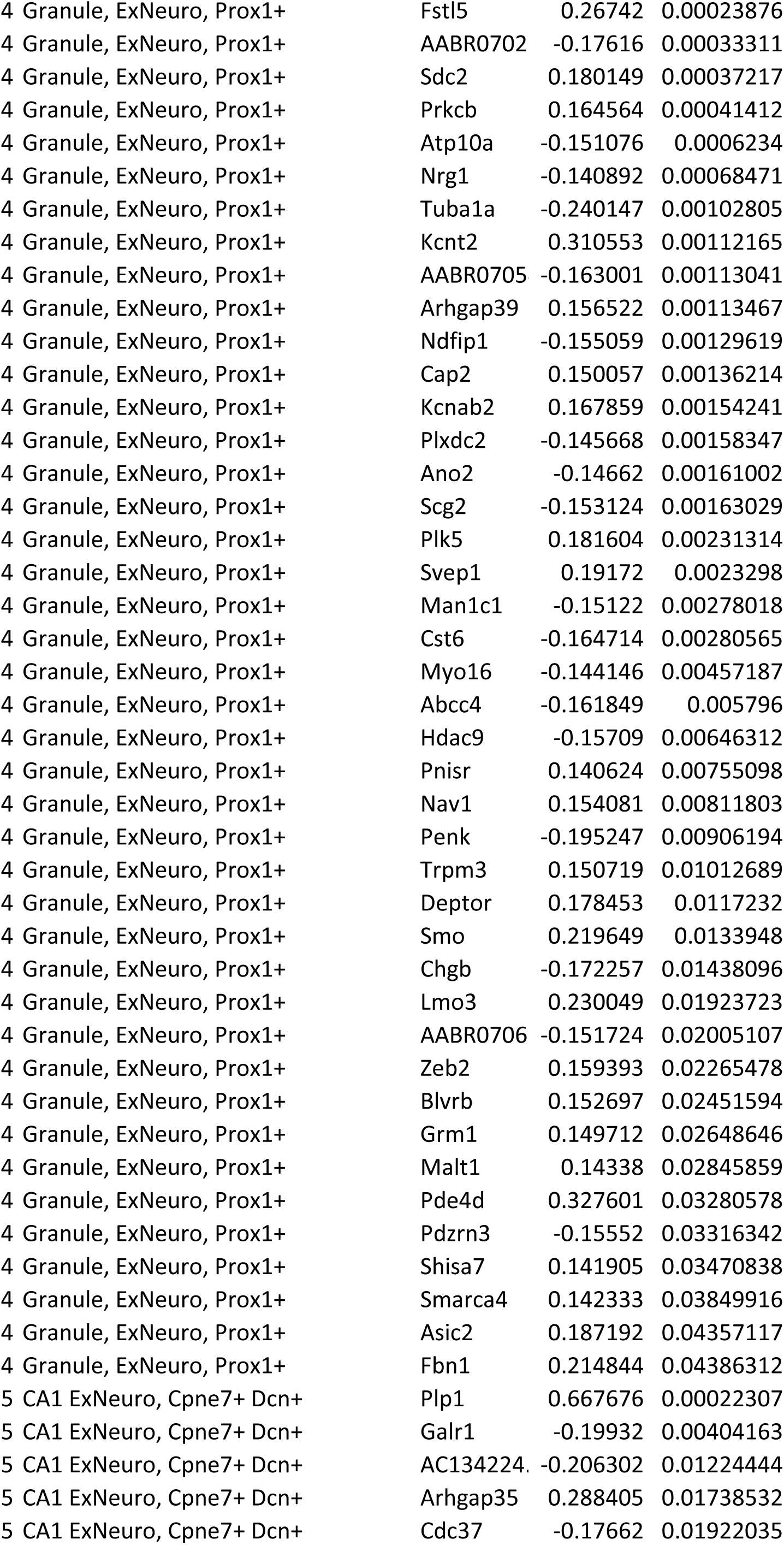
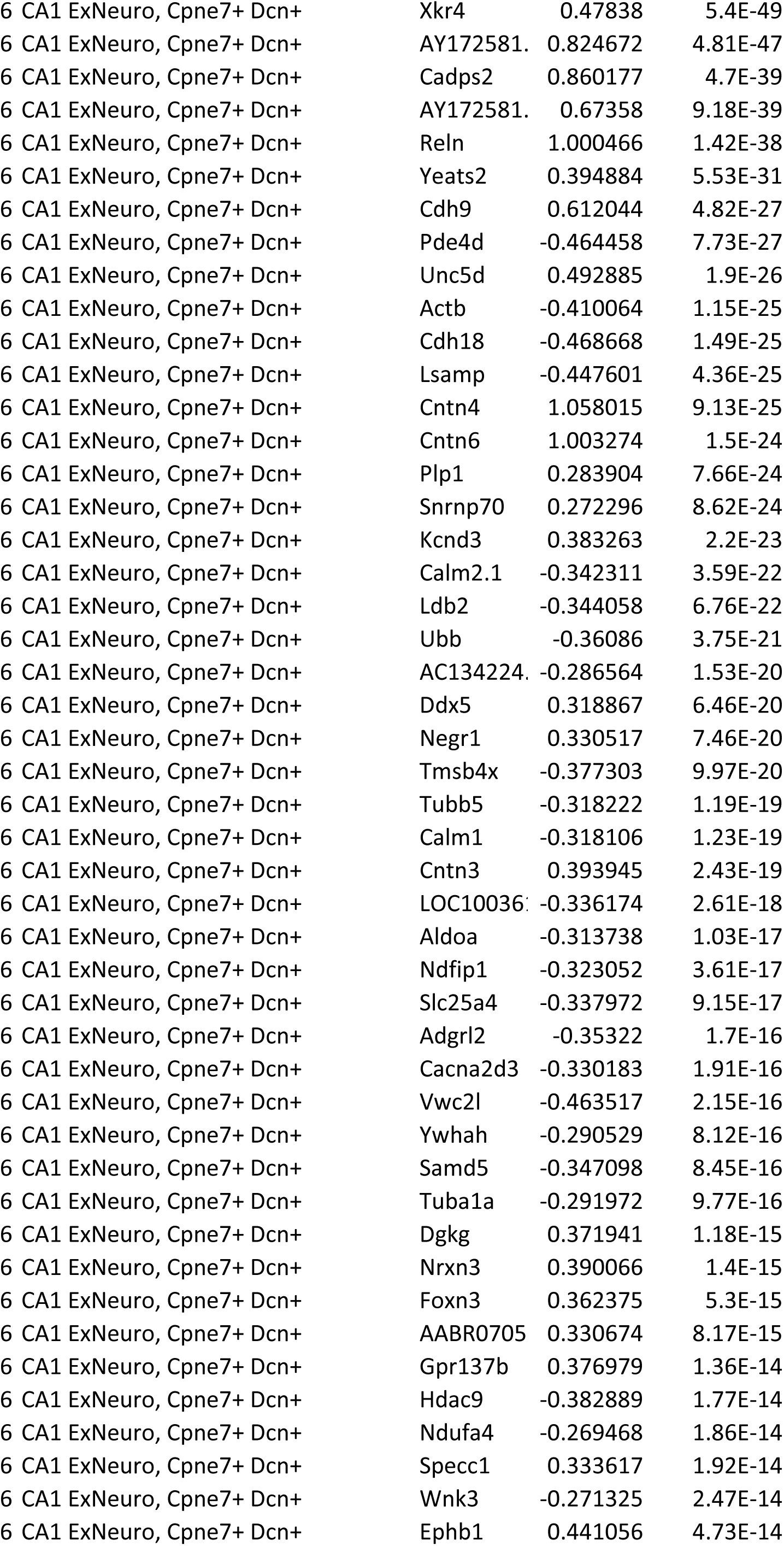
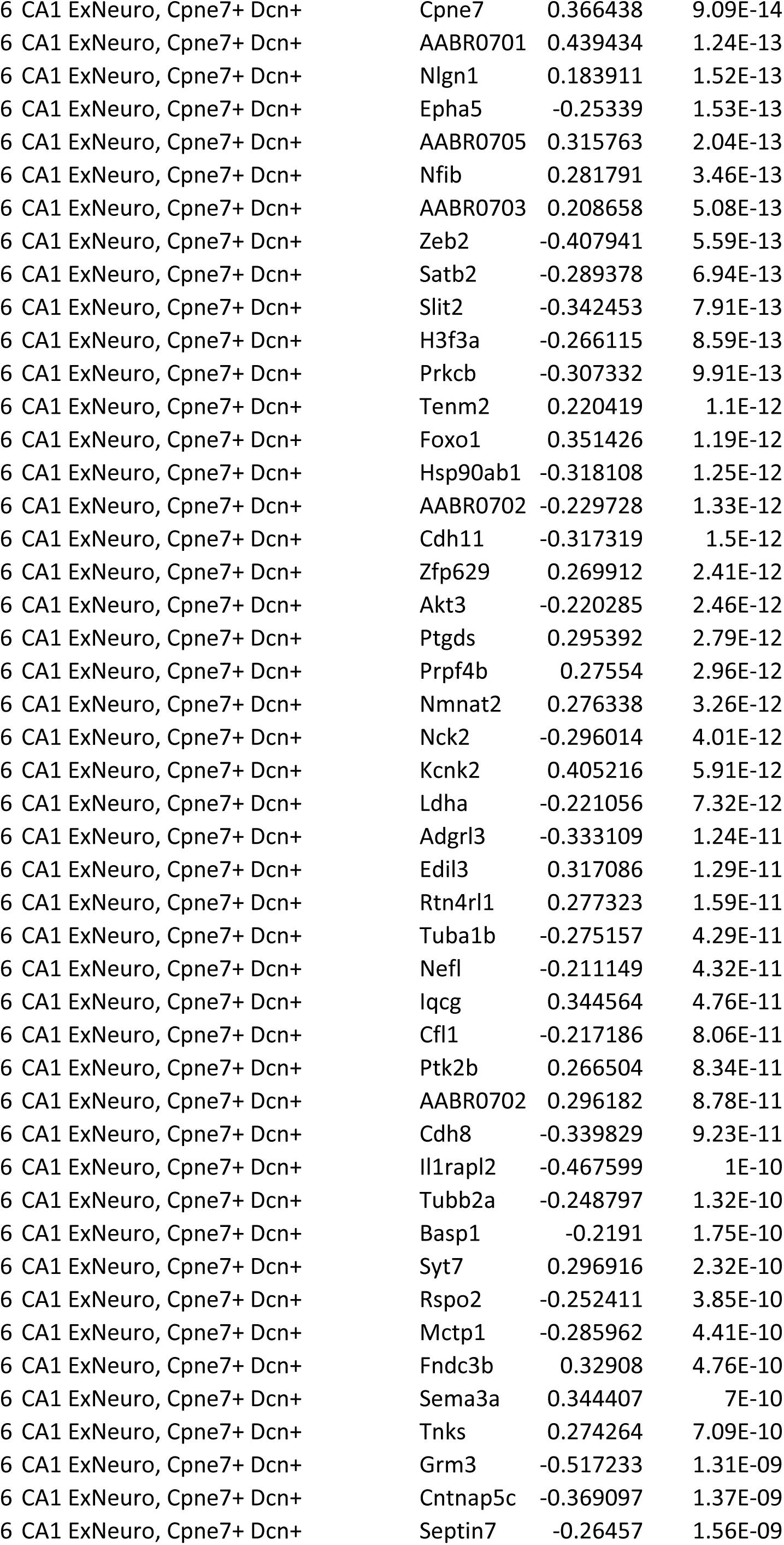
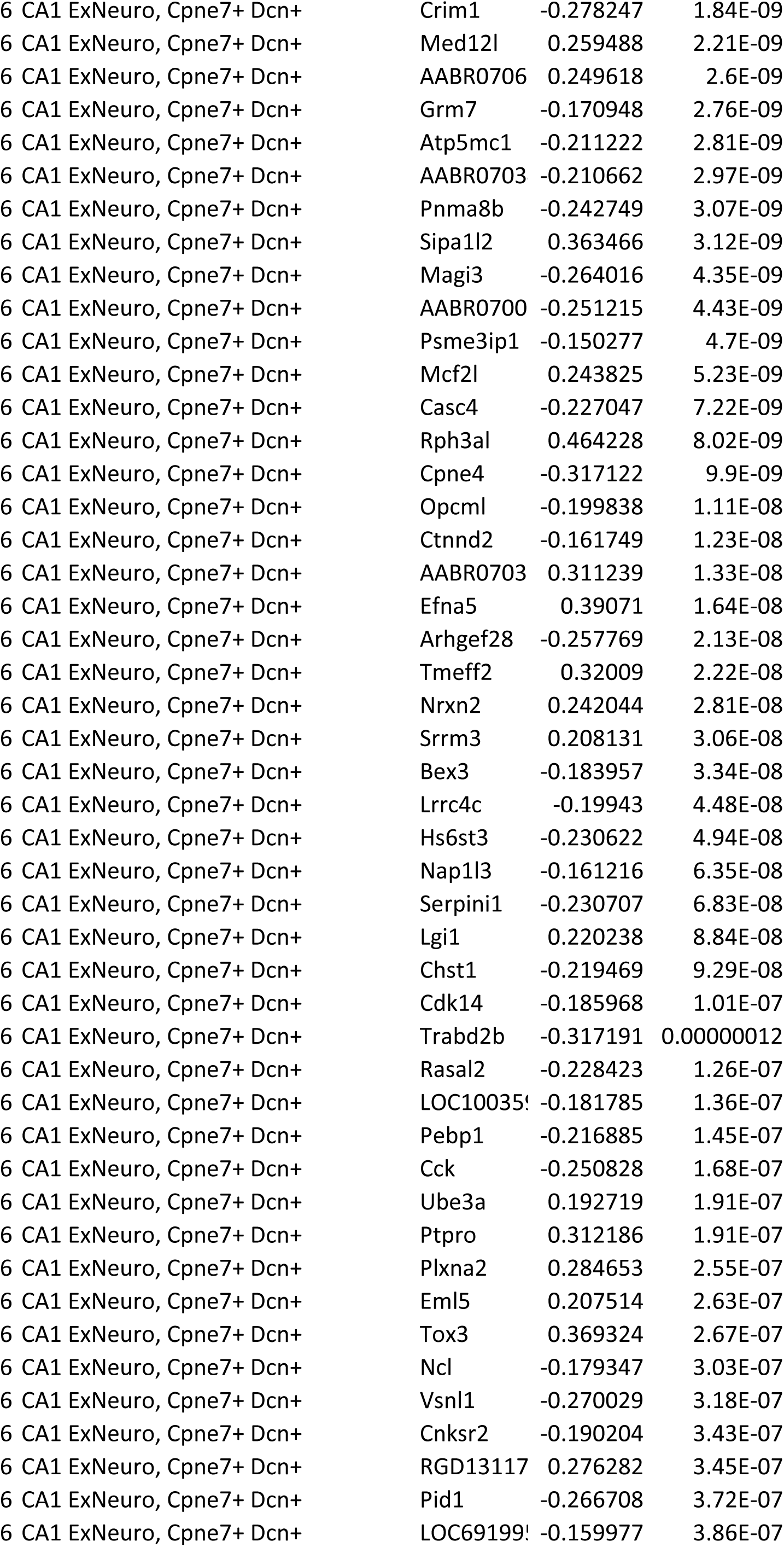
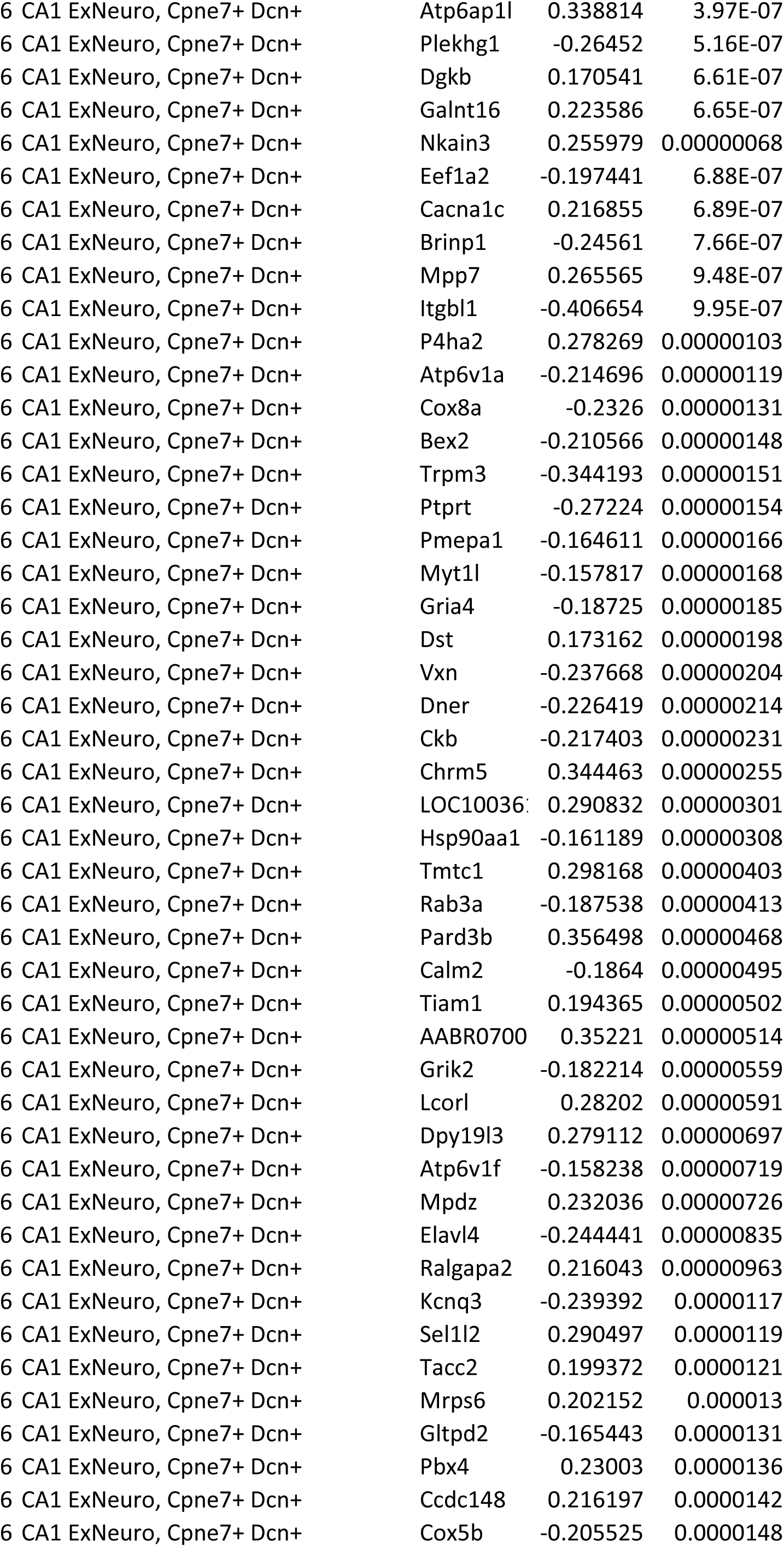
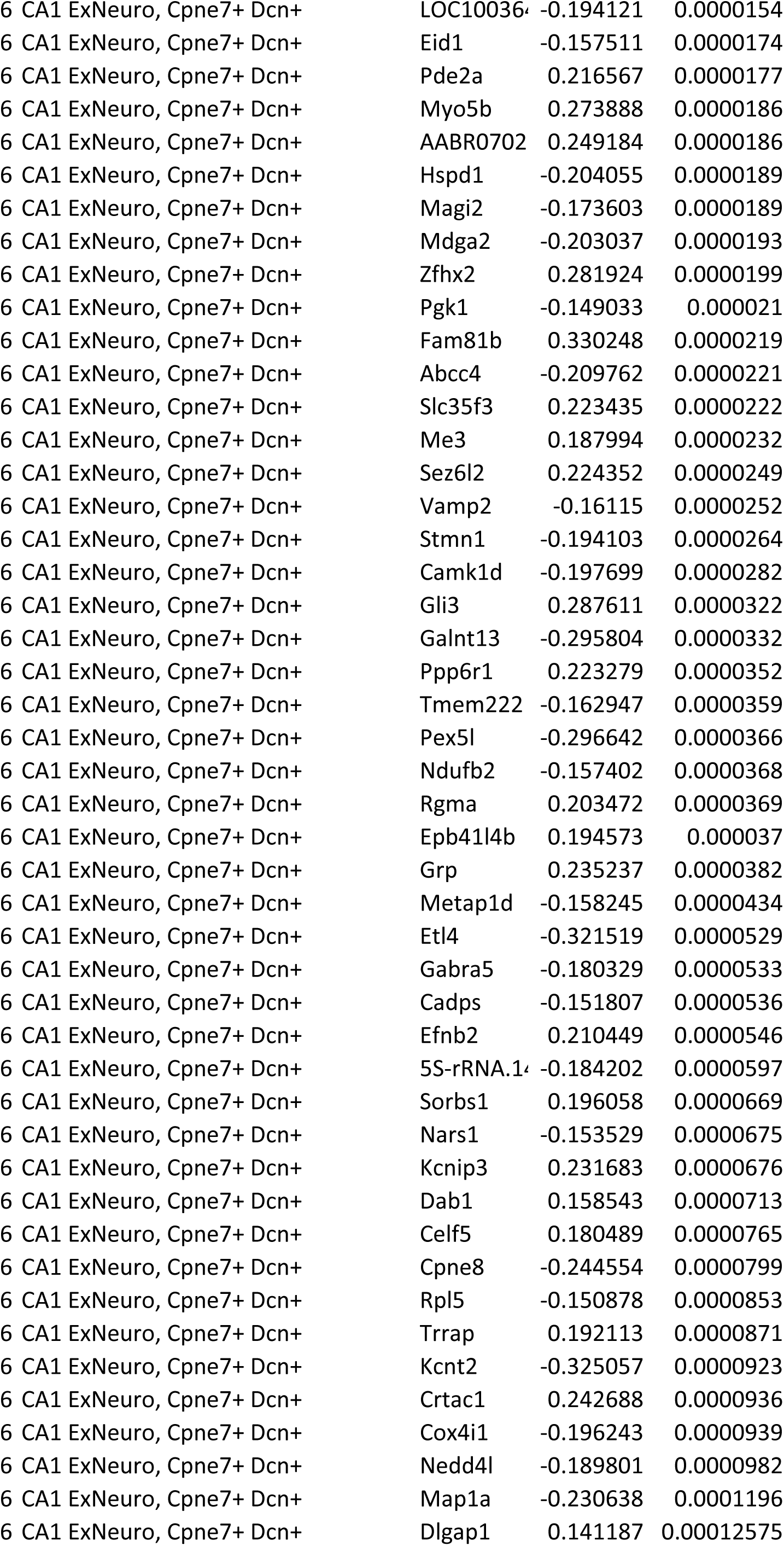
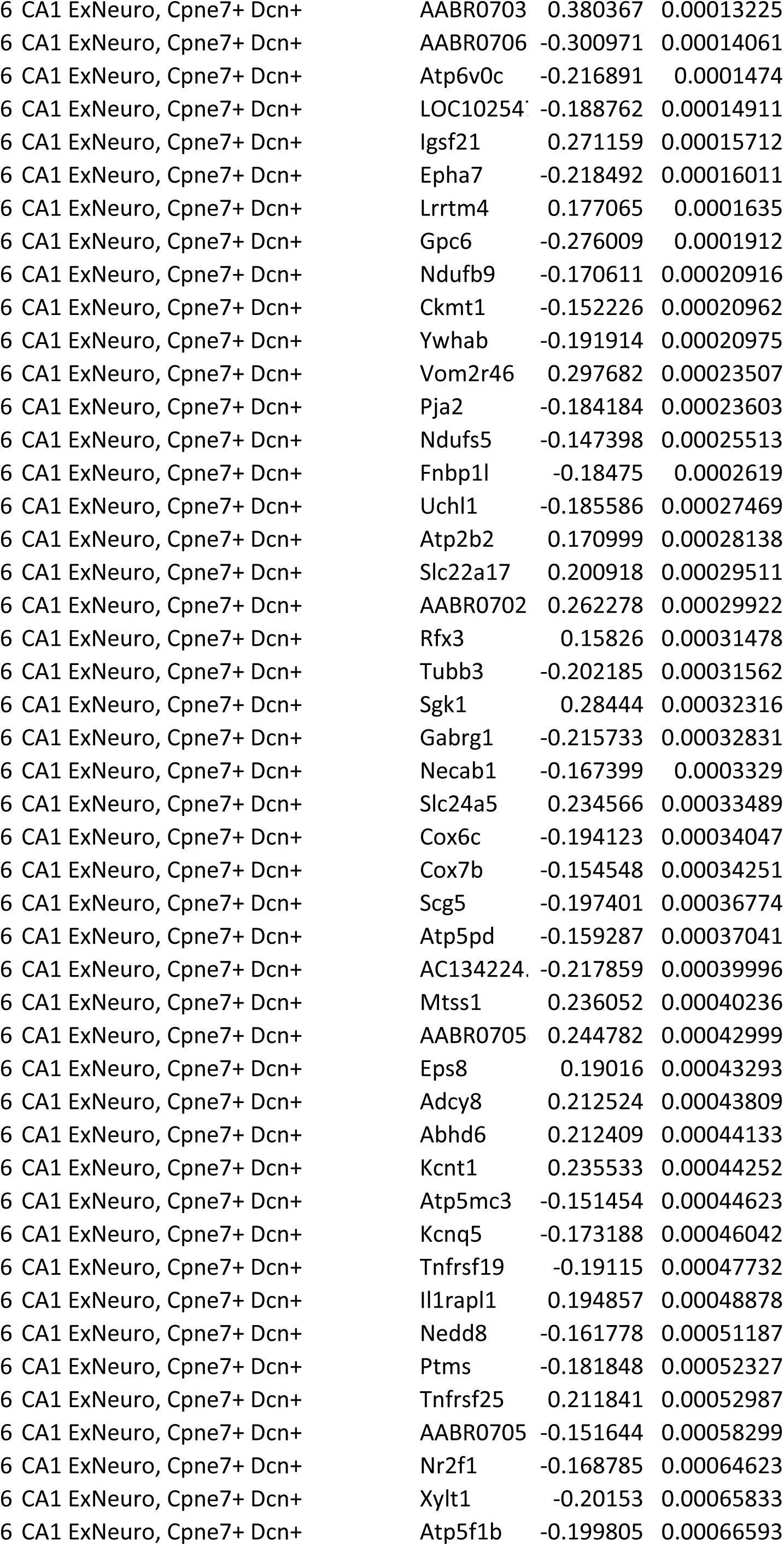
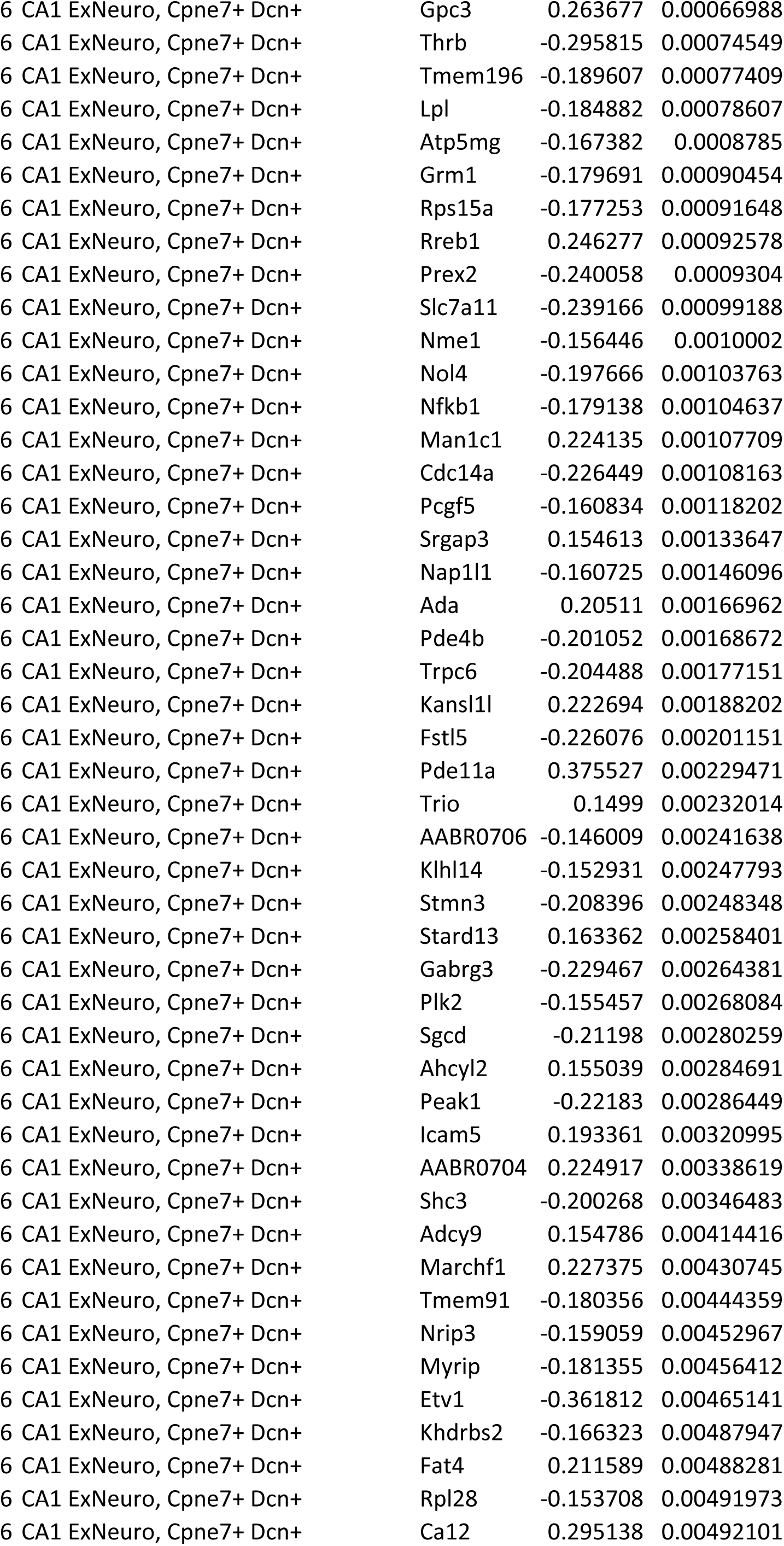
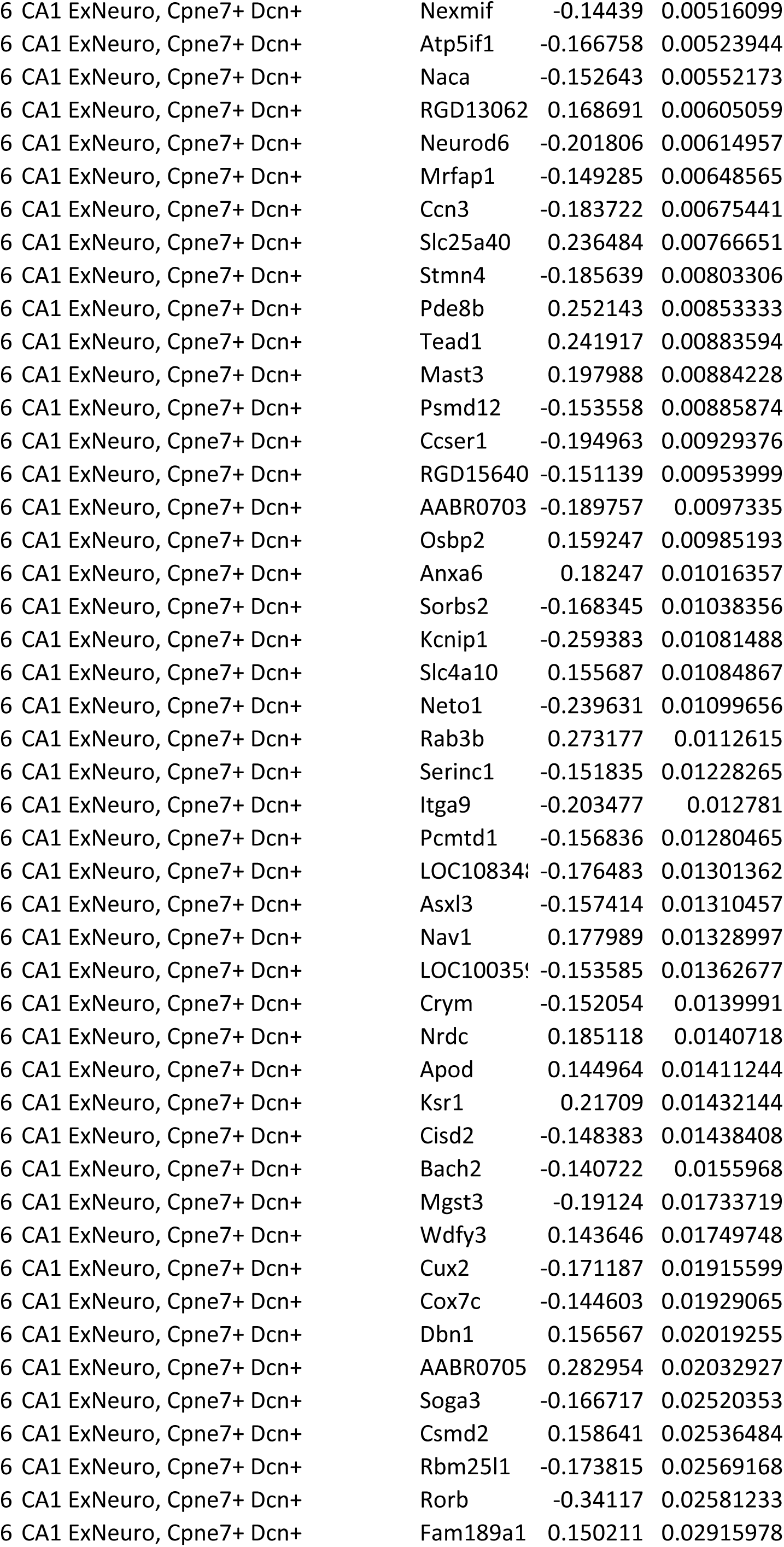
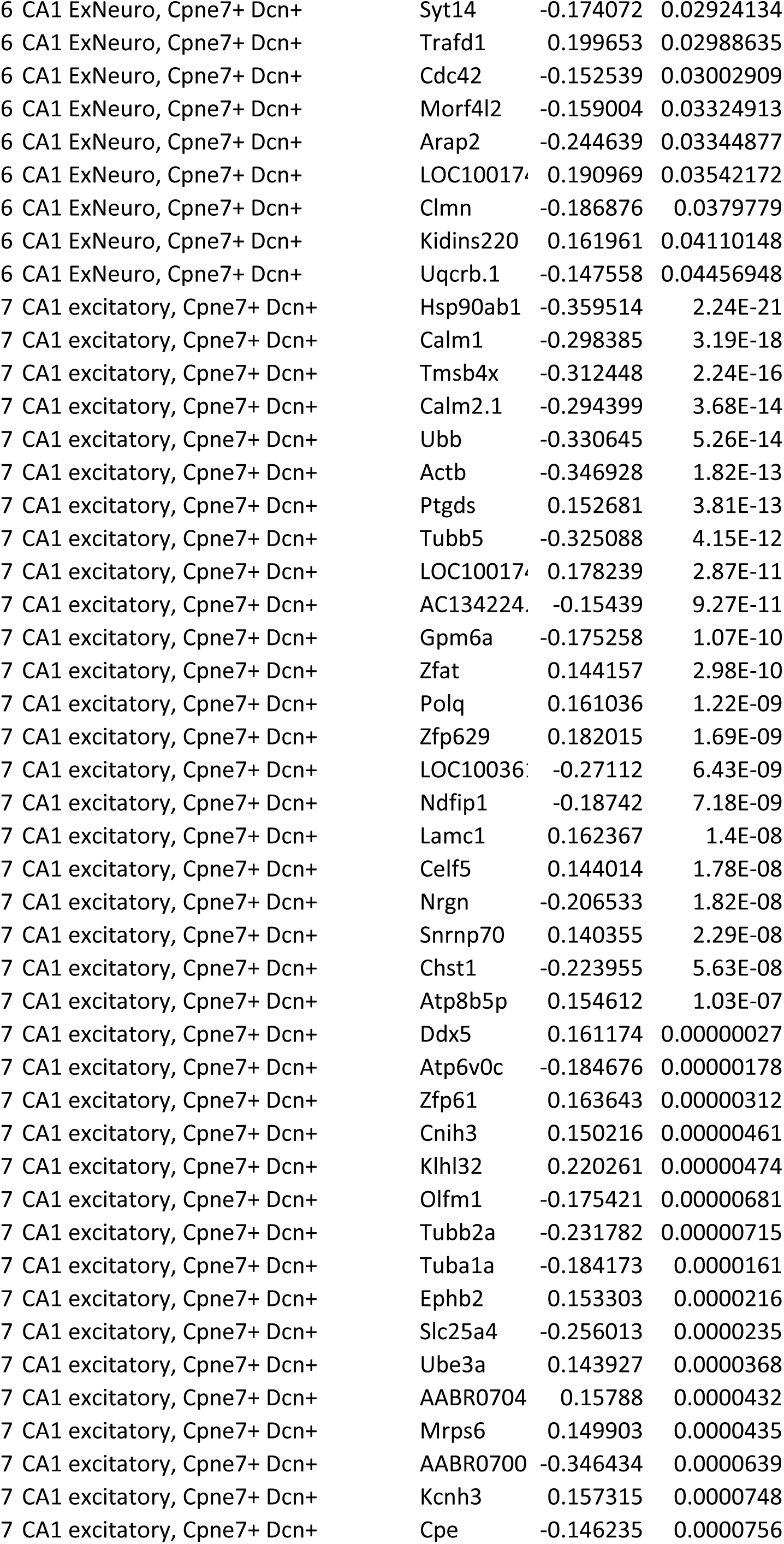
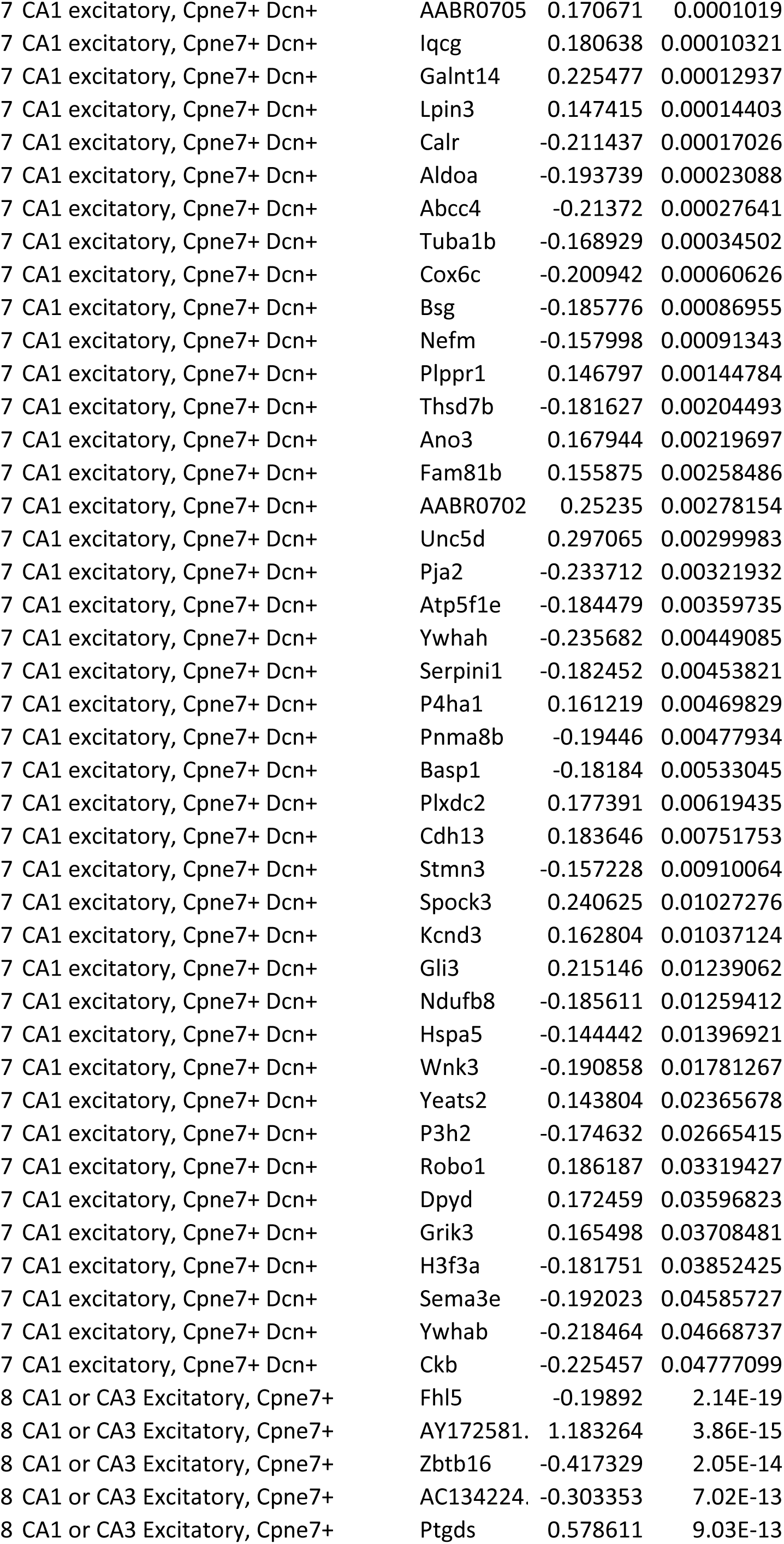
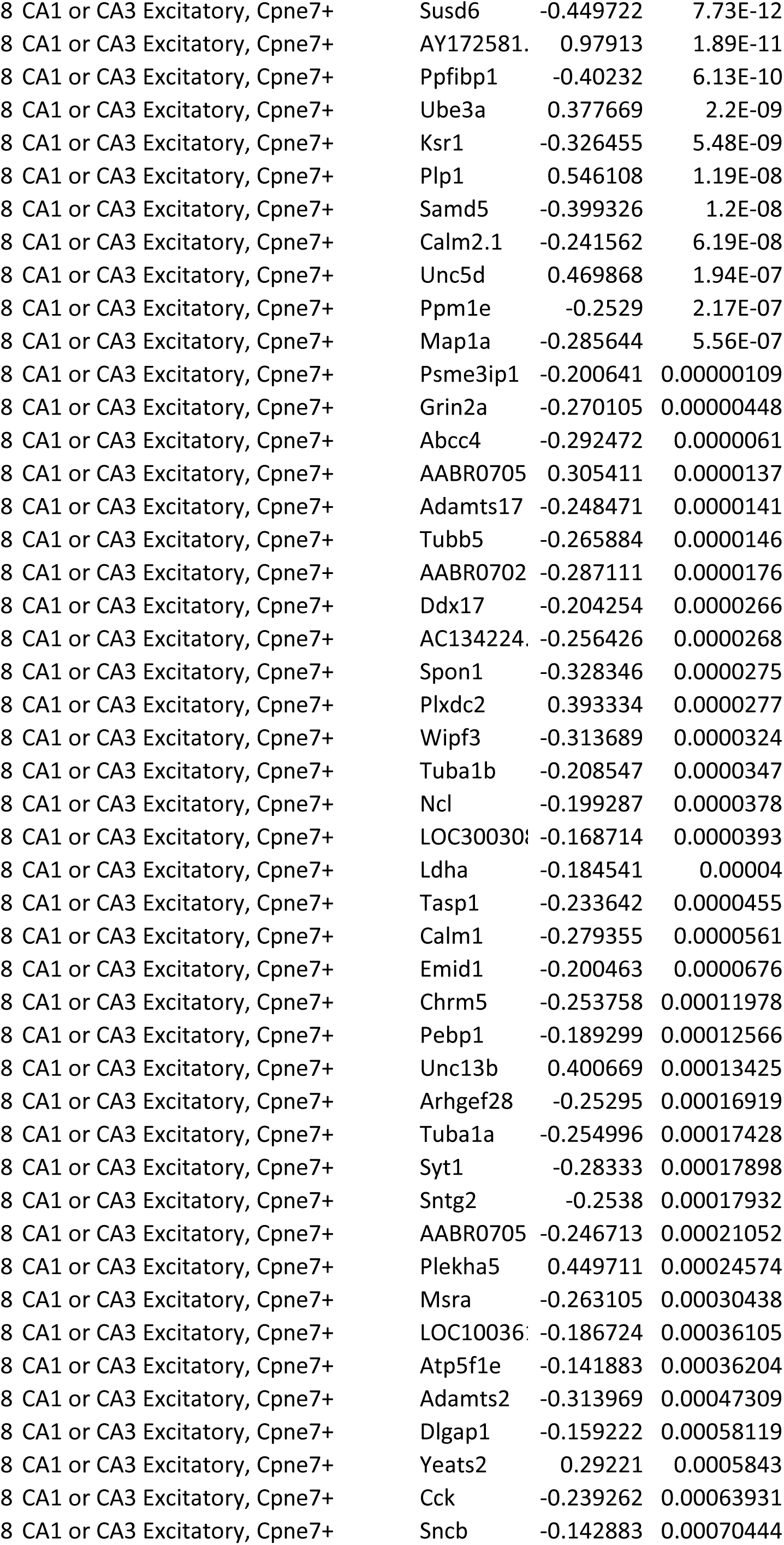
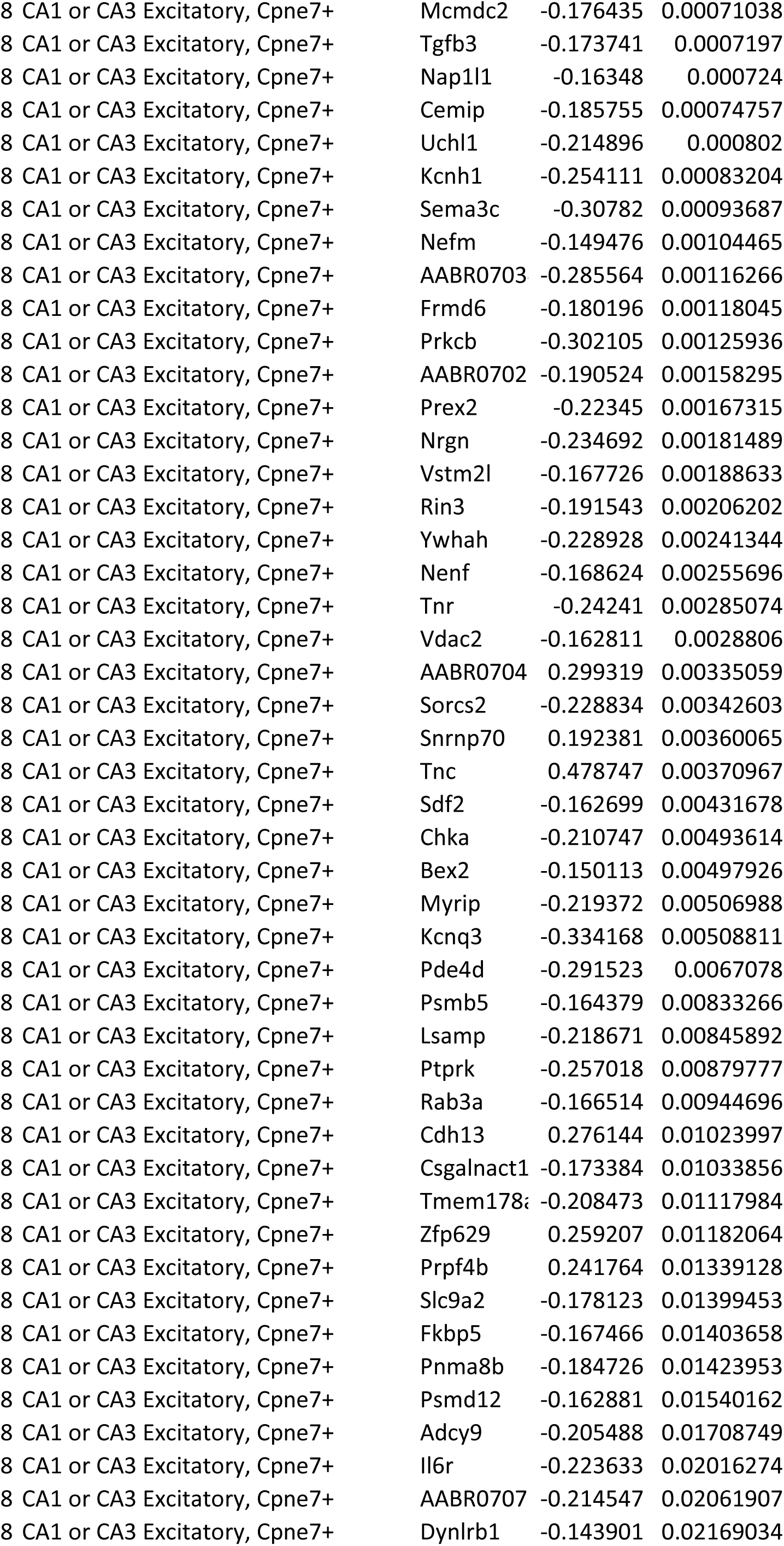
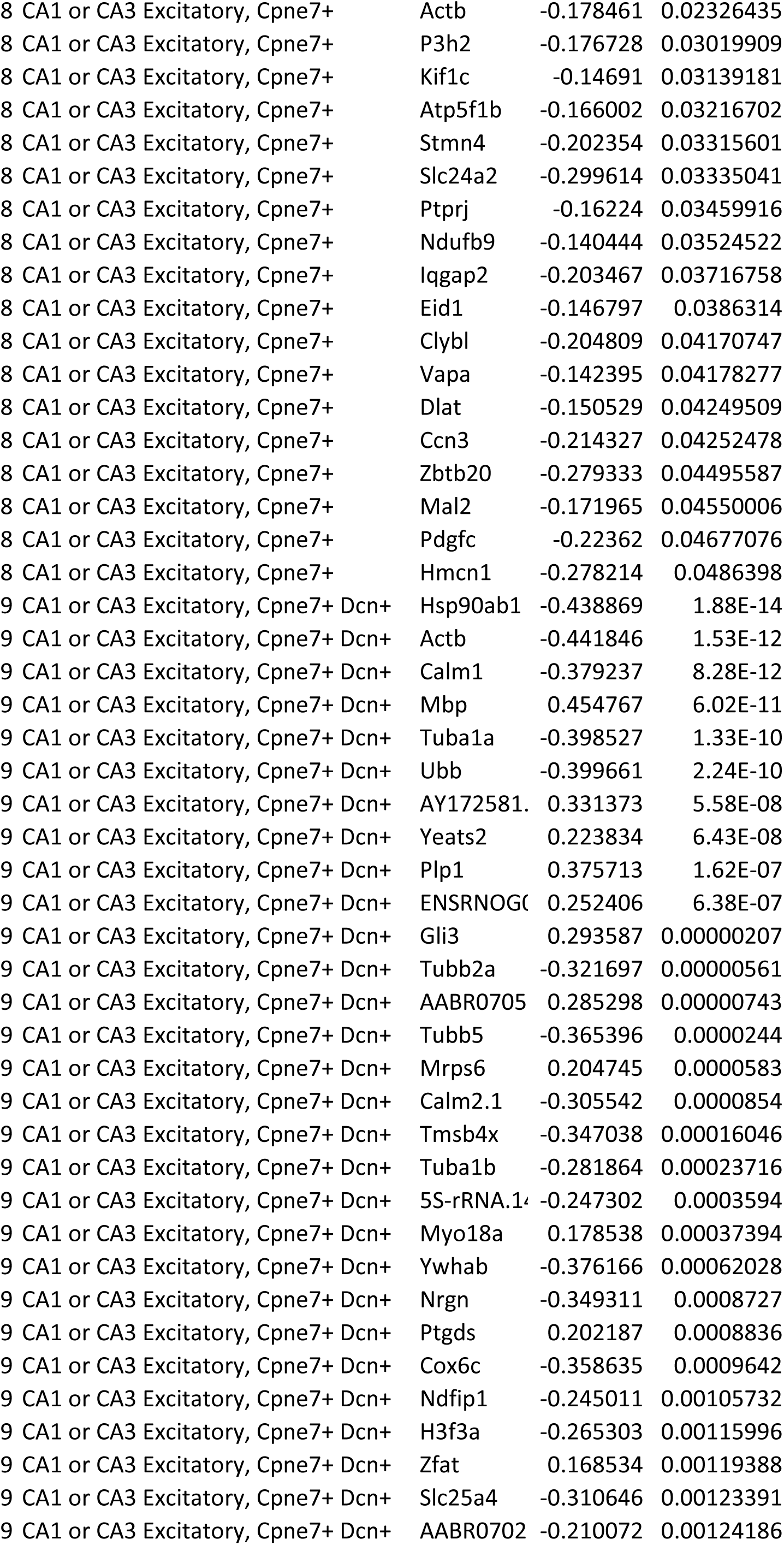
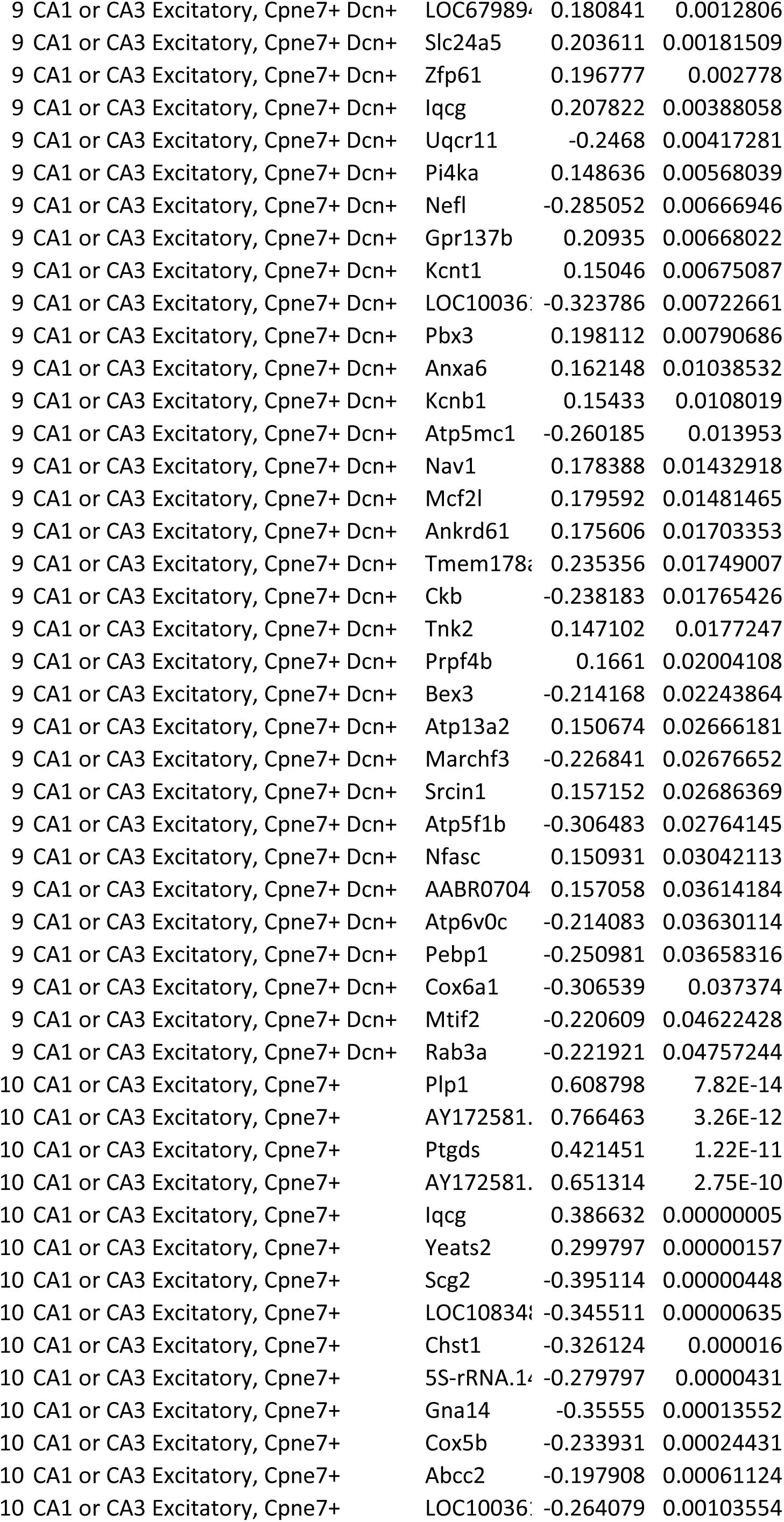
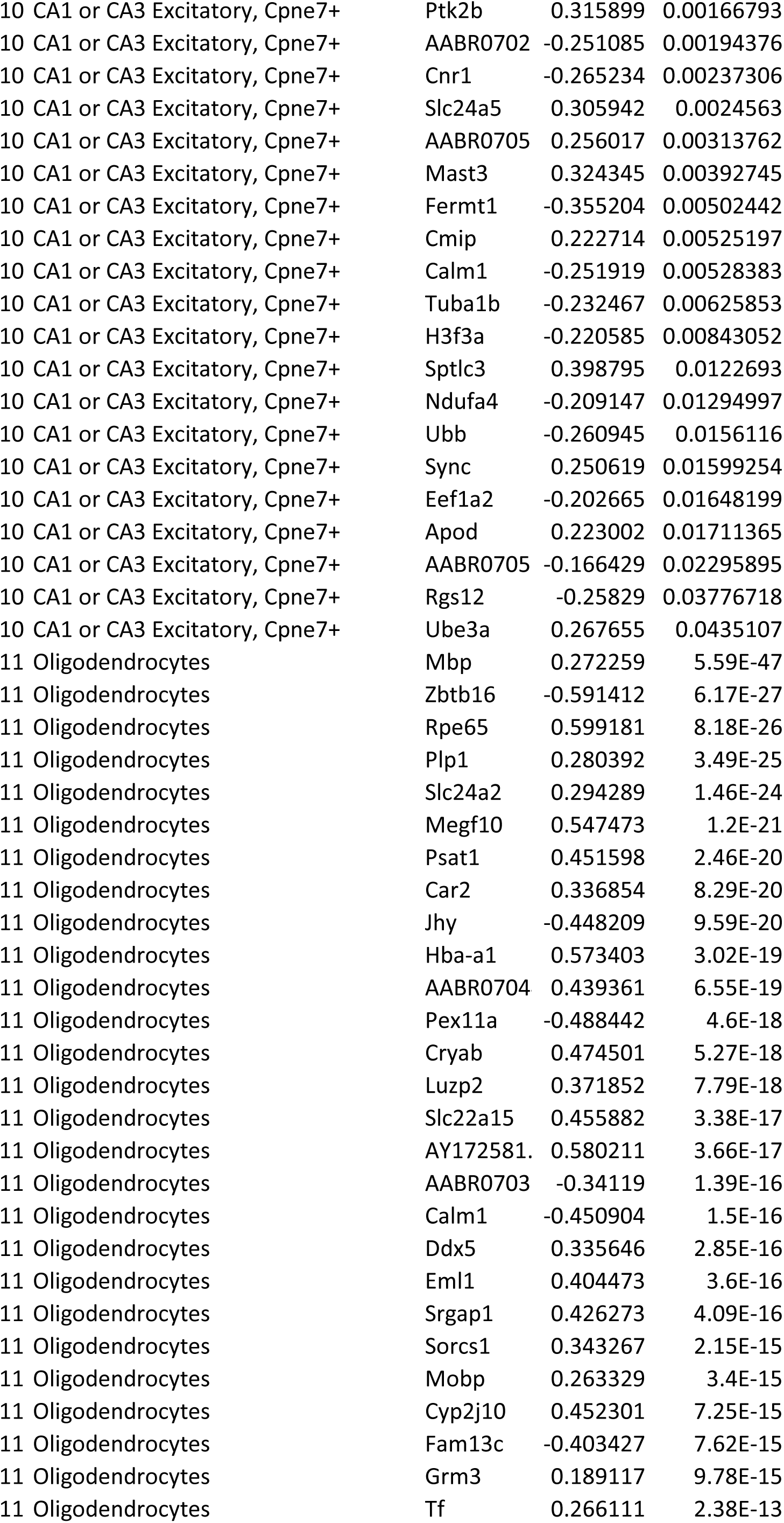
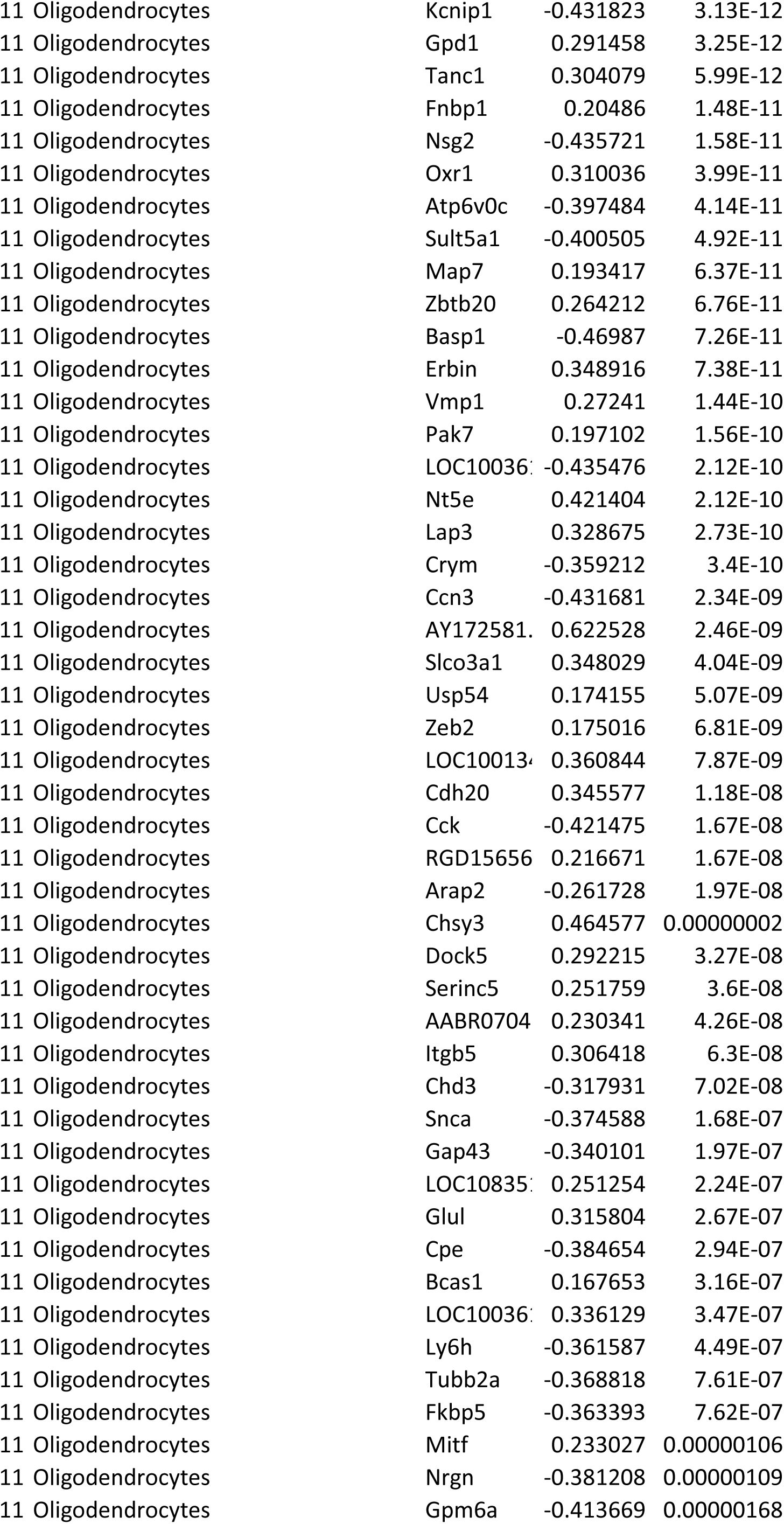
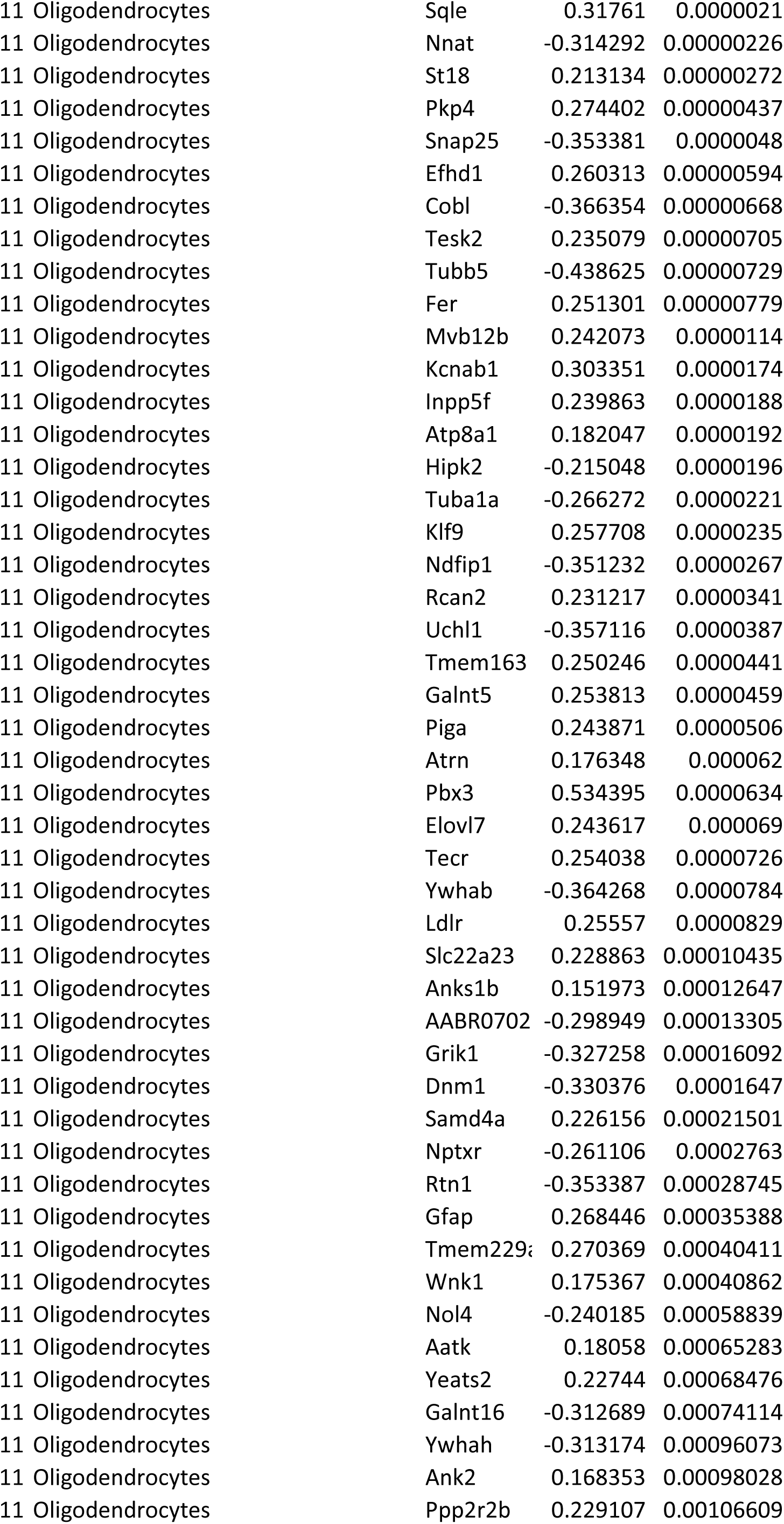
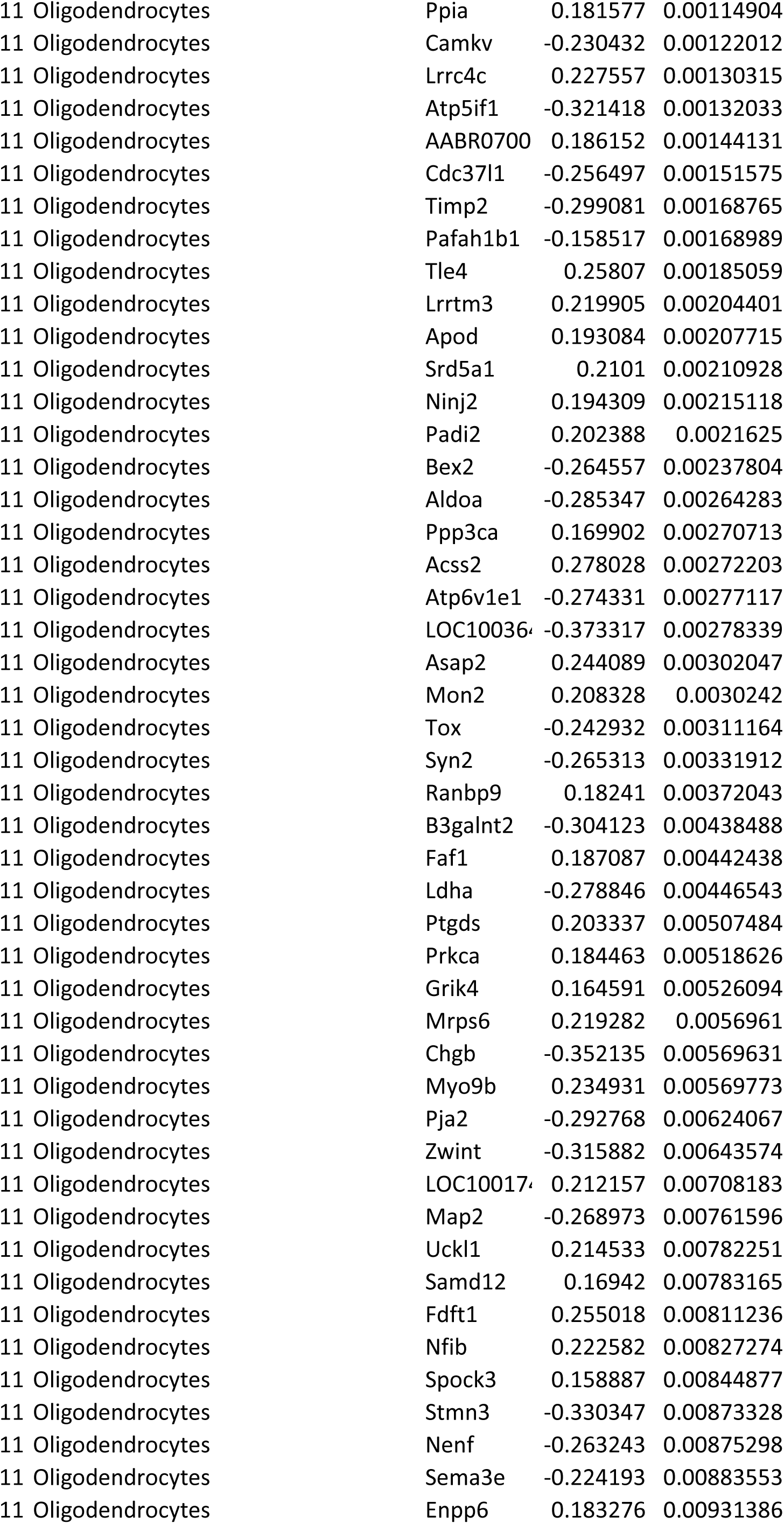
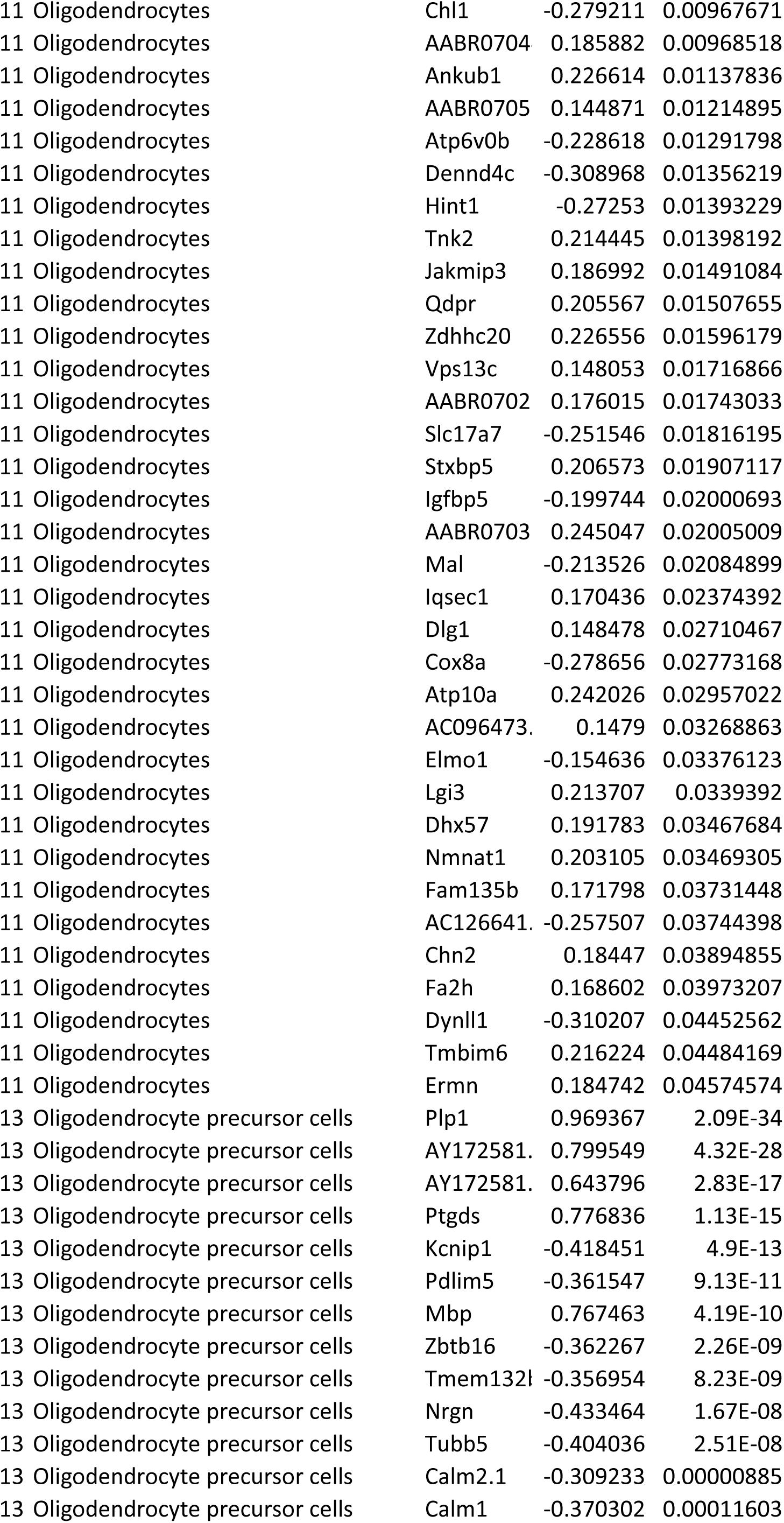
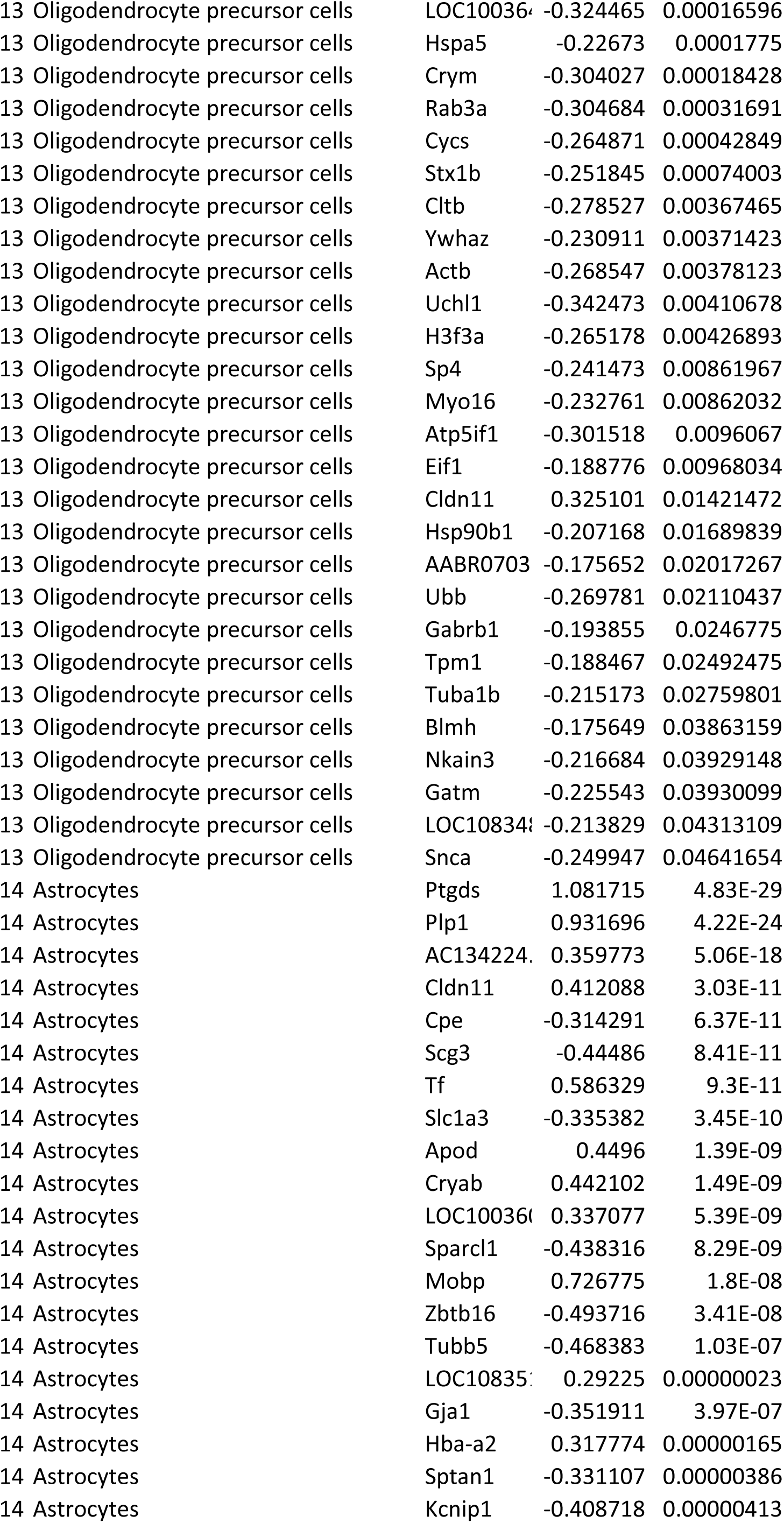
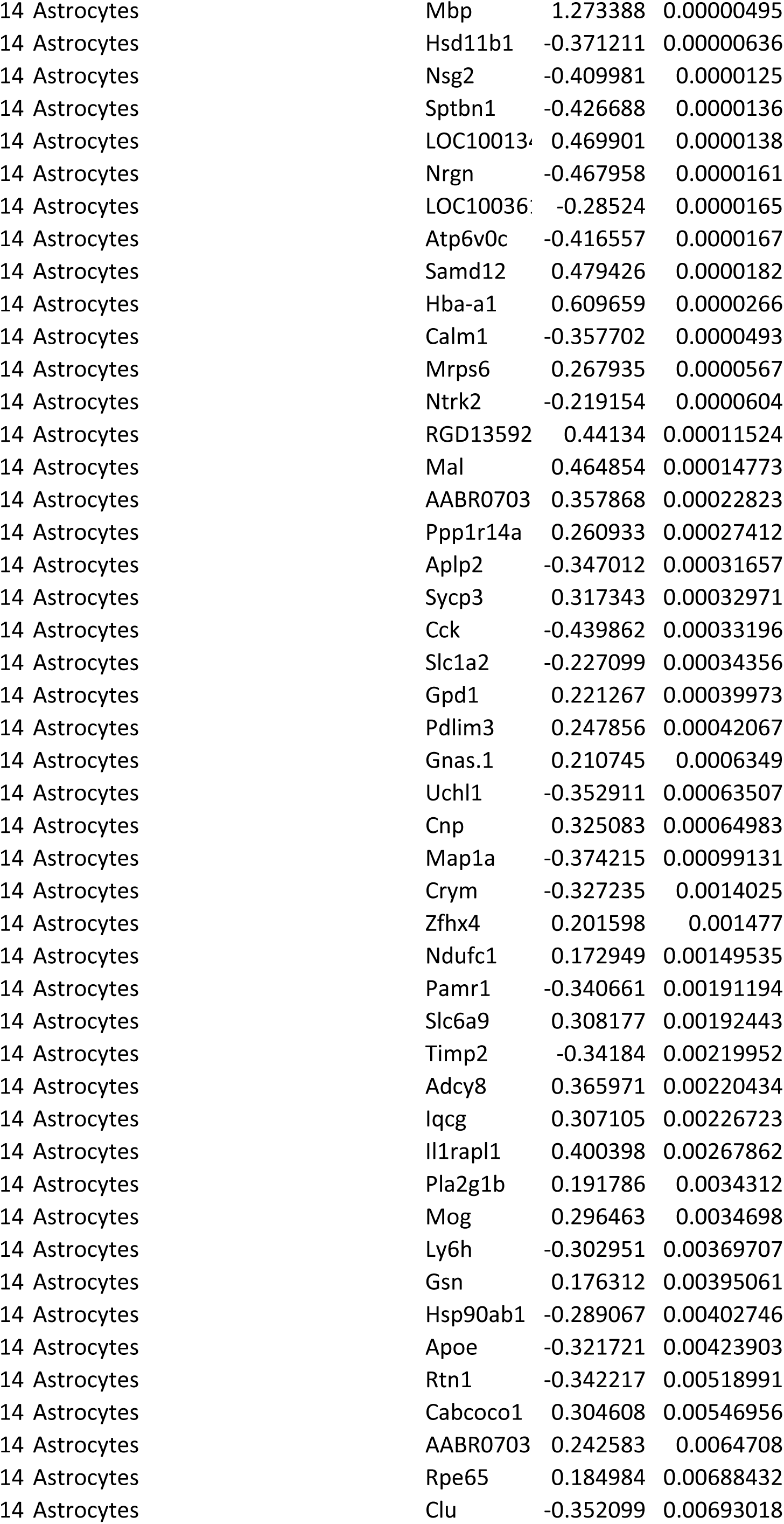
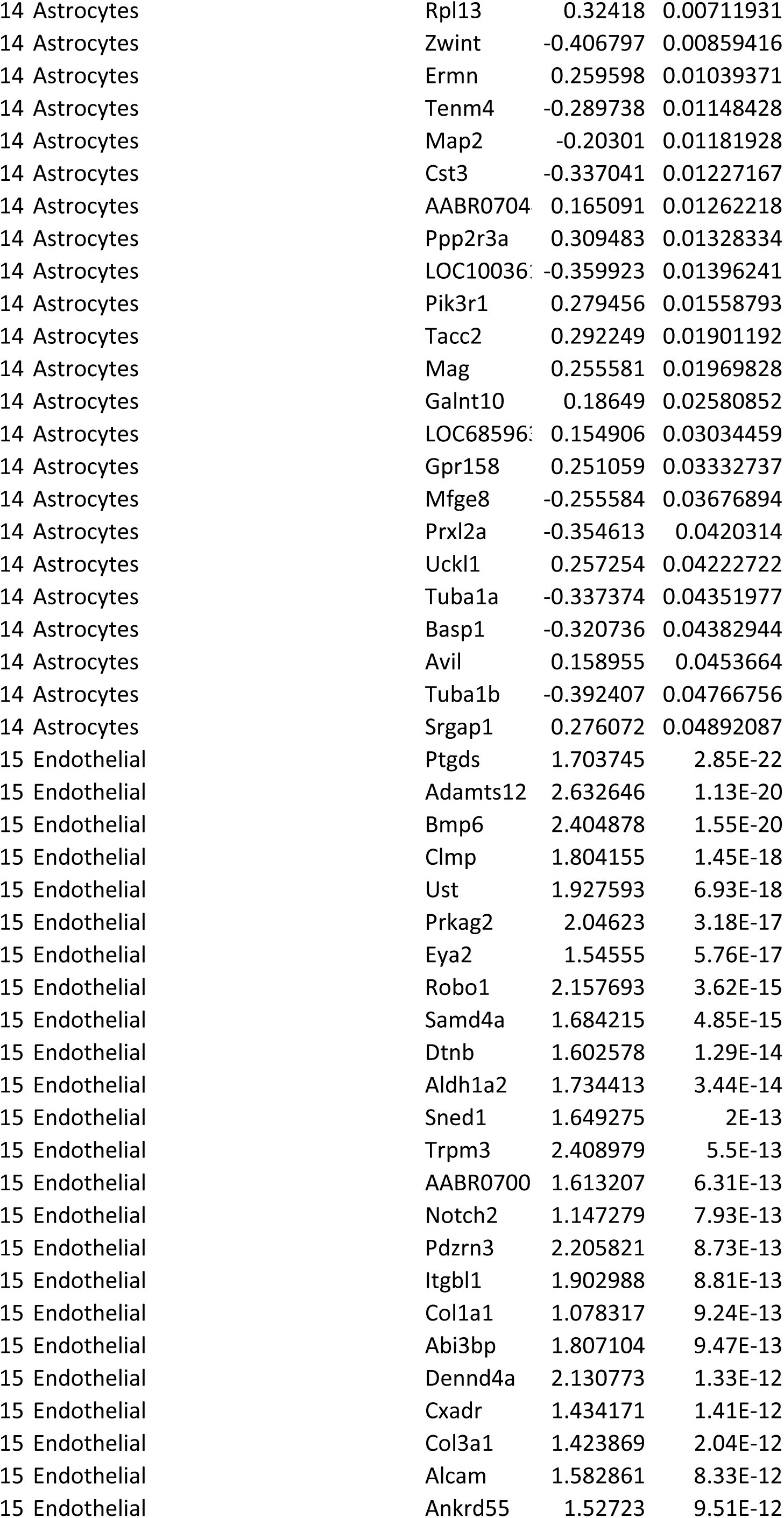
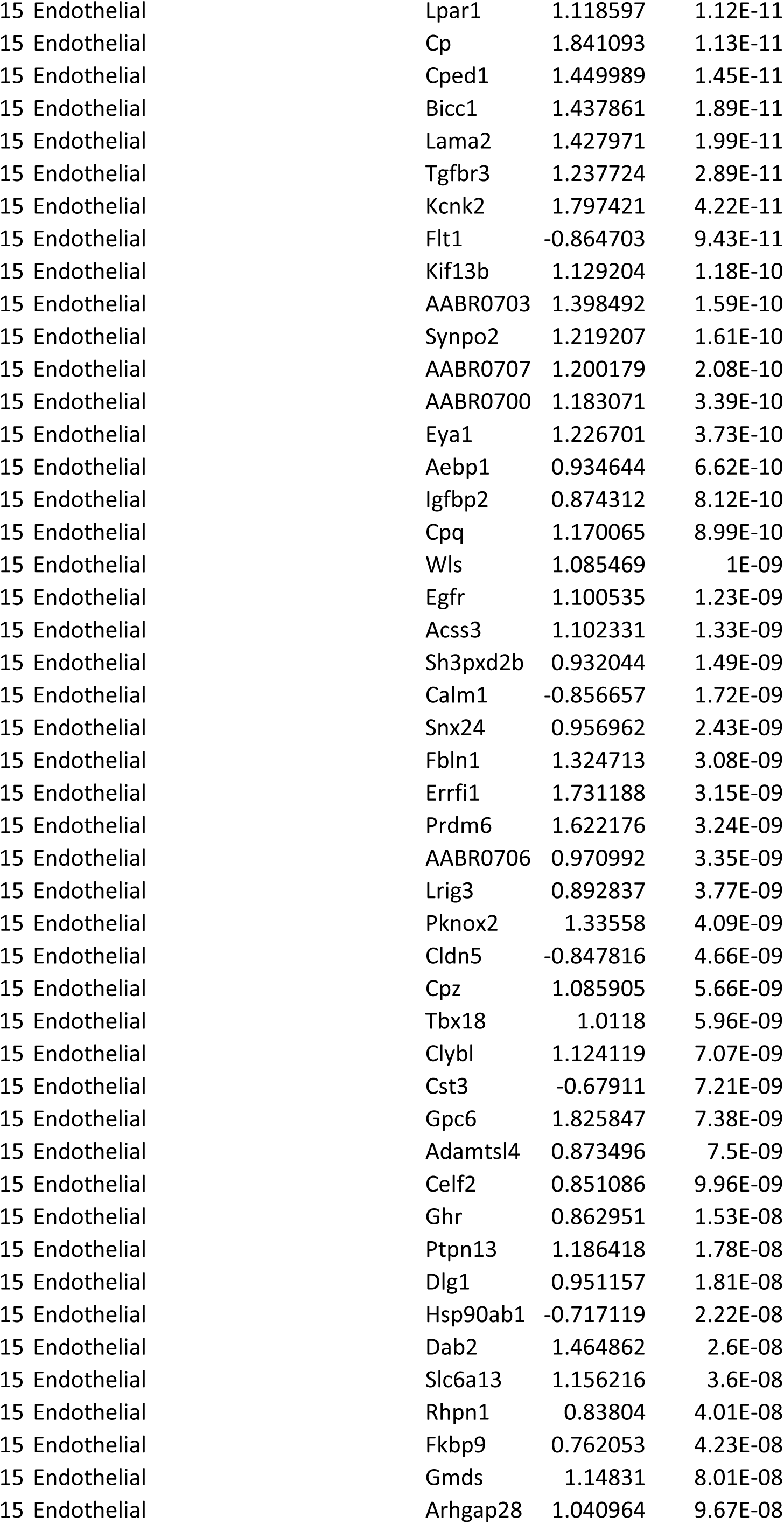
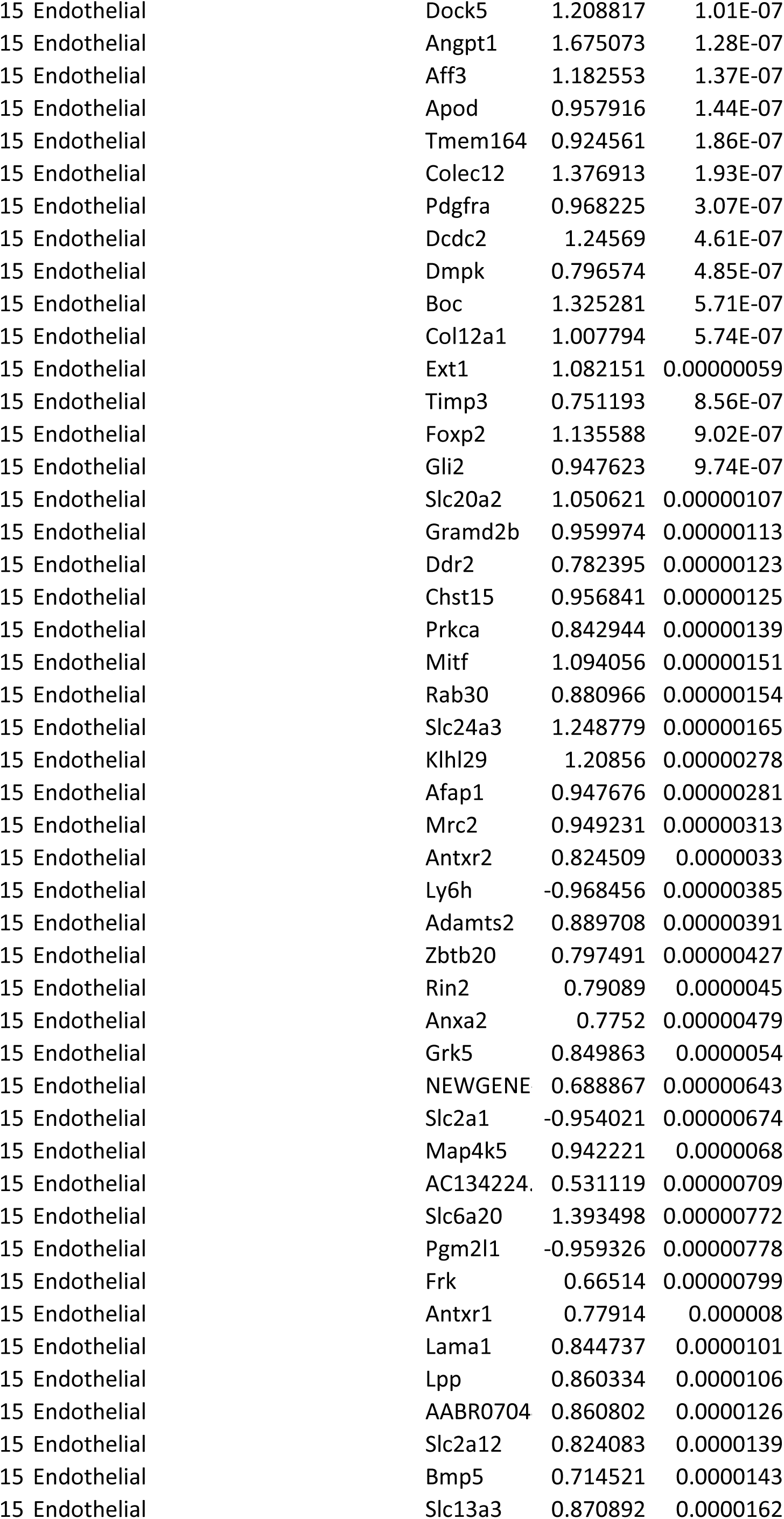
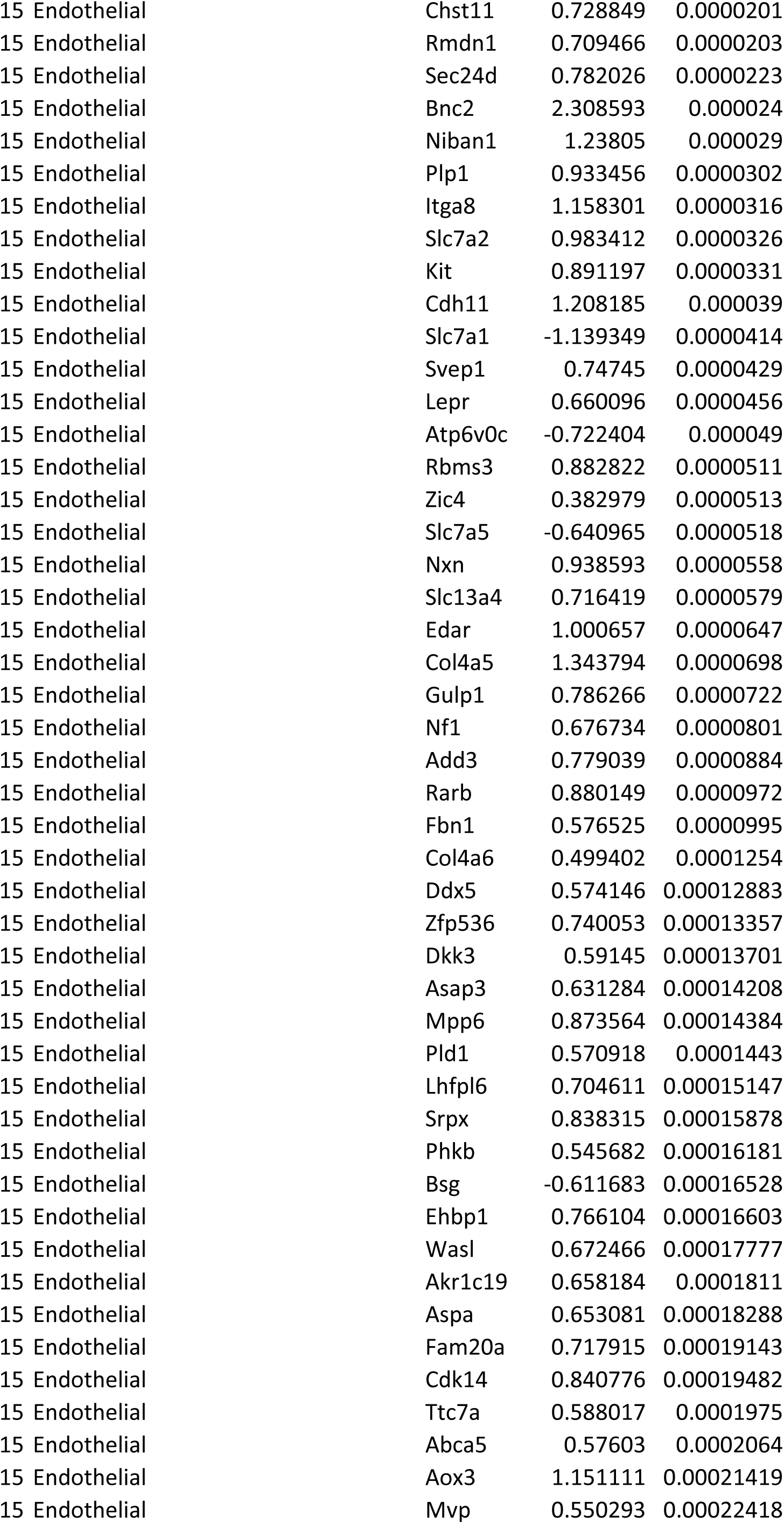
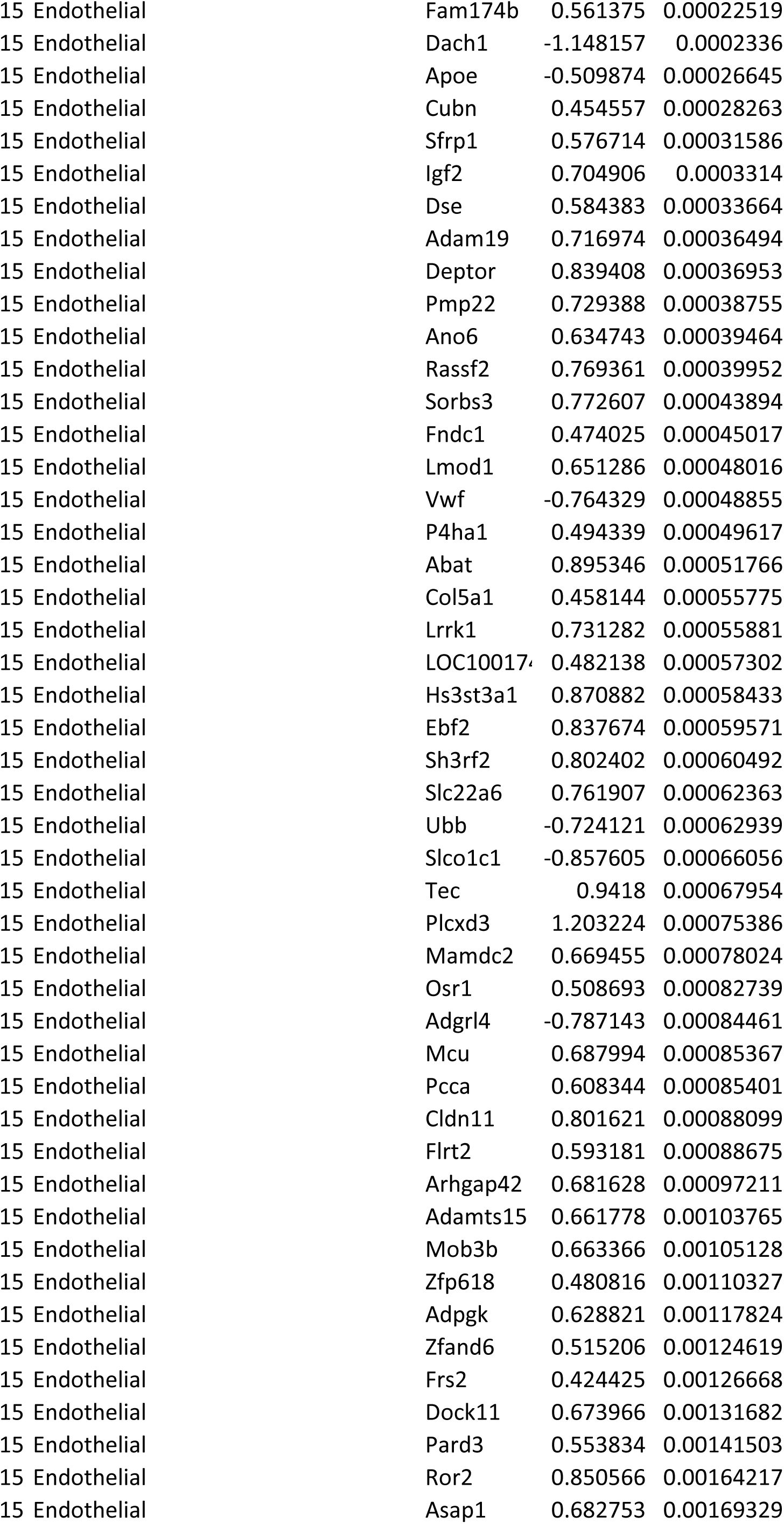
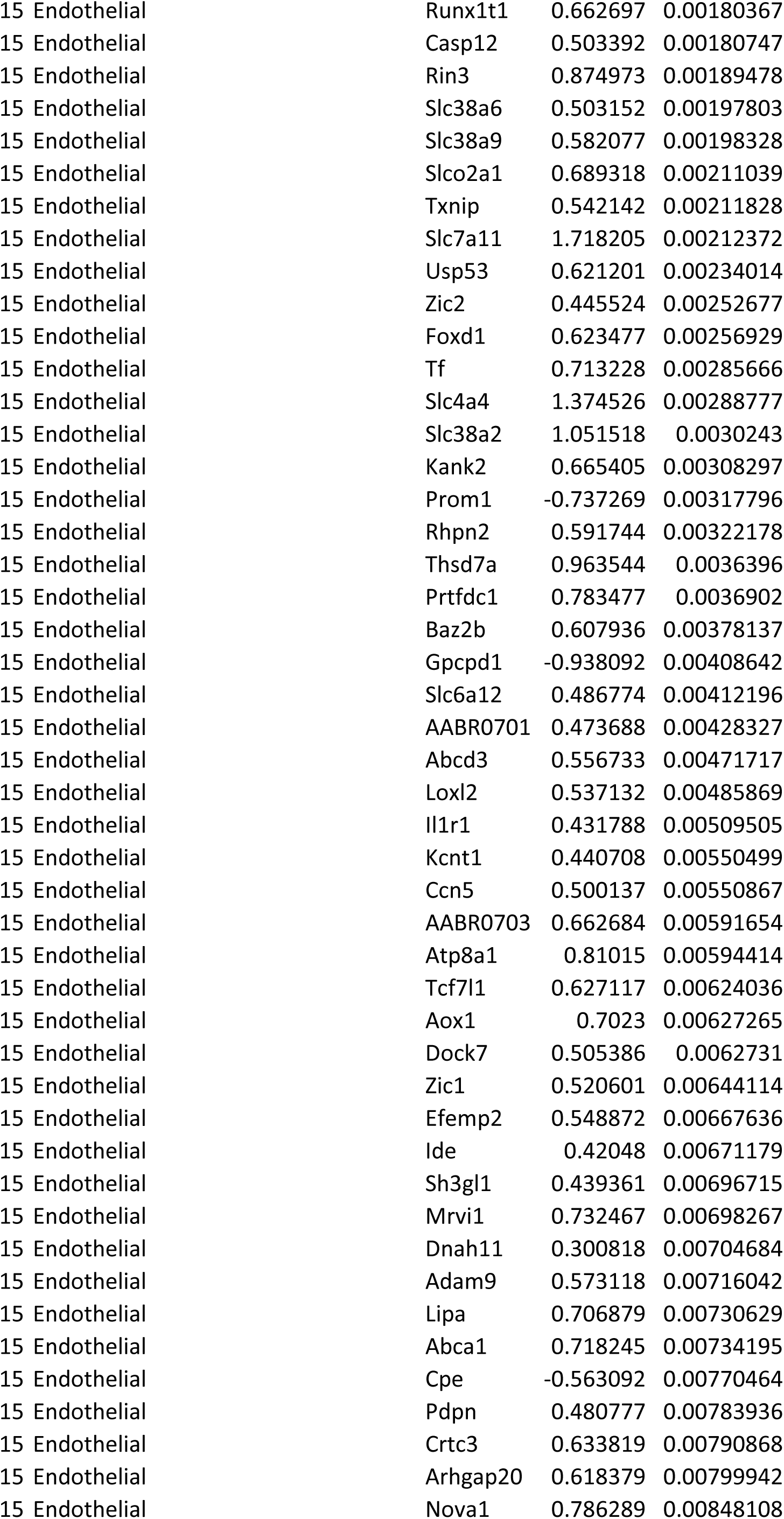
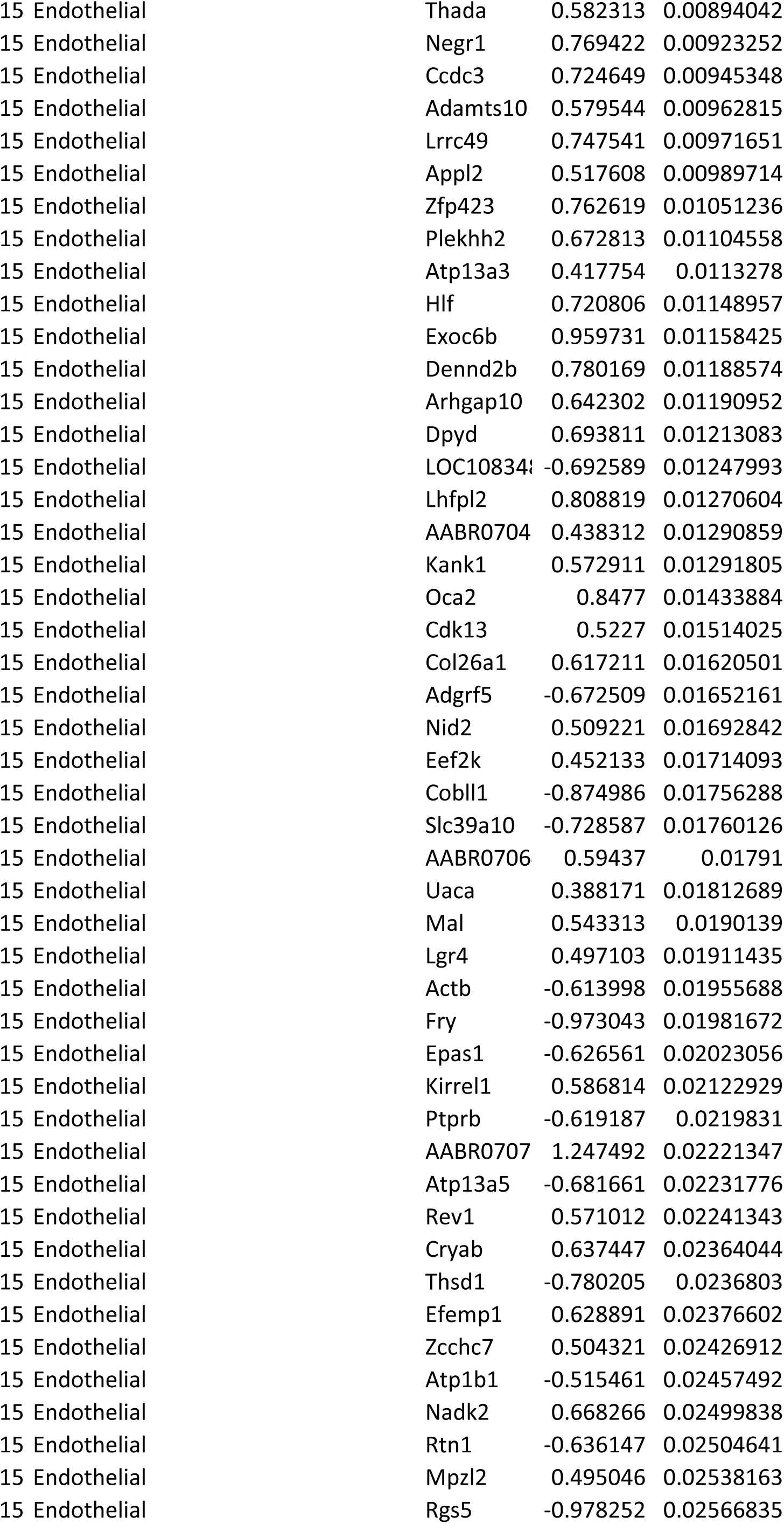
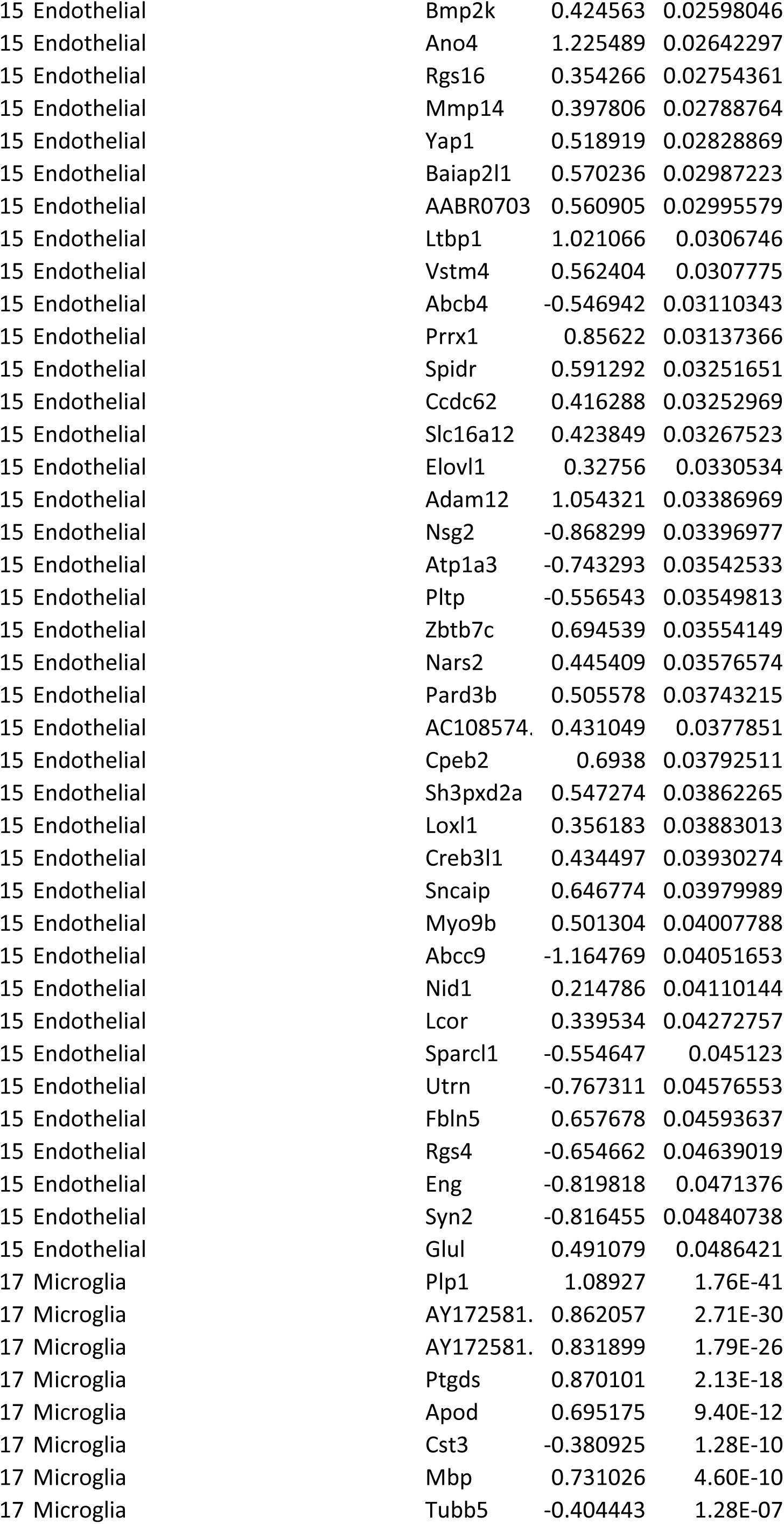
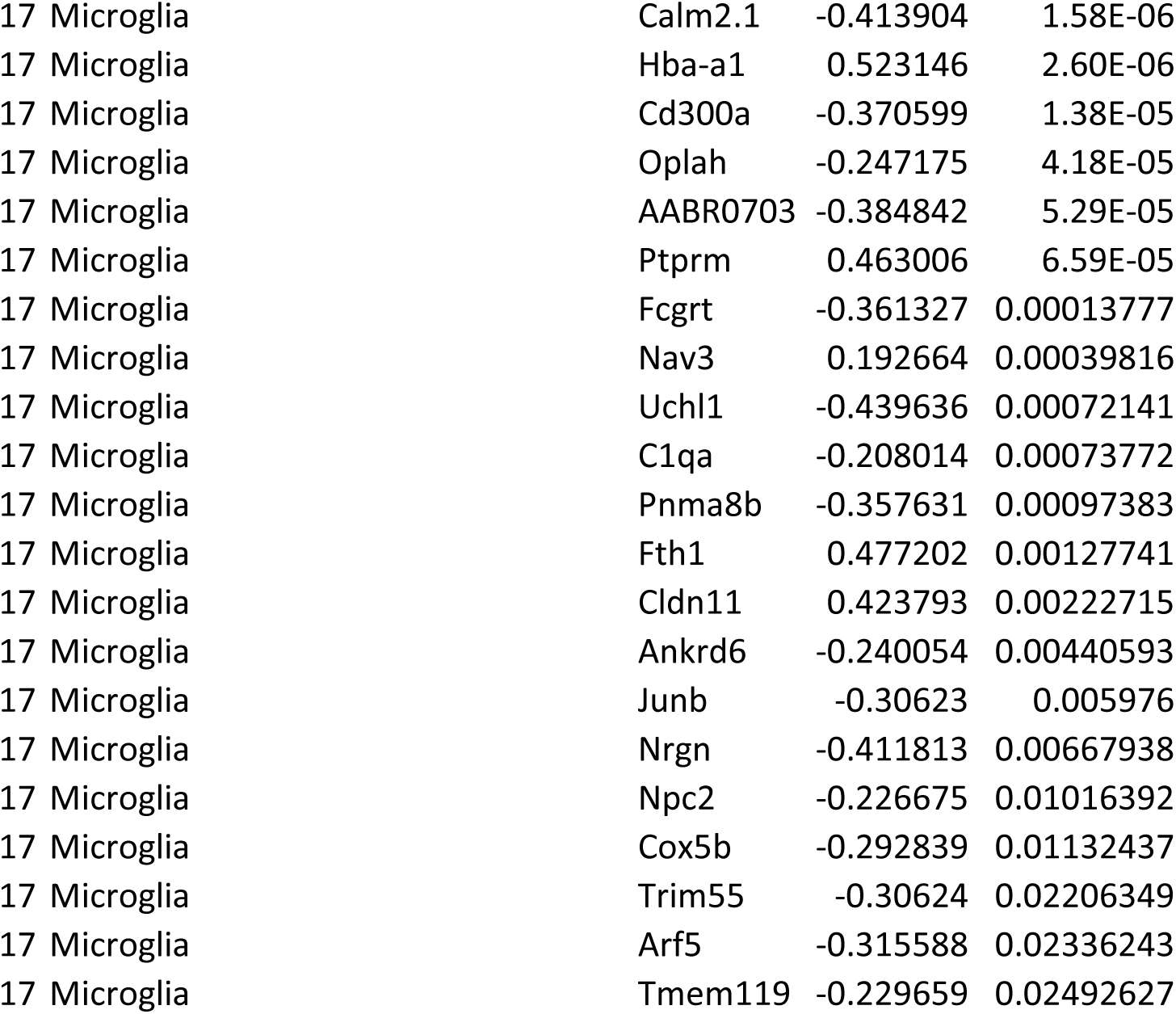
List of differentially expressed genes by cell cluster.

**Supplemental Table 3.**
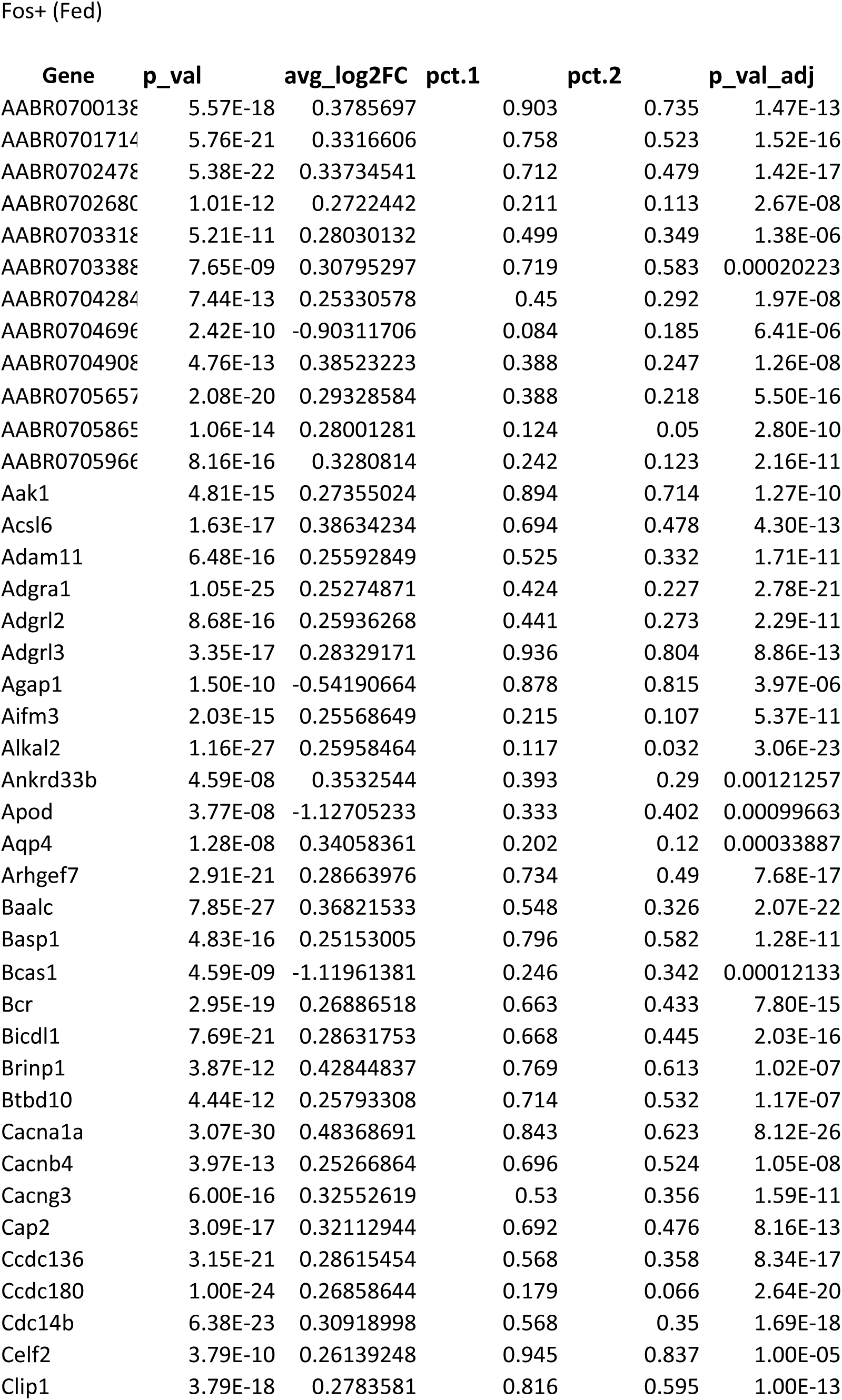
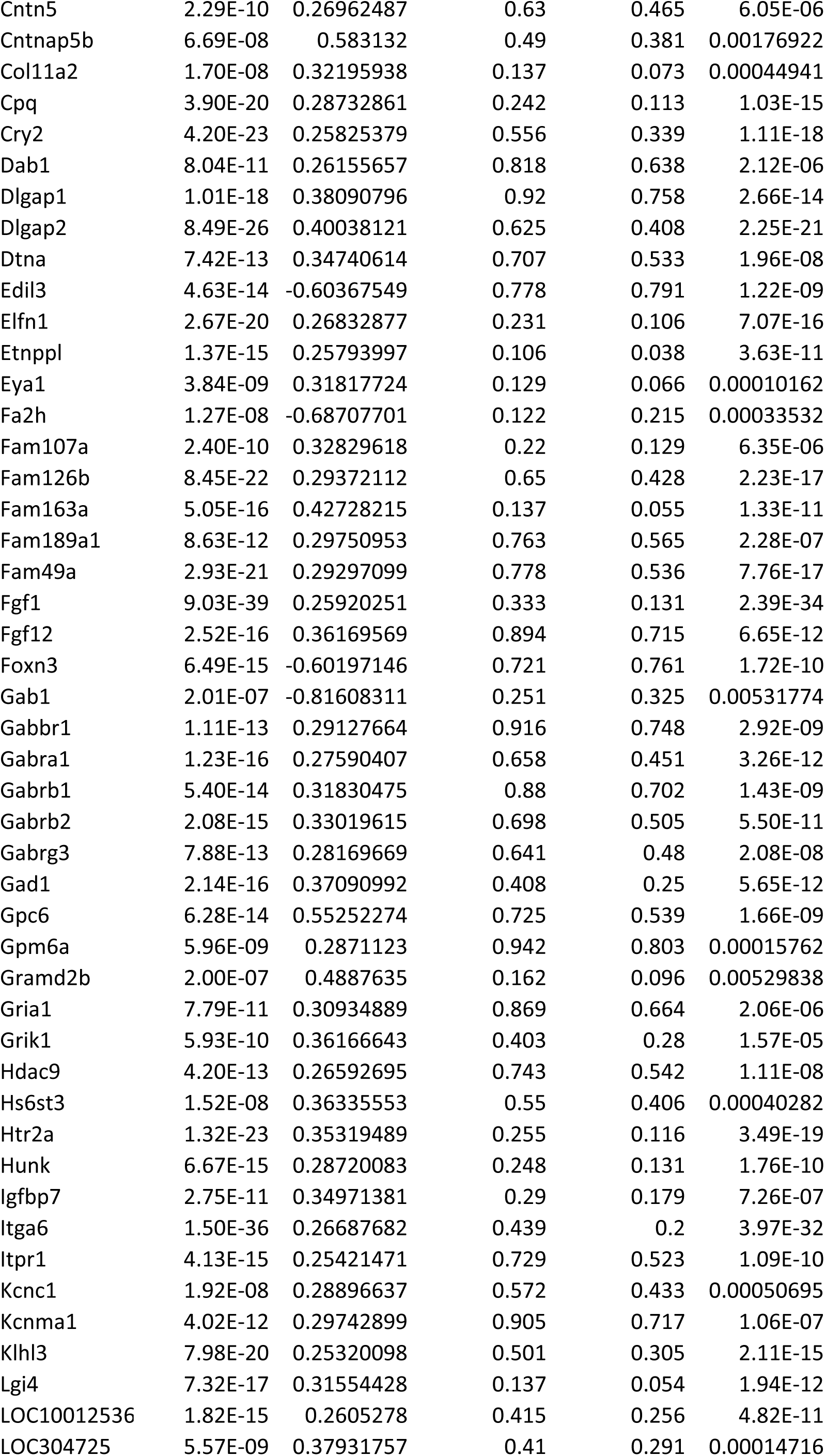
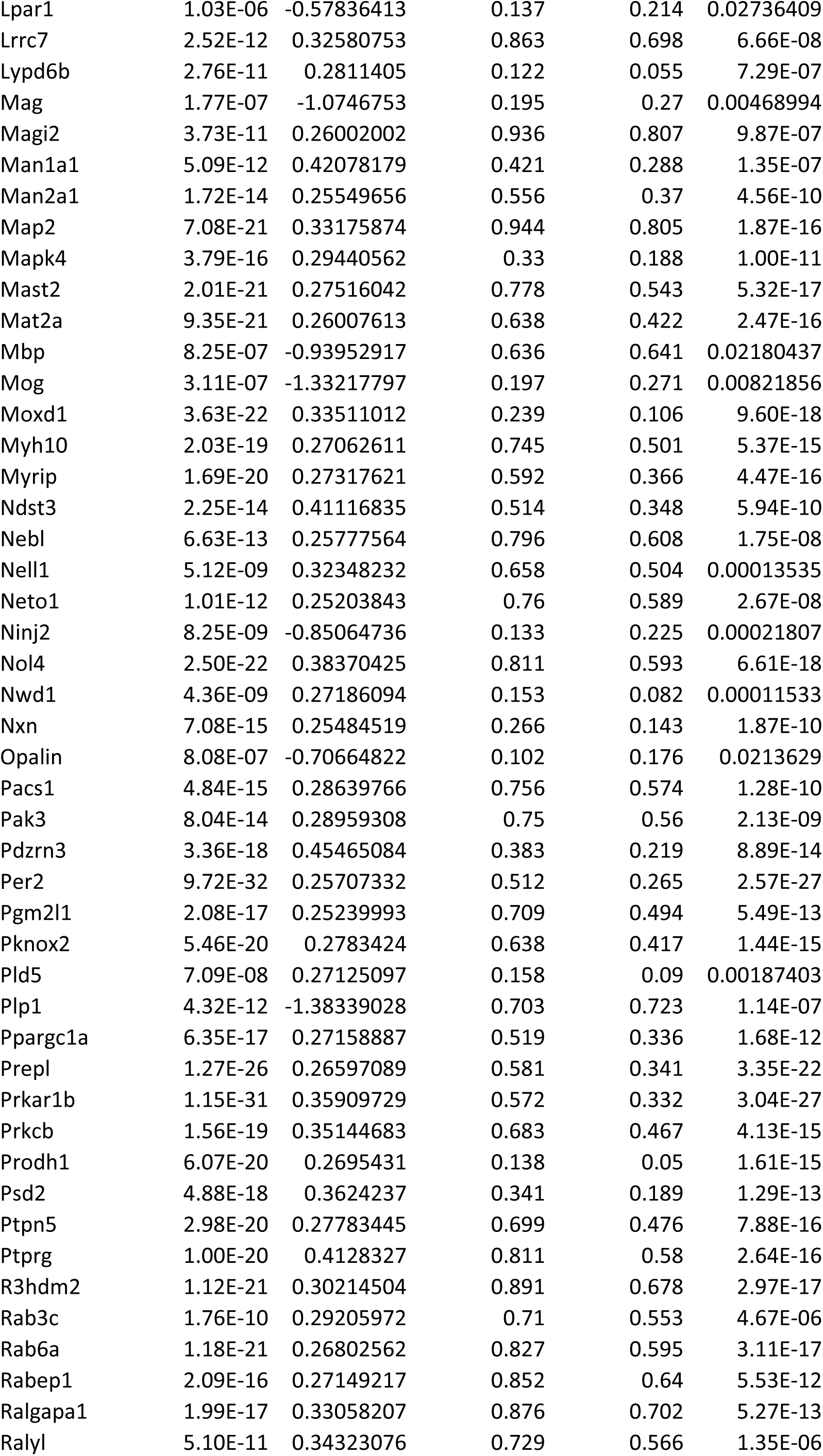
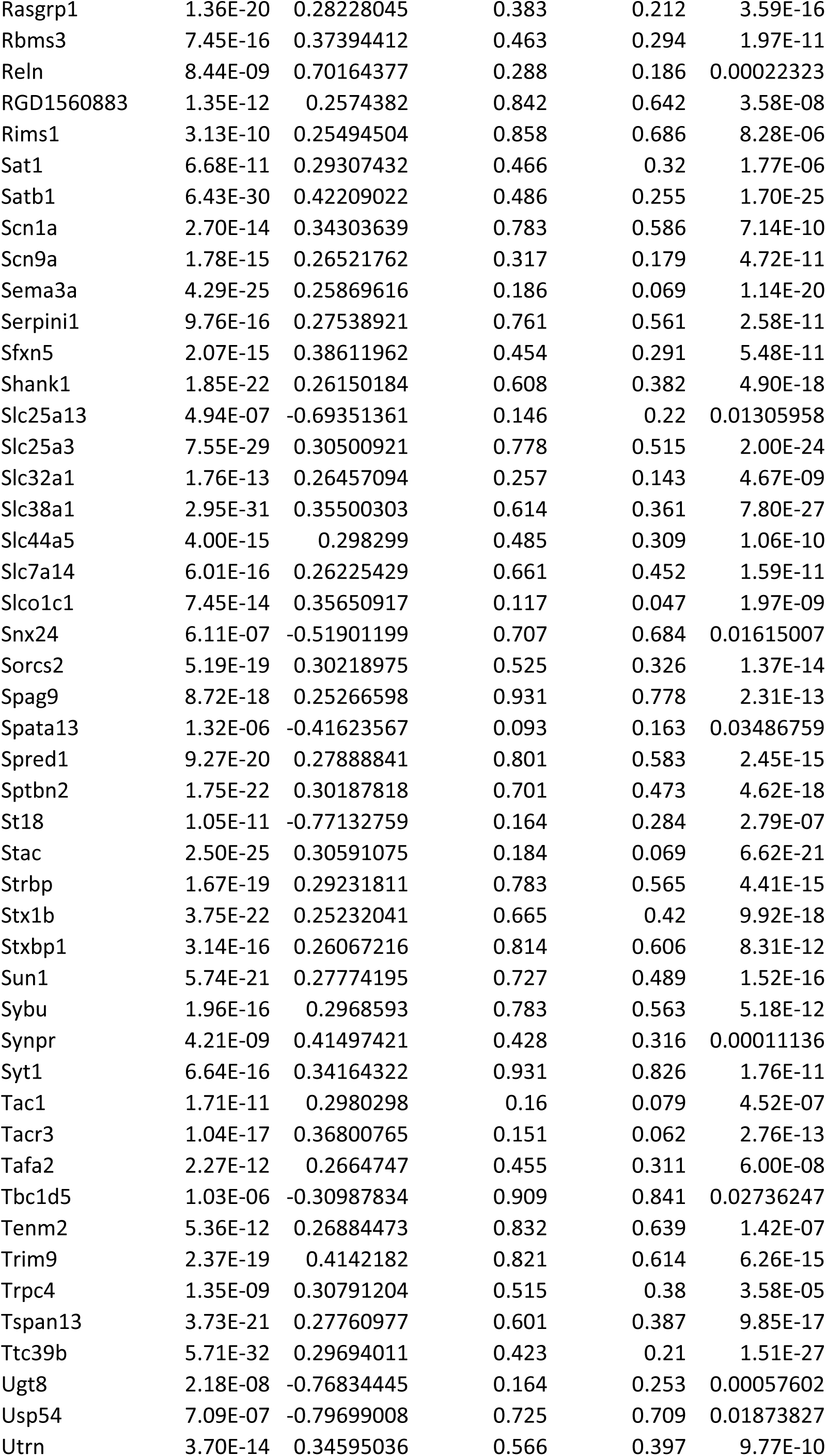
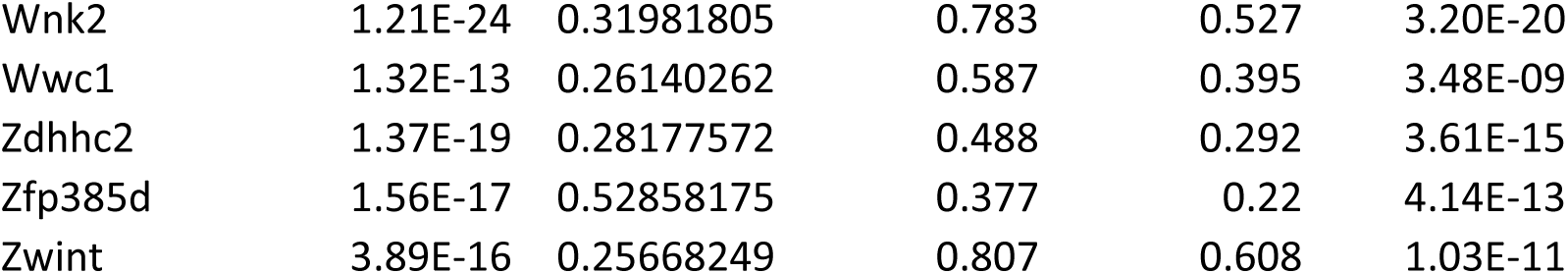
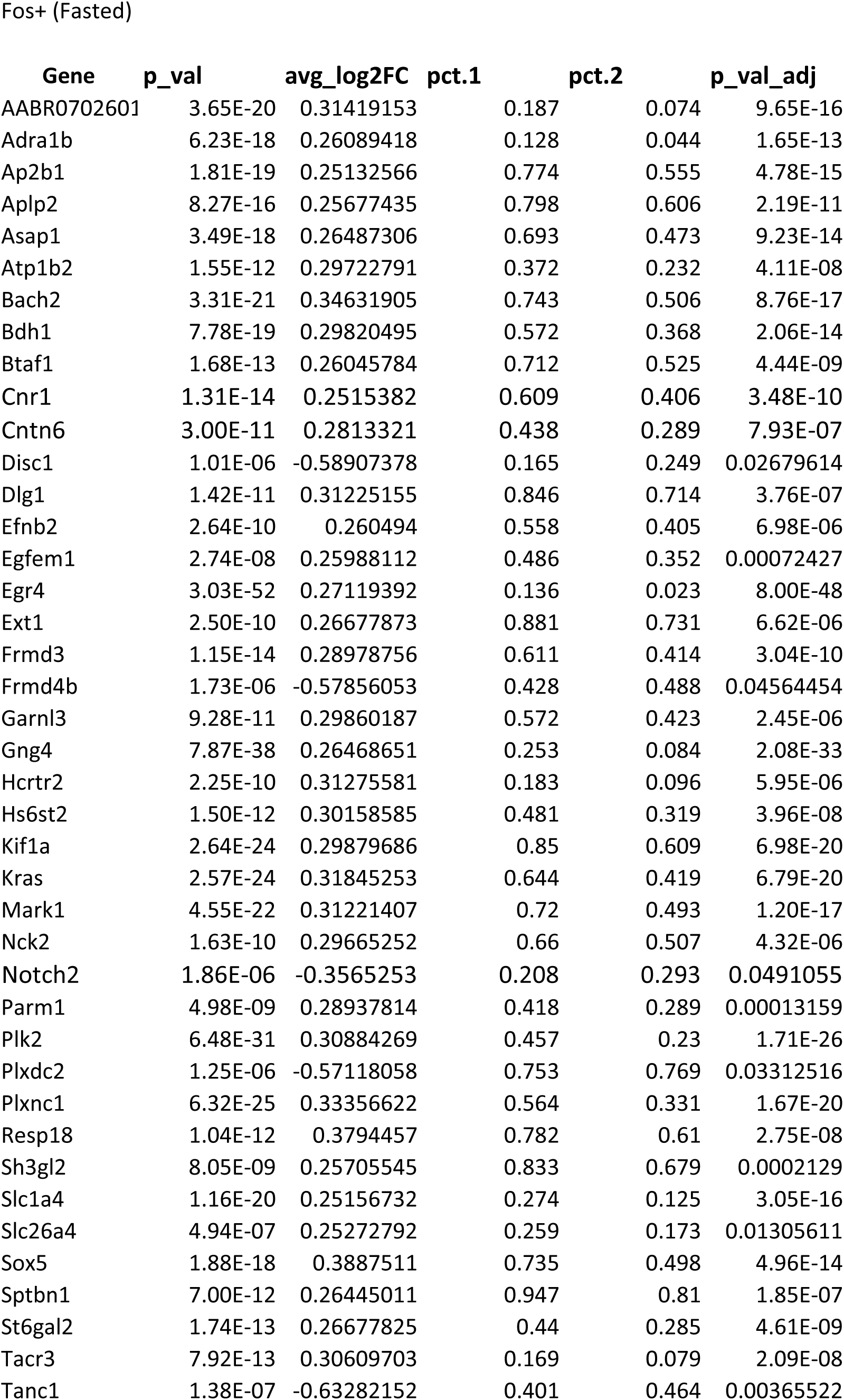
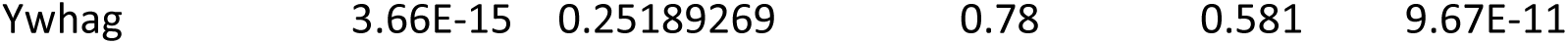
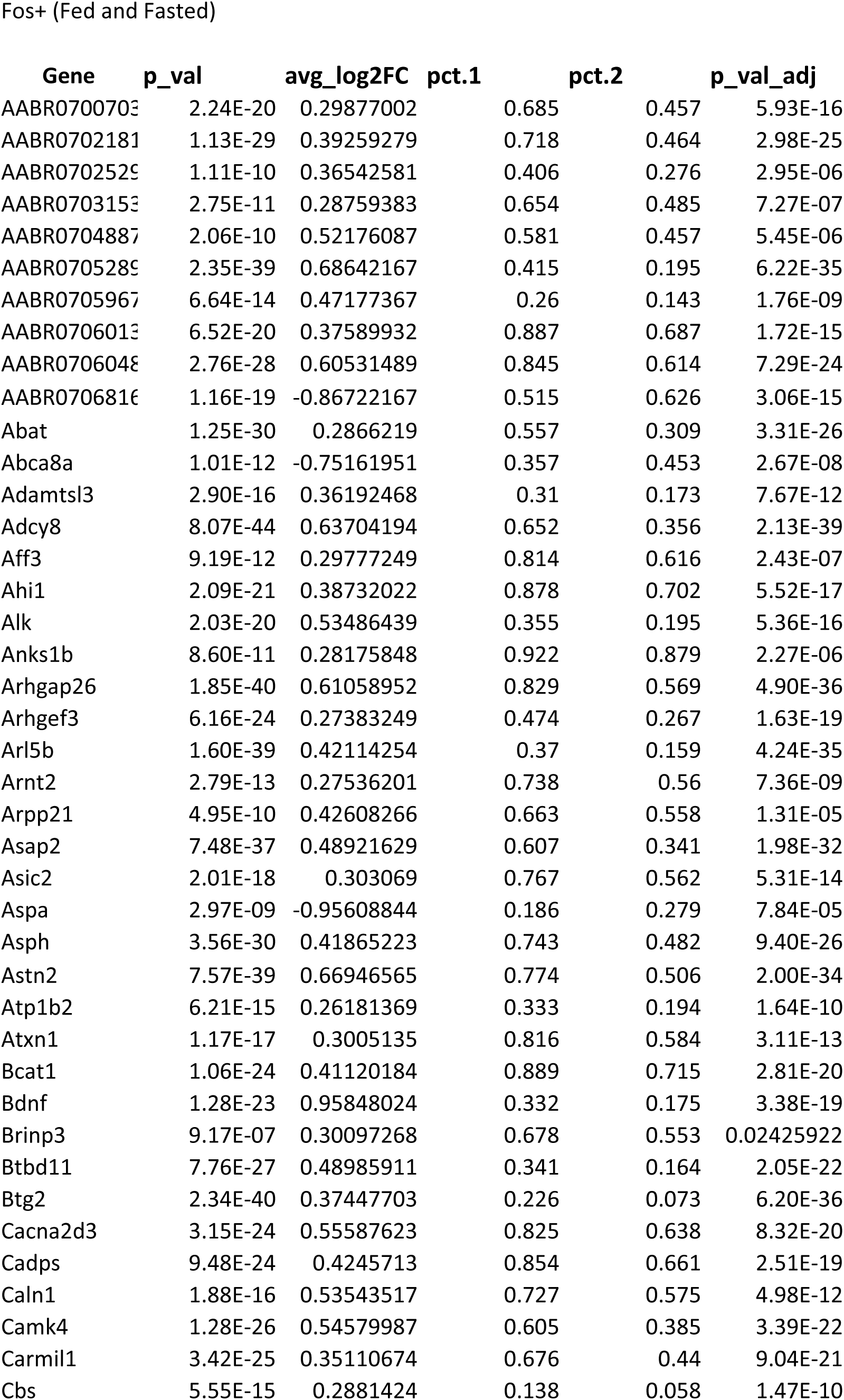
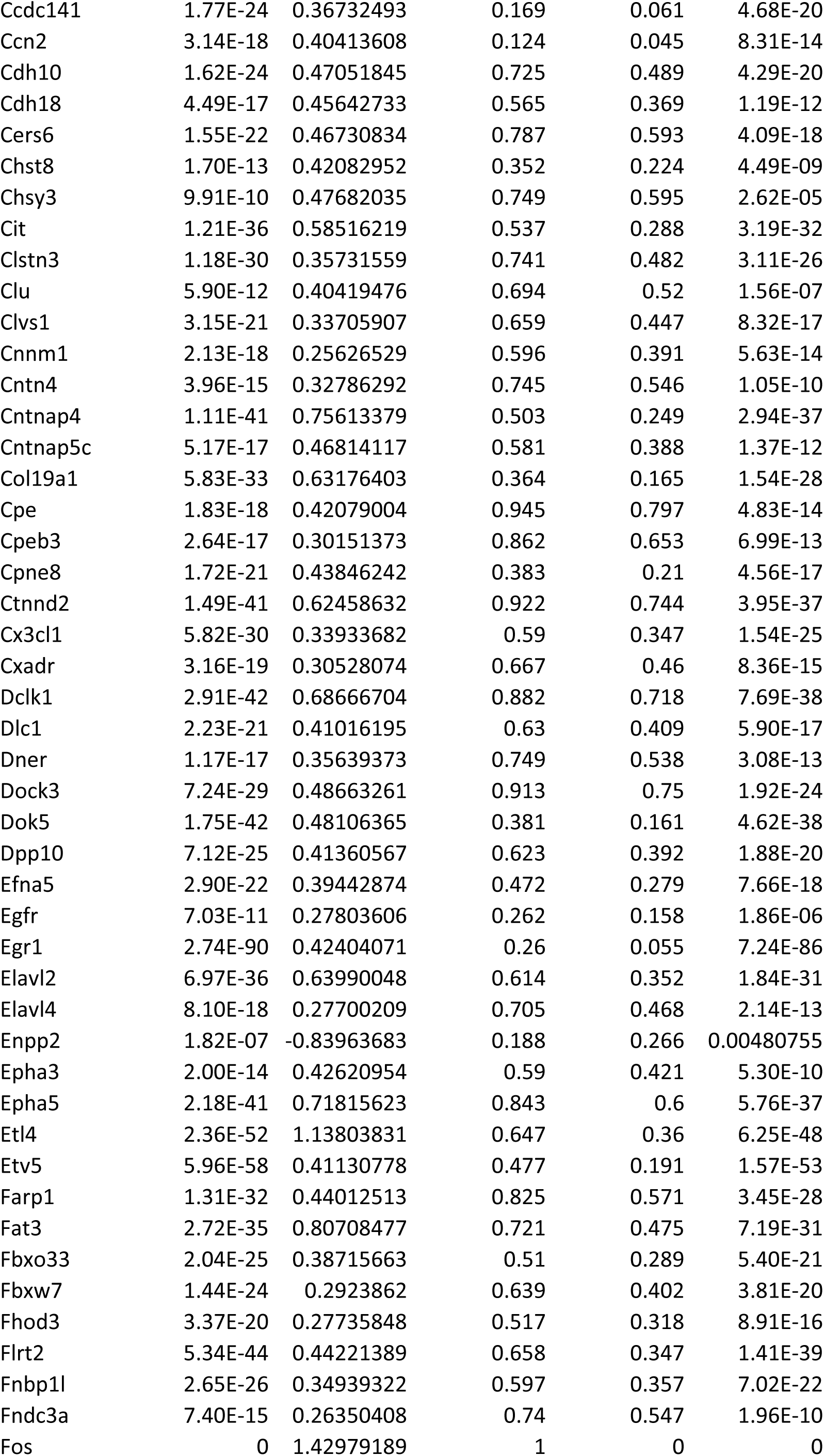
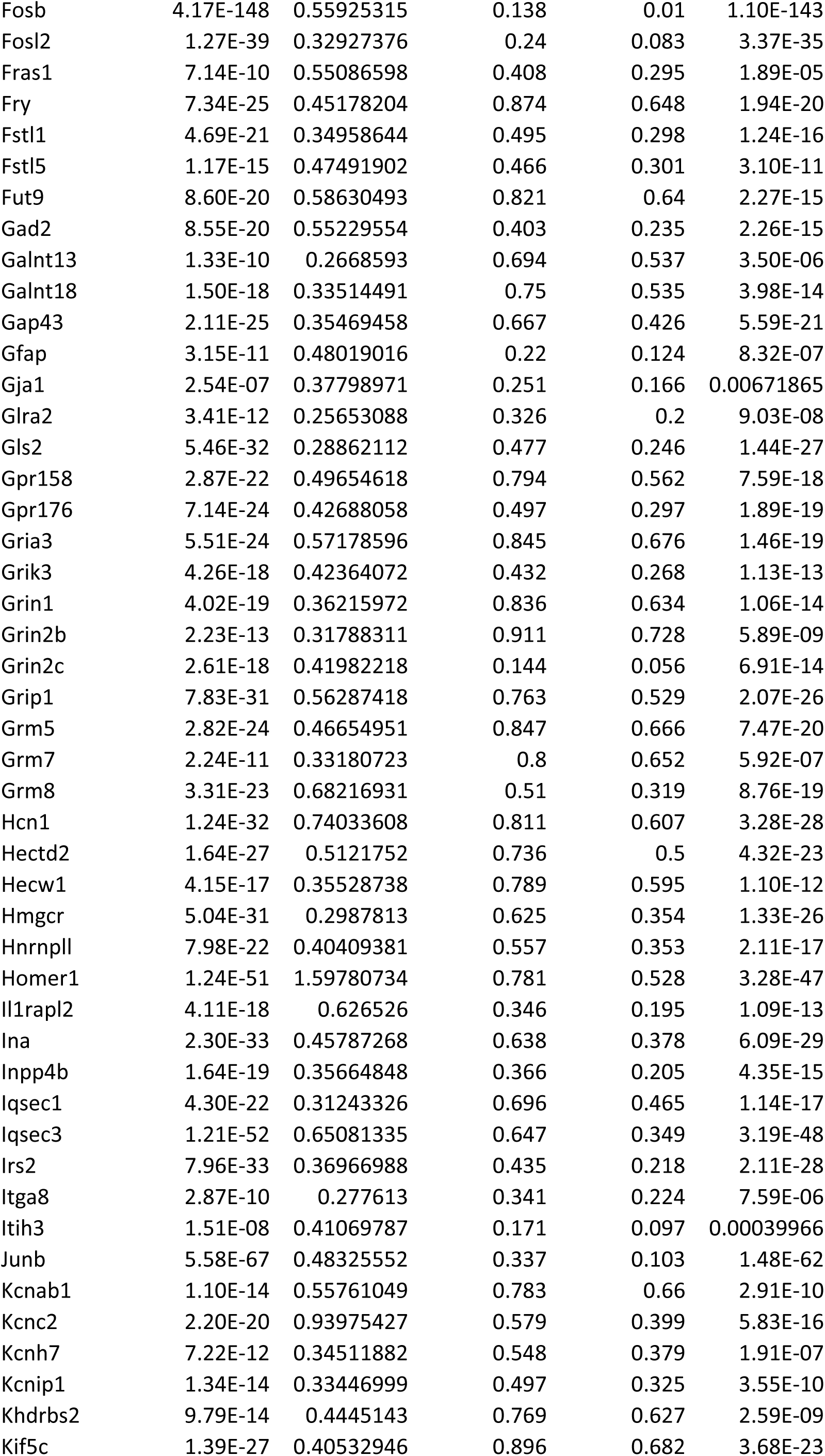
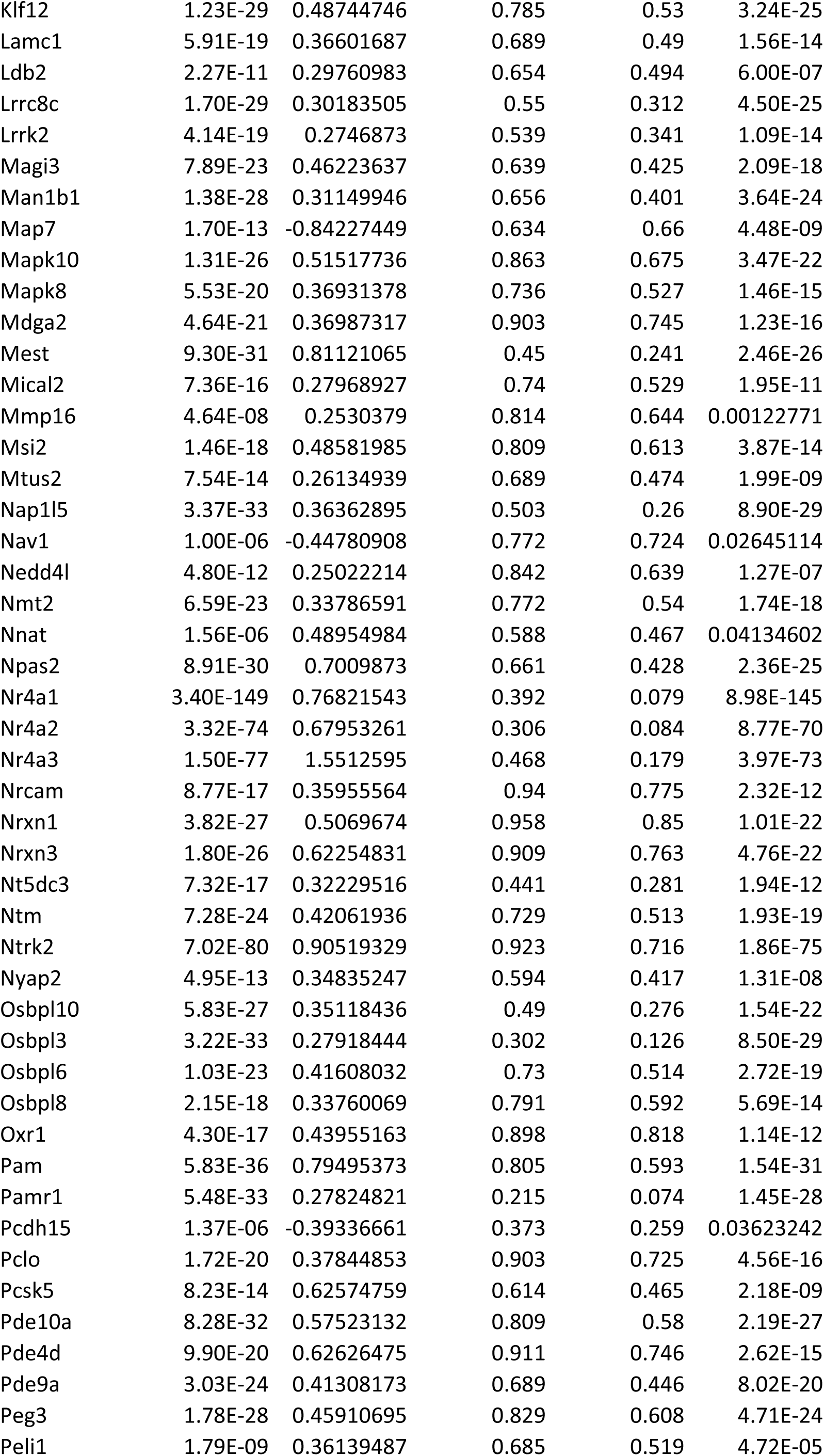
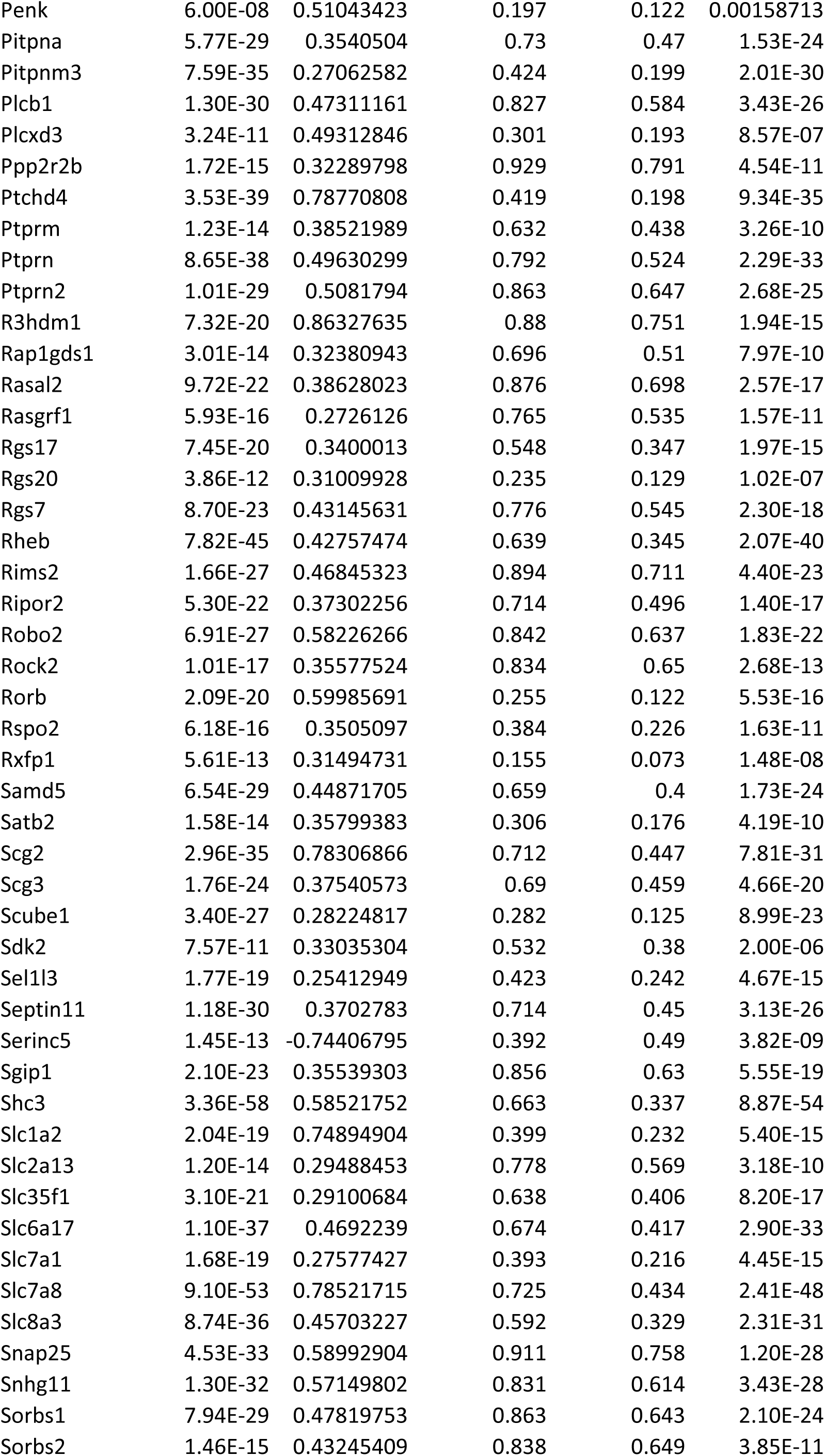
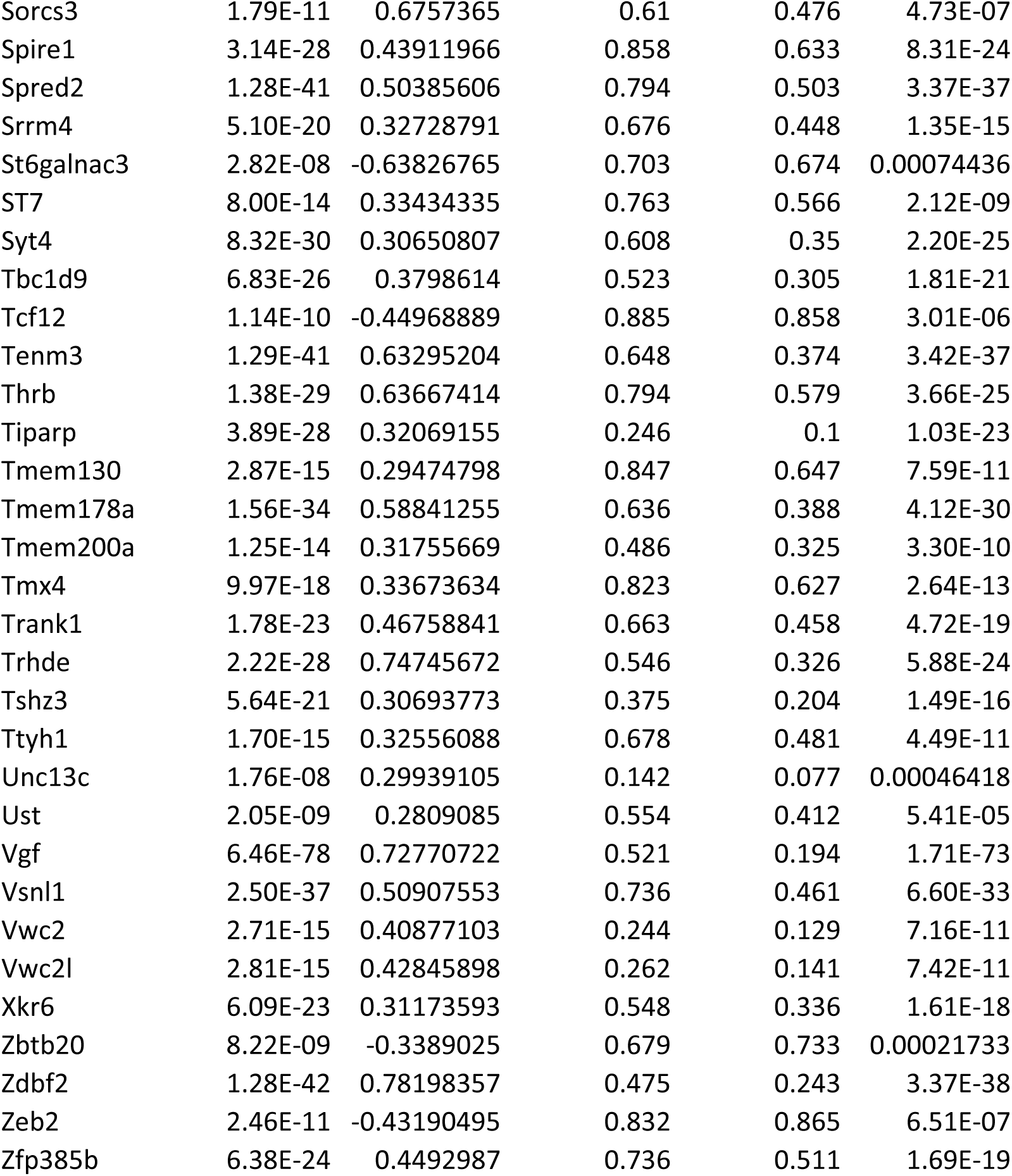
Genes expressed in Fos+ cells.

